# A synthetic gene circuit confers salt tolerance in plants

**DOI:** 10.64898/2026.04.29.721683

**Authors:** Tessema K. Kassaw, Kevin J. Morey, Alberto J. Donayre-Torres, Mauricio S. Antunes, Joshua T. Polito, Elizabeth A. H. Pilon-Smits, Susan Abernathy, Ryan Lonergan, Chad M. Ternes, June I. Medford

**Affiliations:** Department of Biology, Colorado State University, Fort Collins, CO 80523 USA

## Abstract

Soil salinity exerts a major constraint on global crop production, while seawater intrusion impacts coastal aquifers and surface waters. Using a blueprint from nature, we designed and implemented genetically encoded circuits that confer high salt tolerance to Arabidopsis and rice plants. Endodermal-like barriers, duplicated to the root epidermis and distally expanded to protect sensitive regions, provide tolerance to salinity levels comparable to seawater. Two additional genetic modules were added to reduce adverse effects of the epidermal barriers on ion uptake (aquaporin module) and to provide osmotic protection (osmolyte module). Arabidopsis and rice containing all three genetic modules can grow with continuous 600 mM NaCl treatment for extended days and reproduce. Transcriptomic analysis suggests that our rational engineering approach primes plants for salt tolerance, even without salt exposure, while detailed ionic analysis uncovers leakage suggesting means for improvement. Our results, duplicating suberin and the Casparian Strip in the epidermis while addressing adverse effects and osmotic stress demonstrate a new approach to enhancing salt tolerance while offering additional insights into plant responses to high salinity.

**One Sentence Summary:** A bio-inspired approach engineering epidermal barriers, symplast transporters and osmotic protection produce salt tolerance in Arabidopsis and rice.

## Main

Soil salinity, driven by factors such as irrigation and seawater intrusion^1^, affects approximately 10.7% of the Earth’s total land area^2^ and threatens crop production worldwide. Plant responses to salt stress entail activation of transporters, signaling components, transcription factors, ROS, and osmotic changes^3^. Efforts to increase plant salt tolerance have produced modest improvements^3^. Here we describe a synthetic biology approach based on anatomical and physiological features found in nature to engineer plants with high salinity tolerance.

Plant roots absorb water, minerals, and nutrients while excluding adverse molecules such as salt. Numerous aspects regulate root uptake, including an internal anatomical barrier, the endodermis, that controls water and ion entry into the plant’s vascular tissues^4^. In mature root regions, the exodermis, a cell layer internally/centripetally adjacent to the epidermis, and periderm provide some protection; however, roots absorb most of their water, ions, and nutrients through young, distal root regions and root hairs^5,6^, which lack such barriers. Arabidopsis roots lack an exodermal barrier and limit periderm development to the upper-most portion of the root of very mature plants^7^. In contrast, in some salt tolerant plants such as mangroves, the exodermis possesses both a Casparian strip (CS) and suberin lamellae^8^. While these structural elements extend distally in more salt tolerant species, they are not present in the root tip^8^ or root hairs. We hypothesized that duplicating root endodermal barriers to the epidermis, combined with distal expansion could replicate natural systems and provide salt tolerance (Fig. 1).

**Fig. 1.**
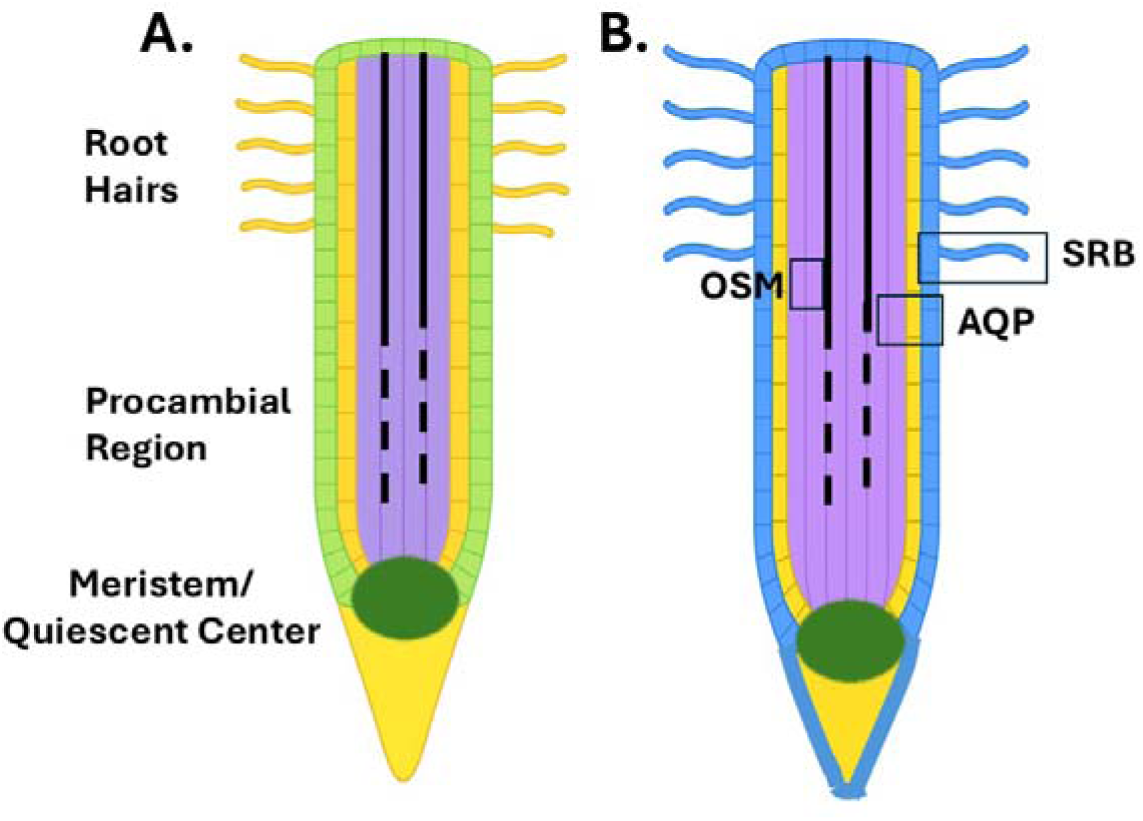
Arabidopsis root schematics of control a. and engineered b. **plants**. Elements found in both: root hairs, procambial region, meristem/quiescent center (green oval). Engineered plants (**b)** contain the Synthetic Root Barrier module, **SRB** (blue colored epidermis and root hairs) designed to expand and duplicate endogenous endodermal-like barriers, suberin and the Casparian Strip, to the root epidermis. Suberin is expressed in root hairs and distally to the root tip, protecting sensitive root elements. The aquaporin module, **AQP**, is designed to mitigate nutrient deficiencies that result from duplicated root barriers. The osmolyte module, **OSM**, is designed to address osmotic stress and retain water in the vascular tissues. The complete gene circuit, Synthetic Salt Tolerance, **SST**, combines all three modules. Rectangles represent the approximate target root areas for expression of the gene circuit.

### Duplicating barriers in the root epidermis

We designed a genetically encoded Synthetic Root Barrier (SRB) gene circuit/module to duplicate internal endodermal barriers to the root epidermis. We first experimentally validated promoters for root cell-specific expression (Supplementary Table 1), and placed genes needed for the CS (*AtMYB36*) and suberin synthesis (*AtMYB41*) under control of the epidermal *AtABCG37/PDR9* promoter^9^. We duplicated the CS in the root epidermis using MYB36, a transcription factor known to direct its formation^1^^5,16^. *MYB36* was placed under the control of the epidermal *ABCG37* promoter^9^ with a 5’ leader sequence (omega) to enhance translation. The epidermal expression of MYB36 is sufficient to produce an additional CS in the root epidermis (Fig. 2A).

**Fig. 2.**
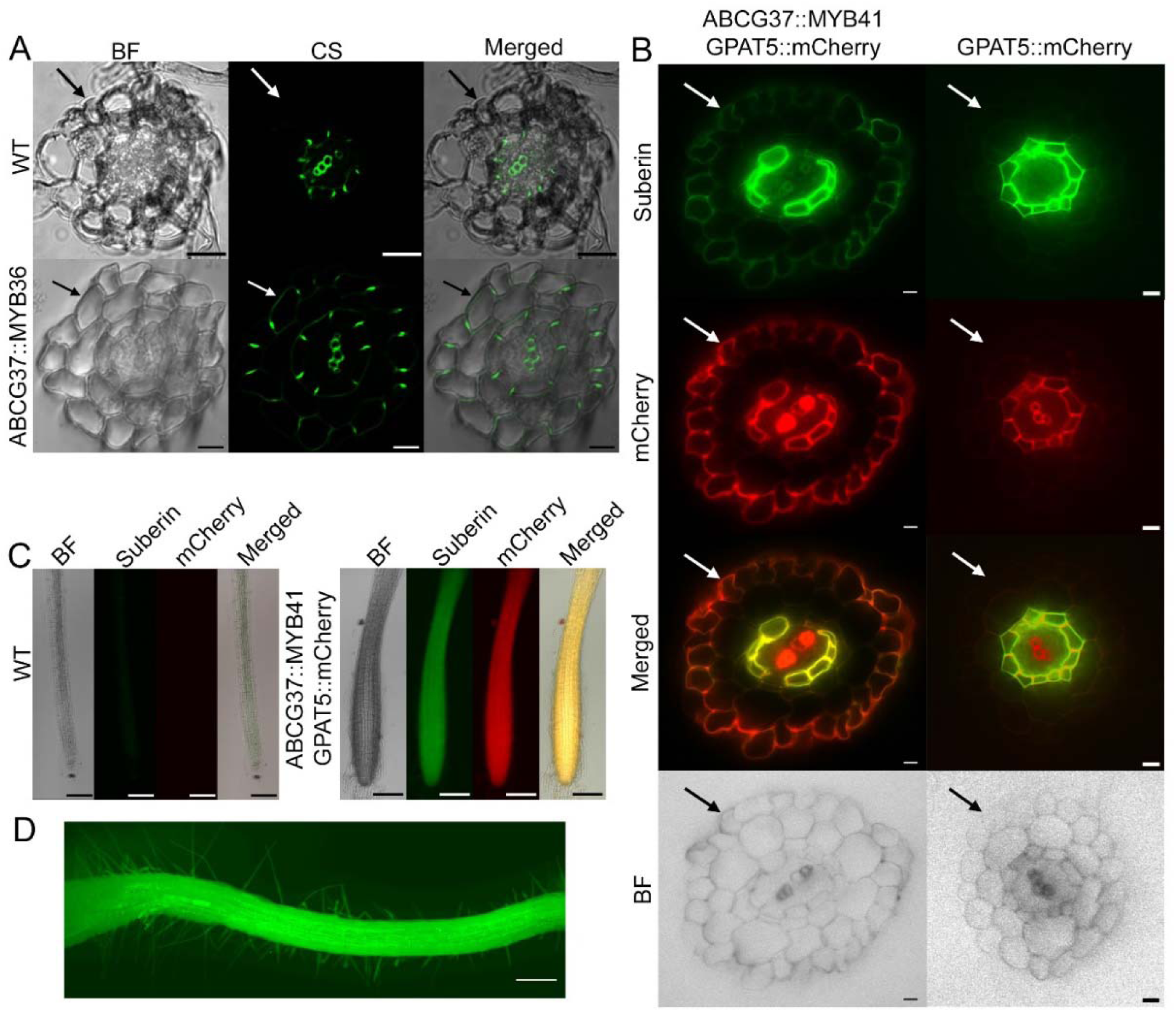
Epidermal production of Casparian strips and suberin in Arabidopsis roots. **a.** Root cross-sections of Columbia Col-0, wildtype (WT) and transgenic plants expressing *ABCG37::MYB36*, sections stained with berberine showing epidermal Casparian strip production. **b.** Root cross-sections showing suberin production in the epidermis in plants expressing (*ABCG37::MYB41S251D*). *GPAT5::mCherry*, is used as a marker for suberin localization; green fluorescence from suberin stained with fluorol yellow. **c.** Whole mounts of roots showing epidermal suberin (fluorol yellow stain) and GPAT5::mCherry expression extends to the root tip. **d**. Longitudinal view, whole mount showing suberin staining (fluorol yellow, green fluorescence) in roots and root hairs of plants transformed with *ABCG37::MYB41^S251D^* in mature roots and root hairs. Scale bars = 10 µm, (**a-b)**, 100 µm, (**c)**, 10 mm, (**d),** BF, Brightfield, CS, Casparian strip.

Previous work indicates that *AtMYB41* constitutive expression produces ectopic suberin deposition in leaves, but not in roots, other than the endodermis^10^. MYB41 activity depends upon phosphorylation of serine 251 by a MAP kinase^11^. We hypothesized that the absence of ectopic suberin in roots results from a lack of MYB41 phosphorylation in non-endodermal cells. Some amino acid substitutions can mimic phosphorylation^12,13^; hence MYB41 was engineered to be orthogonal to MAP kinase controls through a phospho-mimic: serine 251 was mutated to aspartate (MYB41^S251D^) and placed under control of the *ABCG37* promoter. Plants with *ABCG37::MYB41^S251D^*accumulate suberin in the root epidermis, distally to the root tip and including root hairs (Fig. 2) (closeup, Extended Data Fig.1). All transgenes described herein (Arabidopsis and rice) are refactored and expression-verified with end-point RT-PCR (Supplementary data 1, 2, Extended Data Figs. 5&6). To facilitate analysis, *GPAT5::mCherry*, a marker for suberin biosynthesis^14^, was co-transformed with *ABCG37::MYB41^S251D^*. *ABCG37::MYB41^S251D^* transgenic lines express the *GPAT5::mCherry* suberin biosynthesis marker and accumulate suberin in the root epidermis distally, including the root tip (Fig. 2B-D). These genetic components direct production of epidermal suberin and epidermal Casparian strip and constitute our SRB module.

Suberin is a complex biopolymer of fatty acid derivatives, phenolics and glycerol, with variable composition and three-dimensional deposition^17,18^. We conducted a preliminary analysis of suberin content in plants engineered with the SRB module and plants with the complete gene circuit for Synthetic Salt Tolerance (SST2-1). A notable and significant difference is found in both engineered lines with the levels of the very long chain fatty alcohol, stearyl alcohol (SRB2-1, 6.68-fold and SST2-1, 10.47-fold increase) while its acid derivative (stearic acid) increased 2.55-fold and 5.34-fold, respectively (Table 1, Supplementary Table 3&4, Supplementary Data 3). As the major aliphatic monomer of suberin, stearyl alcohol is critical to forming root apoplastic barriers^17^, hence our results suggest that our synthetic epidermal barrier may provide a functional a similar apoplastic barrier.

**Table 1.** Suberin composition in plants with epidermal suberin (SRB module) or the full SST circuit treated with water or 200 mM NaCl. Shaded cells indicate significant differences relative to Col-0 controls (P <0.05).

|  | Compound | Fold Change (vs Col-0) |  |  |  |
| --- | --- | --- | --- | --- | --- |
|  |  | Water |  | 200 mM NaCl |  |
|  |  | SST2-1 | SRB2-1 | SST2-1 | SRB2-1 |
| Waxes | Stearyl alcohol | 10.47 | 6.68 | 1.24 | 1.00 |
|  | Palmitic acid | 0.91 | 0.52 | 0.81 | 0.75 |
|  | Phthalic acid-like | 0.94 | 0.44 | 0.46 | 0.14 |
|  | Docosanoic acid, 2,3-propyl ester | 0.92 | 0.82 | 0.67 | 1.27 |

Natural root endodermal suberin is known to be regulated by environmental feedback, with NaCl stimulating its accumulation^19^. Control plants (Col-0) treated with 200 mM NaCl had 13-fold higher stearyl alcohol content and 2.6-fold higher phthalic acid-like waxes (Suberin Fold change tap water vs salt treatment, Supplement Table 4). In contrast, both SRB2-1 and SST2-1 simply doubled their stearyl alcohol content. One possibility for a lower stearyl alcohol content in engineered lines with salt (13-fold vs 2-fold), is that the synthetic epidermal suberin provides a barrier to salt.

We next tested our SRB plants for salt tolerance under high salt conditions (600 mM NaCl, 12 days total salinization treatment). Under low light, SRB plants showed salt tolerance; however, these plants also displayed variable degrees of stunting and chlorosis (Supplementary Fig. 1A-C). Because endodermal barriers play a critical role in regulating water and ion uptake^20^, epidermal barrier duplication could have a detrimental impact on plant growth. When plants with the synthetic epidermal barriers are grown in high-light, in addition to showing stunting, SRB plants show a strong chlorotic phenotype (Fig. 3, center), suggesting that plants are deficient in nutrient ions (e.g., iron, magnesium) and require an additional rational engineering component to overcome these effects.

**Fig. 3.**
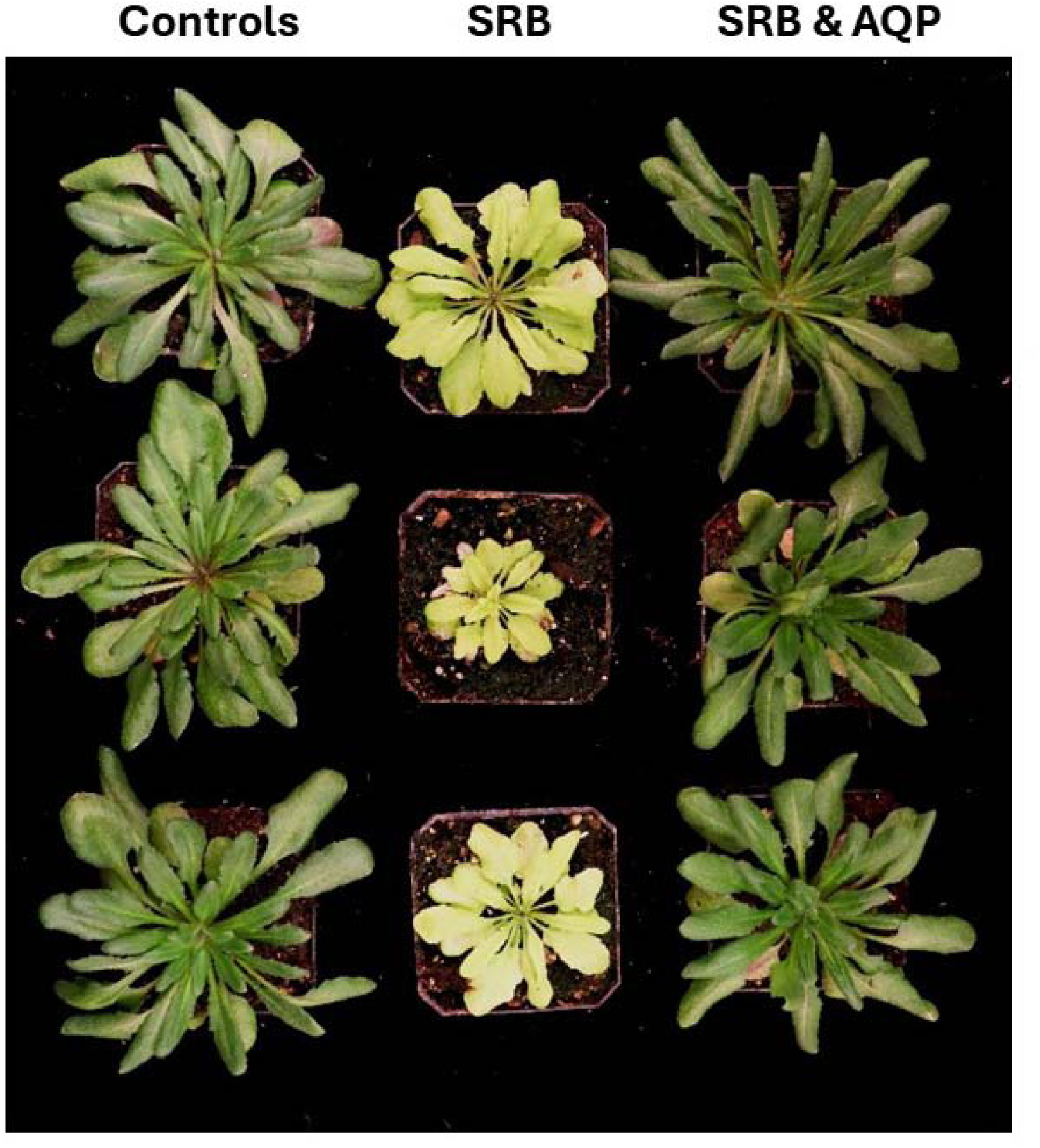
Characteristics of plants containing the synthetic root barriers (SRB) and aquaporin (AQP) modules grown under high light conditions. *Left*, Col-0 controls; Center, SRB plants; *Right*, plants with both the SRB and AQP modules. Plants containing the SRB module are chlorotic and stunted (center) suggesting a deficiency in nutrient ions. Co-expression of the AQP module with the SRB module corrects these phenotypes (*right*).

### Aquaporins mitigate issues from duplicated root barriers

Plants transport ions and water through natural endodermal barriers via the symplast, a highly regulated membrane system. Plant aquaporins are primarily found in plasma and tonoplast membranes and facilitate transport of water, solutes, and gases, as well as sequestering salt in the vacuole^21,22^. We reasoned that aquaporins from mangroves, while relatively uncharacterized, would likely have superior functionality to facilitate symplastic water flow, and, salt sequestration, while assisting needed ion uptake. Therefore, we designed a genetically encoded Aquaporin module (AQP) comprised of mangrove Plasma membrane Intrinsic Proteins (PIPs) and Tonoplast Intrinsic Proteins (TIPs). Aquaporins can be post-translationally regulated^23^ and information about the regulation of mangrove aquaporins expressed in Arabidopsis roots is lacking. Hence, we first performed an initial functional screen for the stability of seven refactored PIPs and four refactored TIPs originally from mangroves and the halophyte, *Thellungiella salsuginea* fused to fluorescent reporters (mCherry or GFP) in Arabidopsis roots (Supplementary Fig. 2, Supplementary Table 2). Our AQP module needs to function both in the epidermis and with our synthetic endodermal-like barriers (SRB). Therefore, we evaluated the function of the PIPs and TIPS concurrently with the SRB. As such, expression of PIPs and TIPs were separately targeted to the epidermis (under the *ABCG37* promoter) and the endodermis (via a *SCARECROW* promoter)^24^. Tissue-specific expression was confirmed for five PIPs and two TIPS, followed by drought and salt tolerance assays to evaluate PIP and TIP function (Supplementary Figs. 3-5; Supplementary Table 2). Based on their response to the two abiotic stresses (Supplementary Table 2), two aquaporin PIPs (AoPIP1.2, BgPIP2) and one TIP (TsTIP1.2) were selected and combined to form our AQP module. Plants with both the AQP and the SRB modules show their co-expression abolishes chlorosis and stunting seen in plants with the synthetic epidermal barriers alone (Fig. 3).

### Osmolyte Module (OSM) enhances water retention

One way salt damages plants is from osmotic stress, with osmolyte accumulation known to contribute to plant salt tolerance^25^. Mangroves naturally produce osmolytes thought to be essential for growth in seawater and hypersaline environments^26–28^. Hence, we designed an Osmolyte module (OSM) to allow Arabidopsis plants to tolerate osmotic stress and retain freshwater (see also accompanying manuscript, Ternes et al.). We directed biosynthesis of an osmolyte, mannitol, to the xylem using refactored *AgM6PR* and *AgMAT2* genes from celery (*Apium graveolens*) for mannitol biosynthesis and transport, respectively. We targeted their expression to the xylem parenchyma (VAC, vessel associated cells), using the Arabidopsis *XCP1* and *CesA7* promoters. The VAC’s cell wall typically has half-bordered pits, allowing contiguous contact with water-conducting xylem vessel elements^29^. The OSM module design is to enhance water retention by increasing the xylem’s solute potential. If our OSM module functions as designed, the engineered plants should exhibit enhanced water movement. We tested this by measuring guttation, a natural, passive process that occurs under excessive root pressure^30,31^. Plants expressing the OSM module guttated approximately 30% more water than controls (Extended Data Fig. 4), indicating that the module is functional.

### Rational engineering of salt tolerance

The three modules (epidermal barriers, aquaporins and osmolyte) were combined to produce a Synthetic Salt Tolerance (SST) gene circuit and introduced into Arabidopsis plants with one modification: PIP and TIP expression is controlled by the *At*KC1 promoter, targeting expression to the epidermis and to a lesser extent throughout the root (Supplementary Table 1, Extended Data Fig. 2). Detailed analysis also shows the *At*KC1 promoter directs expression throughout the plant, including in living vasculature and leaf mesophyll tissues^32,33^. Two SST versions were produced: SST1: with genetic components split between two T-DNAs (initial analysis); and SST2 (most studies), with all genetic components in a single T-DNA. Both SST lines had equivalent responses, but because segregation of two T-DNAs introduces added difficulties, we focused on SST2 lines. SST plants were grown in short days for 28 days prior to salt treatment (Materials and Methods, Supplementary Fig. 8A). To screen our SST transgenic plants for tolerance to high levels of salt (600 mM, comparable to seawater) we used a 100 mM NaCl salt step-up, two days at each concentration for a 10-day period, followed by an additional 6-8 days in 600 mM NaCl (Materials and Methods, Supplementary Fig. 8B). Hence plants are in saline conditions under low humidity (10-23%) for at least 16-18 days with the last 6-8 days in 600 mM NaCl (or as notated; details of plant growth and salinization: Materials and Methods, Supplementary Fig. 8A-C).

Primary transgenic plants were selected based on overall plant health, expression of all genes in the engineered circuit, and the presence of synthetic epidermal root barriers (Extended Data Figs. 3, 5 &6). Plant health was assessed both visually and by analysis of chlorophyll fluorescence kinetics, a well-characterized, quantitative trait known to be disrupted by salt stress^34^, which we augmented with Principal Component Analysis (PCA)(Materials and Methods, Supplementary Data 4-6). Quantitative analysis of QY_max_ (F_v_/F_m_, the most predictable parameter of stress) shows that it was possible to select SST plants grown with 600 mM salt that are not statistically different from Col-0 controls grown with tap water (Fig. 4A). Visual assessment of plants, controls (Col-0) and engineered plants treated with 600 mM NaCl for 16-18 days shows that both Col-0 and SST plants were affected by 600 mM salt, yet SST2 lines bolted, flowered and formed seed while Col-0 plants are increasingly stressed and died (Fig. 4B-C). Even with continuous growth in 600 mM NaCl, plants engineered with the SST gene circuit formed mature seed, (Fig. 4, Extended Data Fig. 8) with similar phenotypes observed in plants treated with 400 and 500 mM NaCl (Extended Data Fig. 7). However, the ability to form mature seed with continuous growth in high levels of NaCl declined from 400 to 600 mM NaCl (Extended Data Table 1). In contrast, the same treatment, Col-0 plants showed strong multiple signs of salt stress, including cessation of growth, anthocyanin accumulation, failure to bolt, reduced photosynthesis and eventual death at all salt concentrations tested (Fig.4, Extended Data Fig. 7&8). Homozygous SST2 plants were used for all subsequent analyses.

**Fig. 4.**
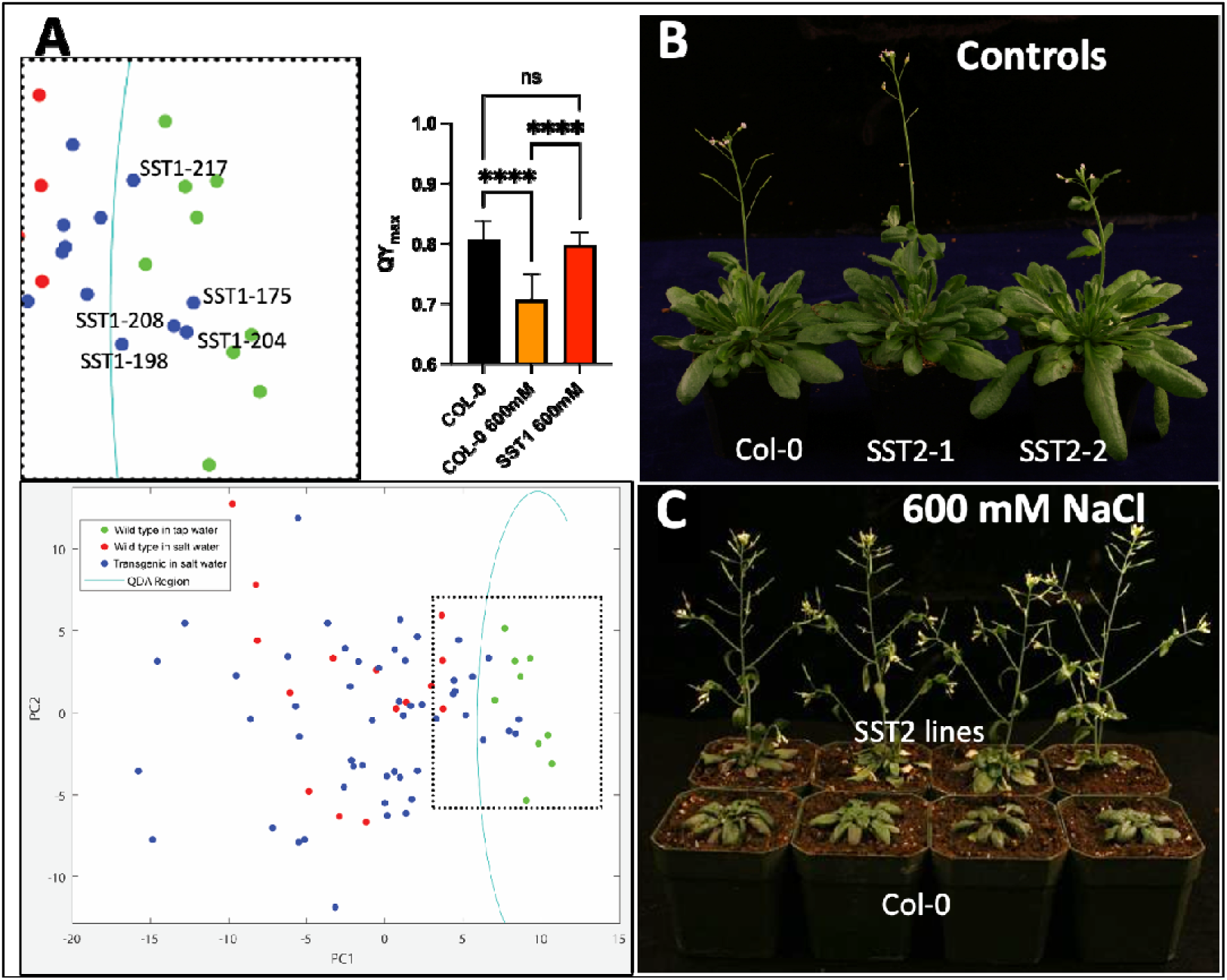
Screening and selection of plants with the Synthetic Salt Tolerance (SST) circuit using chlorophyll kinetics and visual analysis. **a.** Principal Component Analysis (PCA) with the boxed area zoomed in at the top; and plot showing significant differences (P<0.0001) in photosynthetic efficiency (QY_max_) between salt-treated (600 mM) SST1 and control plants (Col-0) treated with water or 600 mM NaCl. b. SST2 plants grown normally with tap water show no adverse effects of the gene circuit. c. SST2 plants treated with 600 mM NaCl and photographed after the last salt treatment. Both SST2 plants and Col-0 control plants are stunted, but SST2 plants can flower and set seed whereas control plants grow increasingly stressed and die.

### Engineered SST2 plants are primed for salt exposure

Salt exposure is well known to alter Arabidopsis gene expression^25^. We used RNA-seq analysis of root tissues from Col-0 and SST2 plants under control conditions (water treated) or treated with 200 mM NaCl. We confirmed numerous up-regulated and down-regulated genes in Col-0 plants treated with salt (Fig. 5A, Supplementary Data 7). Scatter plots comparing Col-0 salt response to tap water treated SST2 plants unexpectedly showed comparable responses (Fig. 5B). Without exposure to salt, SST2 plants up-regulated 38 genes and downregulated 54 genes in common with salt-treated Col-0 (Fig. 5C, Extended Data Table 2, Supplementary Table 5, Supplementary Data 7). Analysis of the top altered genes in SST2 plants showed intricate and epigenetic responses (Extended Data Table 2). Upregulated genes include a protease inhibitor– like protein (Drought-Repressed 4), typically expressed in distal root regions and likely altered from our engineered changes to the root. Increased expression of *Anahita*, linked to stress-related fecundity and transposable element methylation, alongside reduced expression of epigenetically silenced targets (*Sadhu3-2*, AT3G42658; AT5G41660), suggests that SST engineering produces broad epigenomic changes. Other upregulated genes include an antisense transcript identified by QTL studies on water availability (AT2G13665), nutrient stress responses (AT2G13665, AT3G52820, AT3G15650), mRNA stability (AT1G04463), a zinc finger transcription factor previously associated with cold and ROS response (AT5G59820) and hormone signaling (*PILS*, AT2G17500; AT5G15500), though the latter are less induced by SST engineering than salt-treated Col-0. Downregulated genes, in addition to those that are targets of epigenetic silencing, include three genes involved in immunity/pathogen responses (homologue of RPW8 (HR4), AT3G50480; LECRK-1.5, AT3G45430; SCOOP, AT5G44568) and metabolism, and plant cell wall stability genes (AT3G47010, AT3G47380). These data show that our SST2 engineering prepares or produces a pre-emptive salt tolerance response.

**Fig. 5.**
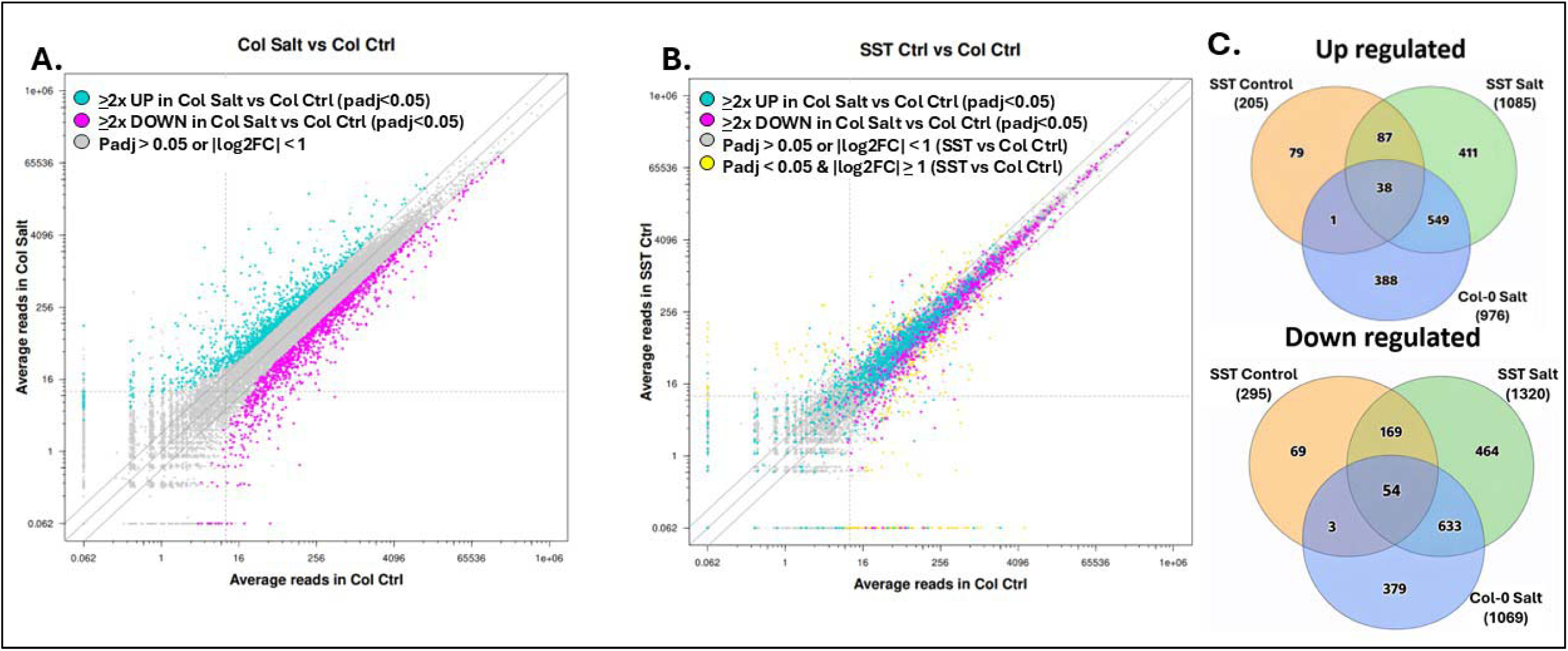
RNA-seq analysis of transcriptional responses to salt stress in SST2 plants. RNA-seq identified DEGs (differentially expressed genes) from Arabidopsis root tissues from control (Col-0) and SST2 plants grown normally with tap water (Control conditions) or with 200 mM NaCl (salt). **a.** Scatter plot showing salt-induced gene expression changes in Col-0 following treatment with 200 mM NaCl. Scatter plots: magenta dots denote downregulated genes; cyan dots denote upregulated genes. **b.** Scatter plot showing expression pattern in Col-0 salt treated plants compared to SST plants treated with water (no salt exposure). **c.** Venn diagrams showing the overlap of upregulated and downregulated genes between water treated SST2 plants and salt-treated Col0 plants.

While our analyses suggest that SST2 engineering leads to a high level of salt tolerance with continuous exposure to 400-600 mM salt, SST2 lines showed increased stress; flowering and seed set declined (Extended Data Table 1). Hence, we analyzed the ion content in shoots and roots with Inductively Coupled Plasma Optical Emission Spectroscopy (ICP-OES) (Supplementary Data 8). Plants were grown and treated with tap water or 200 mM NaCl for two weeks and data normalized to either Col-0 plants treated with tap water (Fig. 6) or to individual lines (i.e., Col-0, SST) treated with tap water (Extended Data Fig 9). With tap water alone, Col-0 plants accumulated 1.5-fold more sodium than SST2 lines in both shoots and roots, except for SST2-1 roots, indicating that our engineered components largely reduce sodium accumulation (Extended Data Fig 9). When grown with 200 mM NaCl (Fig. 6A, Extended Data Fig 9), three out of four SST lines showed significantly reduced sodium accumulation in their shoots, with lines SST2-2 and SST2-5 showing 20-24% reduction in shoot sodium levels. Line SST2-1, showed a similar, though not significantly different, reduction. In roots, line SST2-1 accumulates significantly less sodium (Fig. 6B), while all other SST2 were not significantly different from Col-0 control plants. One possible reason for the observed sodium accumulation in SST2 shoots is the expression of aquaporins directed by the AtKC1 promoter, which is also active in shoots and leaves, potentially leading to higher accumulation of sodium chloride in these tissues. Indeed, expression of *Thellungiella salsuginea* TIP1;2, used in our AQP module, has been reported to facilitate the entry of Na+ ions into plant cell vacuoles under high salinity conditions^35^.

**Fig. 6.**
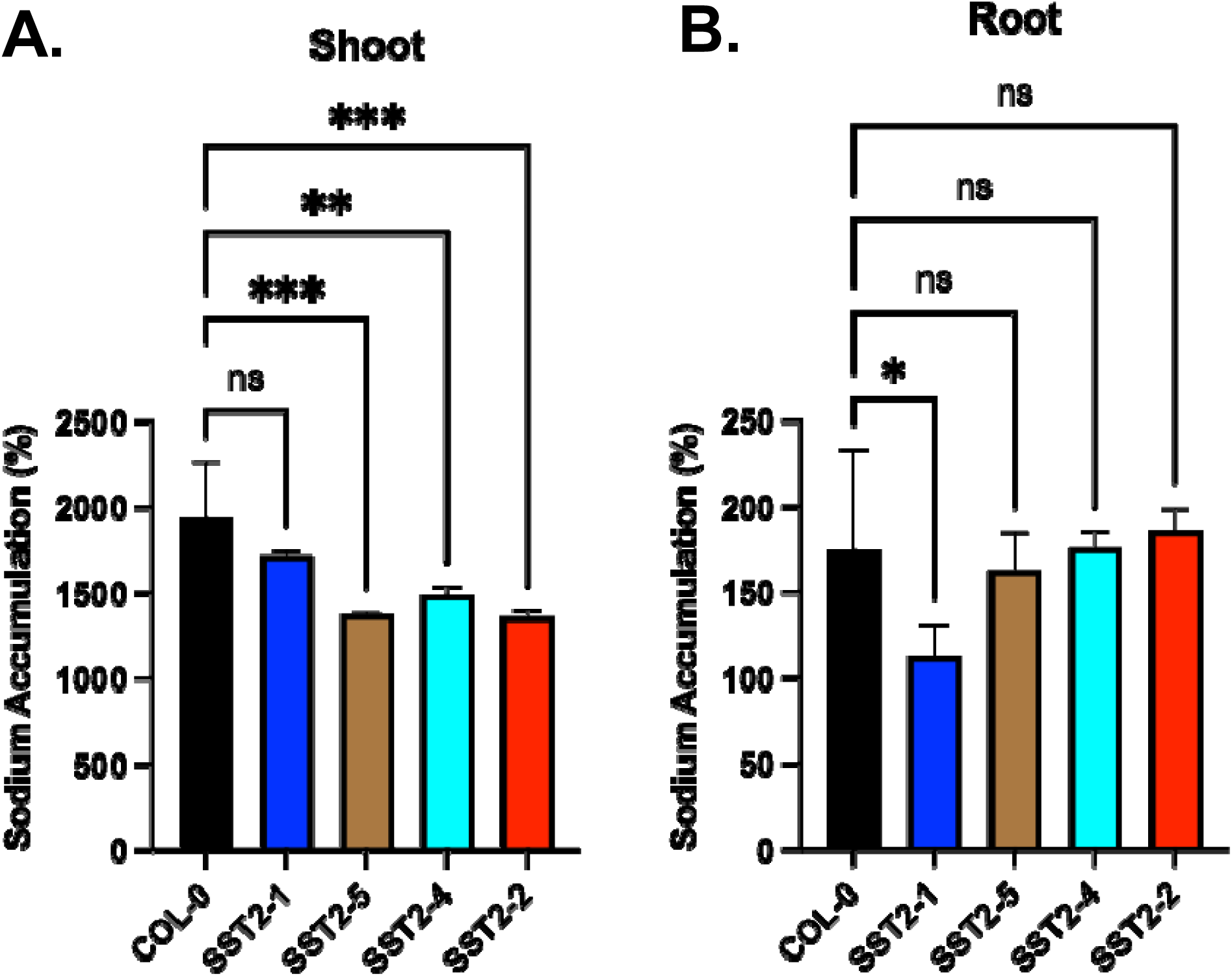
ICP analysis of sodium accumulation. Data are normalized to Col-0 plants treated with tap water. Data are plotted as percent sodium accumulation. **a.** Shoots treated with 200 mM NaCl. **b.** Roots treated with 200 mM NaCl. One-way ANOVA. Dunnett’s post-hoc multiple comparisons test. *P<0.05, **P<0.01, ***P,0.001, ****P<0.0001, and, ns, no significant difference.

ICP analysis also provided insight into how the SST circuit impacted plant ion transport broadly. Notably, SST2-1 plants contain only 11% of iron levels relative to controls, while other independent lines have 33-64% of Col-0 iron levels in their shoots (Supplementary Fig. 16). The reduction in iron in SST2 shoots is consistent with the observed chlorosis (Fig. 3). Interestingly, treatment with 200 mM NaCl, significantly increased iron content in SST2 shoots, but not in roots, in all lines (Supplementary Fig. 6). Magnesium, calcium, and most ions analyzed followed a pattern similar to iron (Supplementary Fig. 6, Supplementary Data 8). However, potassium levels were not significantly different in plants treated with water or with 200 mM NaCl, except for SST2-1 salt-treated roots (Supplementary Fig. 6, Supplementary Data 8).

### SST circuit confers salt tolerance in Rice

To test the general applicability of our approach, we re-engineered the SST circuit for expression in rice, *Oryza sativa*, variety Kitaake (Materials and Methods). We produced a synthetic salt tolerant gene circuit for rice, OsSST1, which relied on curated literature data to select appropriate promoters for gene expression comparable to those used in Arabidopsis (Supplementary Fig. 7, Supplementary Table 7, Supplemental Data 1). We screened second generation OsSST1 rice plants at the young stage when they are highly sensitive to salt, using plants grown in soil under greenhouse conditions with pots placed in large tanks of tap water or salt water (Extended Data Fig. 10 & 11, Materials and Methods 4D, Supplementary Fig. 9). Salt concentration was increased every two days in 100 mM increments until levels reached 600 mM NaCl. Plants were maintained at 600 mM NaCl for 7-10 days and evaluated (17-20 days in salinization conditions). Control Kitaake rice plants exposed to 600 mM NaCl, had brown and shriveled leaves, and the plants failed to recover when transferred to tap water indicating Kitaake plants are highly susceptible to salt stress (Fig. 7)(Standard Evaluation Score, SES, 9)^36^. In contrast, OsSST rice plants only show leaf damage on the older leaves, with some overall stunting, yet have the ability to grow in high salt levels with green and expansive leaves (Fig. 7B) indicative of a SES of 3-4, or tolerant. Moreover, OsSST plants recovered when transferred to a tap water tank and set seed normally.

**Fig. 7.**
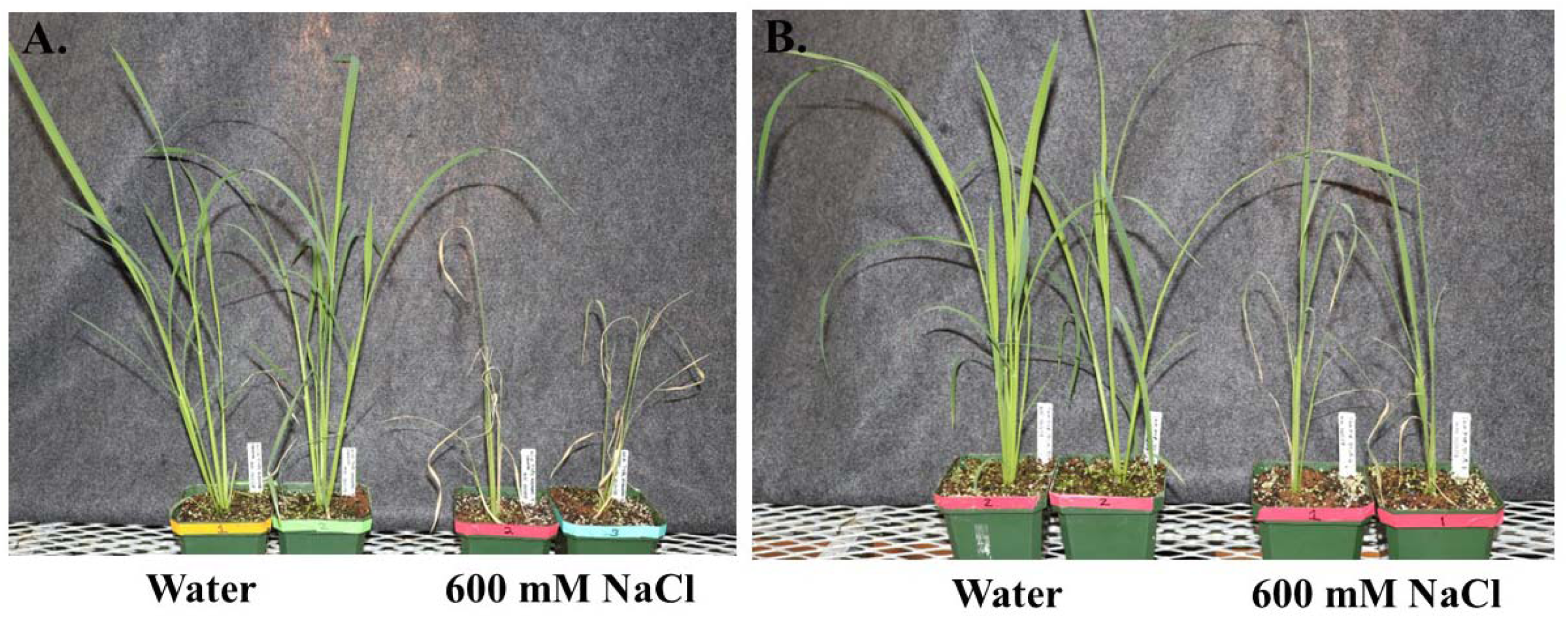
Rice OsSST gene circuit confers salt tolerance in rice. **a**. Kitaake Control Plants treated with water or with 600 mM NaCl. **b**. OsSST engineered rice plants treated with water or with 600 mM NaCl.

## Discussion

Our strategy for improving plant salt tolerance followed a blueprint from mangroves, where apoplastic barriers are duplicated with symplastic transport and osmotic protection enhanced. These genetic modules produced highly salt tolerant Arabidopsis and rice while providing insights to mechanisms and genes underlying tolerance. Our SRB module (suberin/CS) expands endodermal-like barriers radially to the epidermis and distally protecting sensitive root regions.

Yet, when the synthetic barriers are used alone, plants can be chlorotic and stunted. We alleviated this by adding aquaporins, though the TIP we selected may have increased salt accumulation in the shoots, suggesting optimization of TIPS or promoters could lead to improvements. Furthermore, we added the OSM (osmolyte) module to address osmotic stress, collectively the three components produced Synthetic Salt Tolerance (SST) plants capable of growth and seed production under continuous seawater-level salinity (600 mM NaCl). Nonetheless, prolonged salt exposure still produces stress and our elemental analysis suggests that improvements to the epidermal barriers and/or vacuolar sequestration may be needed. Our results also show that synthetic epidermal suberin deposition remains environmentally regulated^37^, highlighting opportunities to decouple and enhance this response. The ability to direct ectopic suberin synthesis and accumulation in roots with our phospho-mimic MYB41^S251D^ opens broader applications that include enhancing resistance to biotic and abiotic stresses.

Transcriptomic analysis of SST2 plants indicates that even when not treated with salt our SST2 plants show expression patterns similar to Col-0 (wildtype plants) treated with salt. This pre-emptive transcriptional response to salt stress is like that of several studies that reported plant transcriptional priming from mild stress or chemical exposure that cause epigenetic, transcriptomic, proteomis and metabolomic^38–40^ alterations. At times these alterations include modulators (e.g., NPR1 and MYB30)^38–40^ that result in transgenic plants pre-disposed to mount a stronger transcriptional response^41,42^. While SST2 plants had not been previously exposed to salt, the pre-emptive transcriptional responses could be a result, in part, from MYB41^S251D^ expression. Numerous genes observed DE genes may enhance plant salt tolerance. While some changes, such as GAE4, involved in cell wall remodeling, may be specific for our engineered changes e.g., synthetic epidermal barriers, other genes involved in epigenetic changes, pathogen responses, water availability, mRNA stability (AT2G13665, AT1G04463, respectively), and metabolism are noteworthy and may guide future efforts.

## Conclusion

This work advances our understanding of plant salt tolerance and establishes a modular framework for engineering stress tolerance. In a related work (Ternes et al.), we show how these components can be integrated with a synthetic plant waterpump circuit to enable plants to produce freshwater.

## Acknowledgements

This work was supported by funding from DARPA (HR0011-16-2-0005). We thank Professor Tai Montgomery and Shuo Liu for help with the transcriptomic analysis.

---

**Extended Data Fig. 1.**
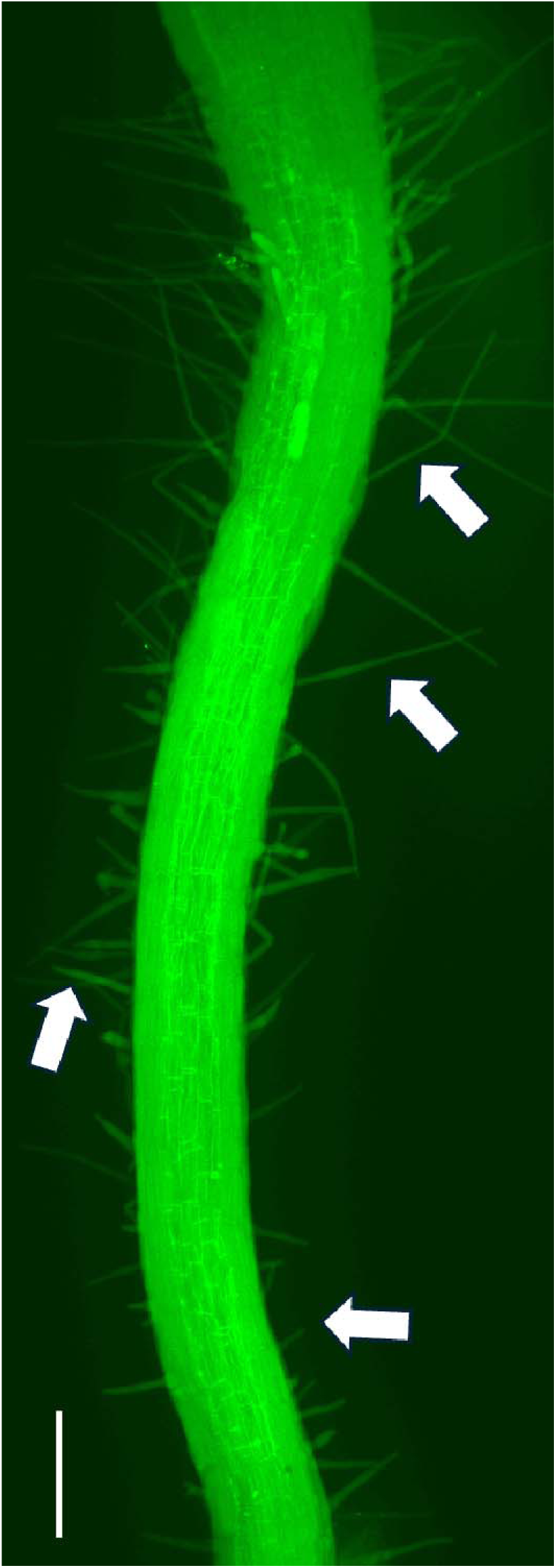
Transgenic plants expressing ABCG37::MYB41^S251D^ accumulate suberin – closeup to show root hairs. Figure 2D (main text) is expanded to clearly show expression in the root hairs. Longitudinal view, whole mount from plants transformed with ABCG37::MYB41^S251D^. Suberin staining (Fluorol Yellow, green) in mature roots and root hairs. Scale bar = 10 mm. Arrows indicate root hairs.

**Extended Data Fig. 2.**
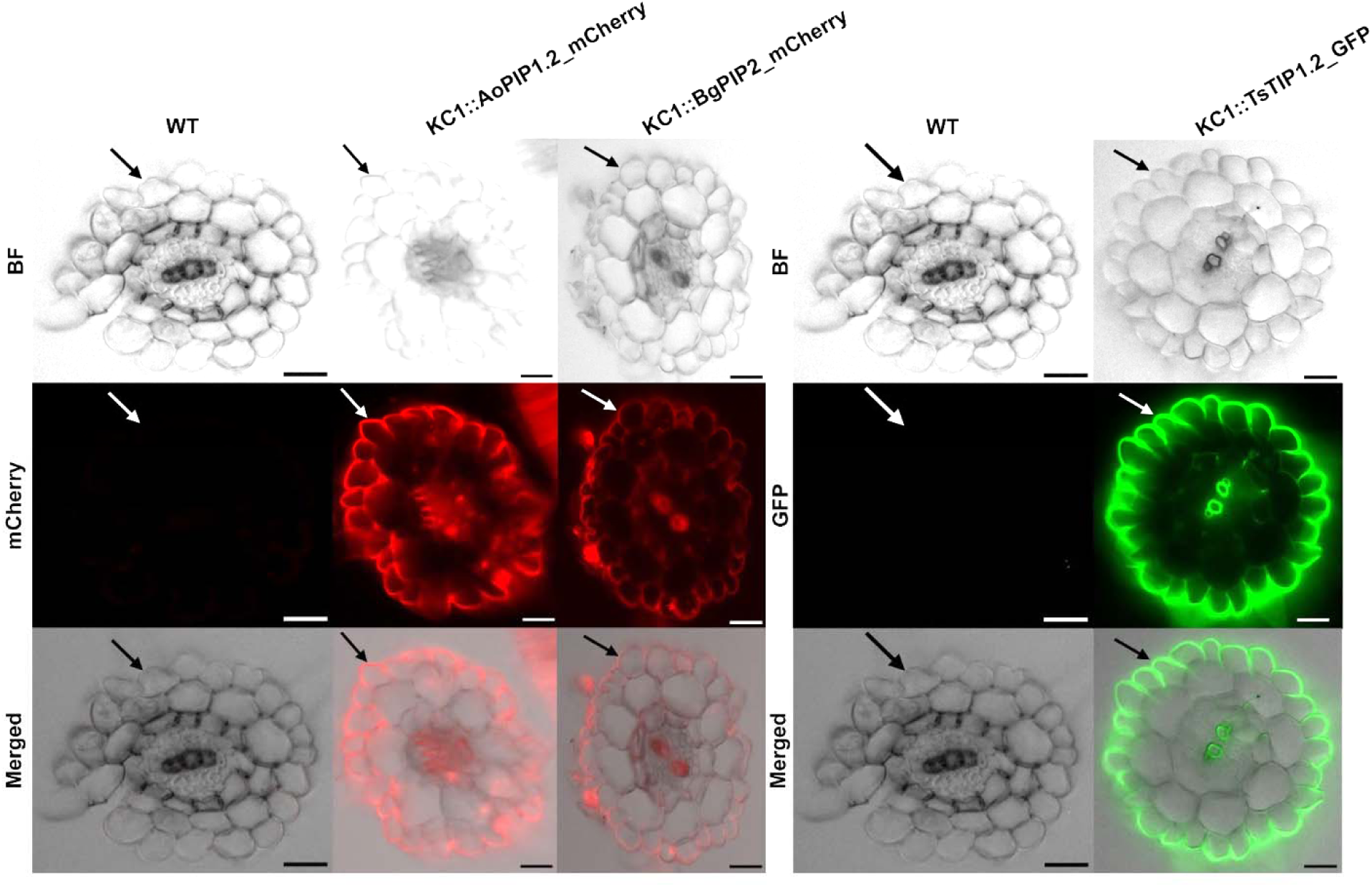
Targeting and expression of mangrove synthetic transport system components in Arabidopsis roots with the KC1 promoter. Root cross sections of Col-0 and transgenic plants containing the specific aquaporin genes used in the SSTs (PIPs and TIP) under control of the KC1 promoter, which directs expression to the epidermis, and to a lesser extent, the cortex and throughout the root (table S1). Aquaporin genes fused to either mCherry or GFP reporter genes. Arrows indicate epidermis. Scale bar = 10 µm.

**Extended Data Fig. 3.**
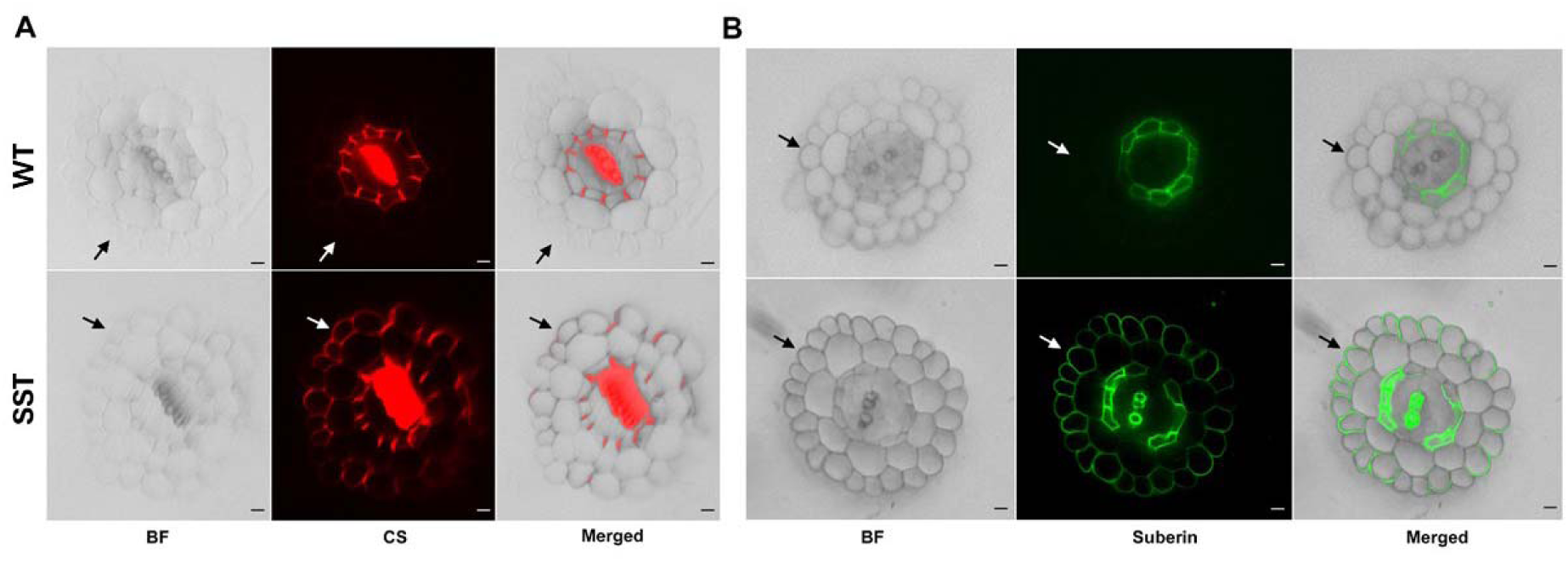
Suberin and CS in root epidermis of SST plants. Root cross-sections of wildtype (WT) and transgenic Arabidopsis plants containing the SST. (**A**) Basic Fuchsin staining showing epidermal Casparian strip production in transgenic roots. (**B)** Fluorol yellow staining shows epidermal suberin production in the transgenic root. Scale bars = 10 μm. BF, Brightfield, CS, Casparian strip. Roots were treated as described in Figure 2 with 1 μm AVG added (See Materials and Methods). Arrows point to epidermis.

**Extended Data Fig. 4.**
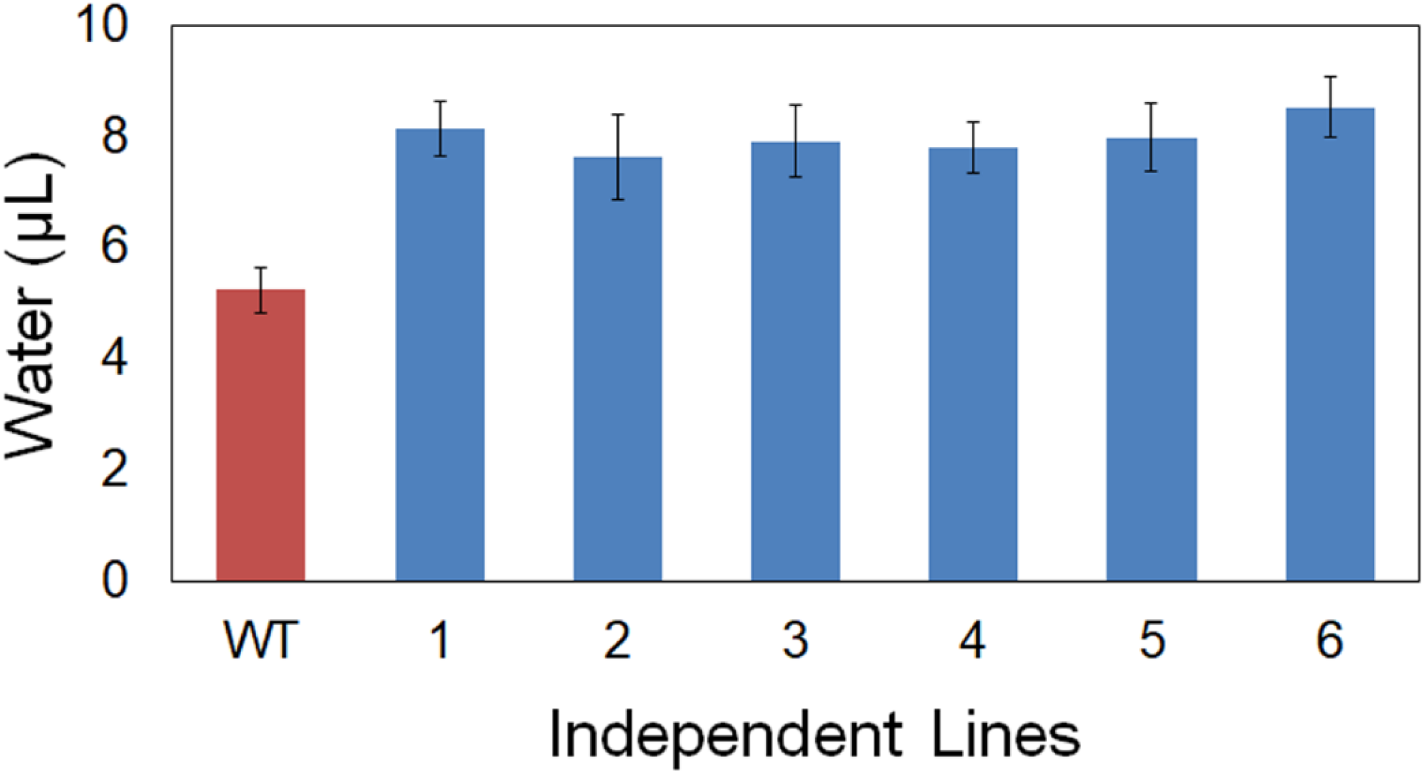
Function of the Osmolyte module in plants. Function of the Osmolyte module in transgenic plants showing approximately 30% enhancement in the volume of water secreted by guttation. All plants treated with tap water. Volumes are normalized for leaf area. All independent OGM lines secreted significantly greater water volumes (Student’s T-Test, p <0.01).

**Extended Data Fig. 5.**
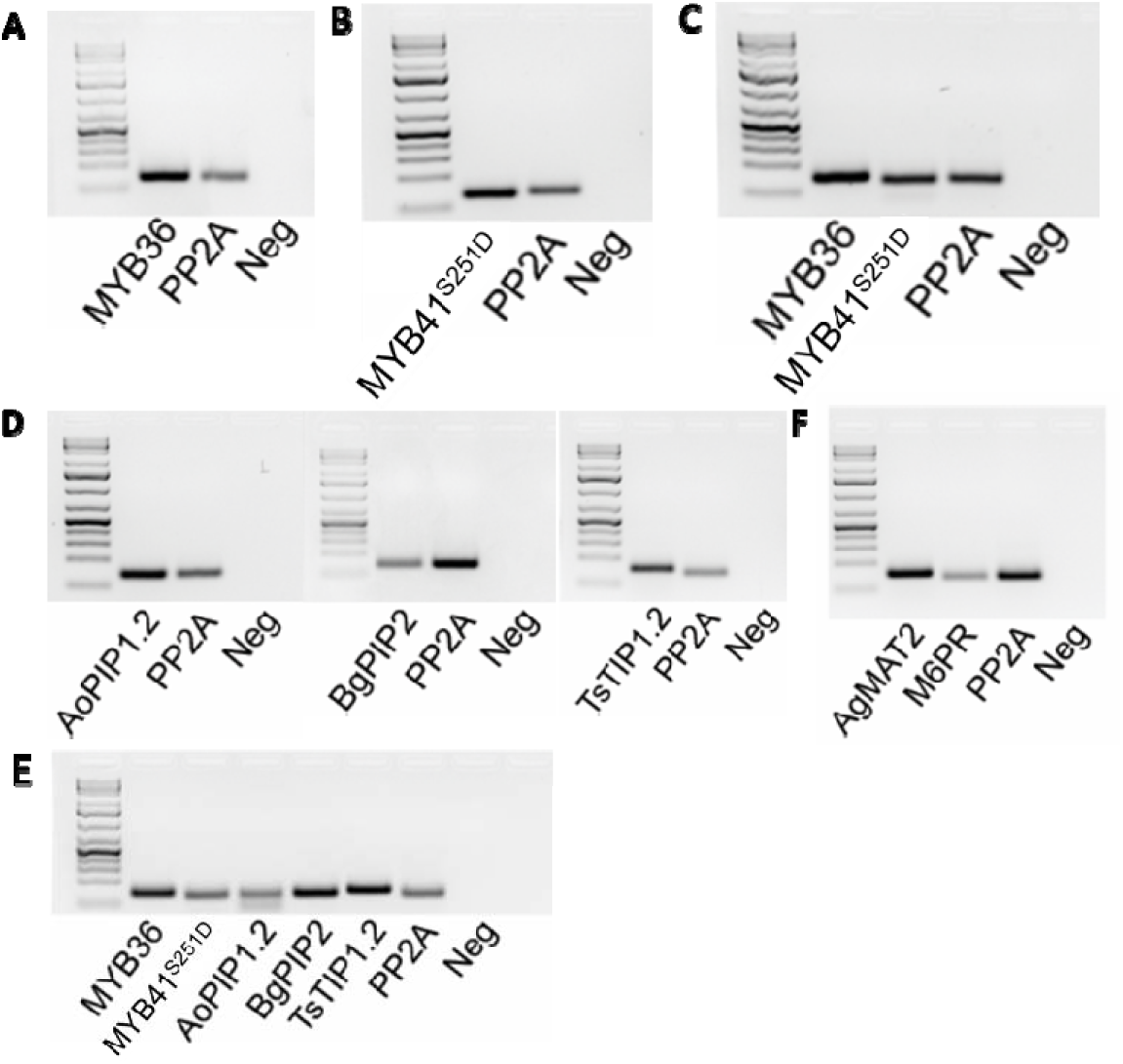
Endpoint RT-PCR analysis verifying expression of transgenes in all transgenic lines described herein. On most panels, PP2A (protein phosphatase 2A) is a housekeeping gene included as a positive control. **(A-C)** Expression of components of the SRB filter-barrier module as assayed with endpoint RT-PCR. (**A**) ABCG37::MYB36, directing expression of the epidermal Casparian strip; plants in Figure 2A. (**B**) ABCG37::MYB41^S251D^, directing expression of epidermal suberin; plants in Figure 2B. (**C**) ABCG37::MYB36 and ABCG37::MYB41^S251D^; plants in Figure 3, center. **(D)** Expression of components of the AQP transport module as assayed with endpoint RT-PCR. Left: KC1::AoPIP1.2, directing expression of the PIP AoPIP1.2. Center: KC1::BgPIP2, directing expression of the PIP BgPIP2. Right: KC1::TsTIP1.2, directing expression of the TIP TsTIP1.2. Plants are found in Figure S6-S7. (**E)** Expression of components of the SRB and AQP modules as assayed with endpoint RT-PCR. Filter-barrier module: ABCG27::MYB36, directing expression of the epidermal Casparian strip, and ABCG37::MYB41S251D, directing expression of epidermal suberin. Synthetic transport system: KC1::AoPIP1.2, KC1::BgPIP2, and KC1::TsTIP1.2, directing expression of the PIP AoPIP1.2 and BgPIP2 and the TIP TsTIP1.2, respectively. Plants are shown in Figure 3, right. **(F)** Expression of components of the Osmotic Genetic Module (OGM) as assayed by endpoint RT-PCR. CESA7::AgMAT2, directing expression of the mannitol transporter in the xylem parenchyma; and XCP1::omega enhancer-M6PR, directing expression of the mannitol biosynthesis gene in the xylem parenchyma. Data in Figure 3B.

**Extended Data Fig. 6.**
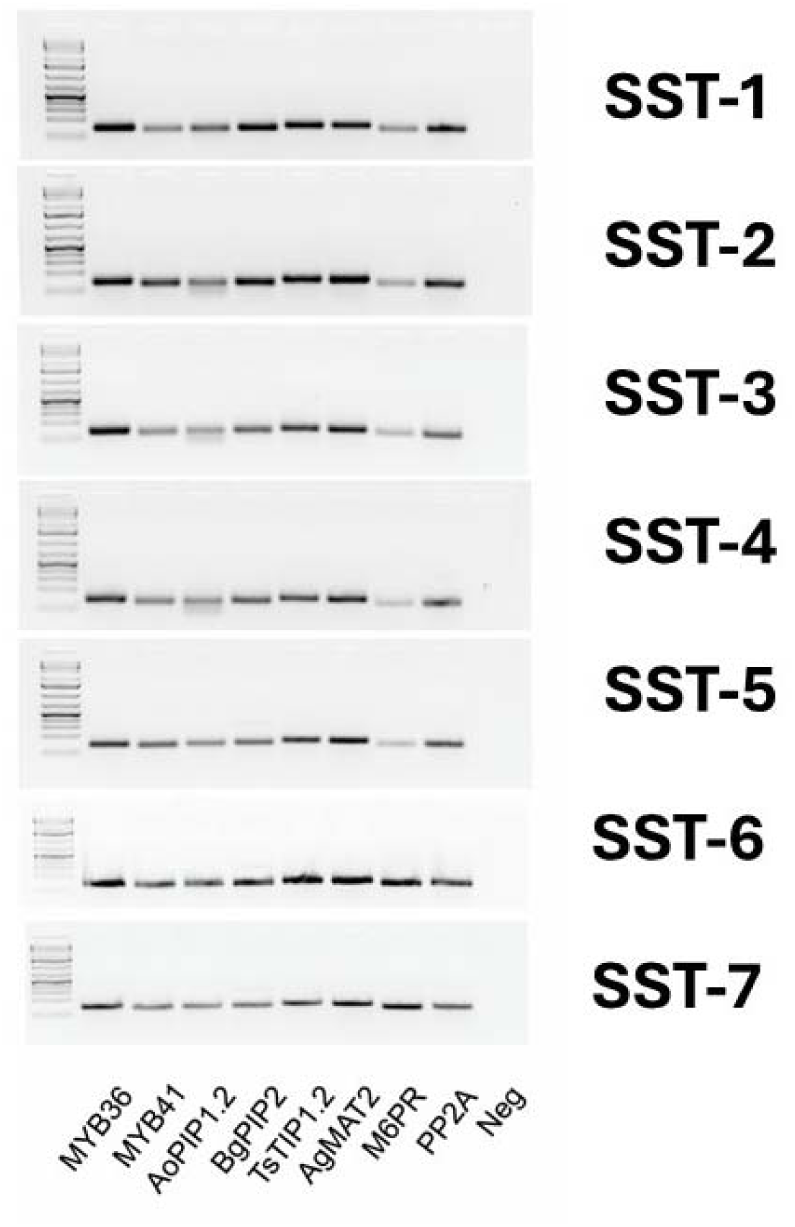
Expression of components of the SST in seven independent lines as assayed by endpoint RT-PCR. SRB: ABCG37::MYB36 and ABCG37::MYB41^S251D^. AQP module: KC1::AoPIP1.2, KC1::BgPIP2, and KC1::TsTIP1.2. Osmolyte Module: CESA7::AgMAT2 and XCP1::omega enhancer-M6PR. SST2-1through SST2-5 are in Figure 5B whereas SST2-6 and SST2-7 are found in Figure 5A. SST-6 and SST-7 lines were not extensively used.

**Extended Data Fig. 7.**
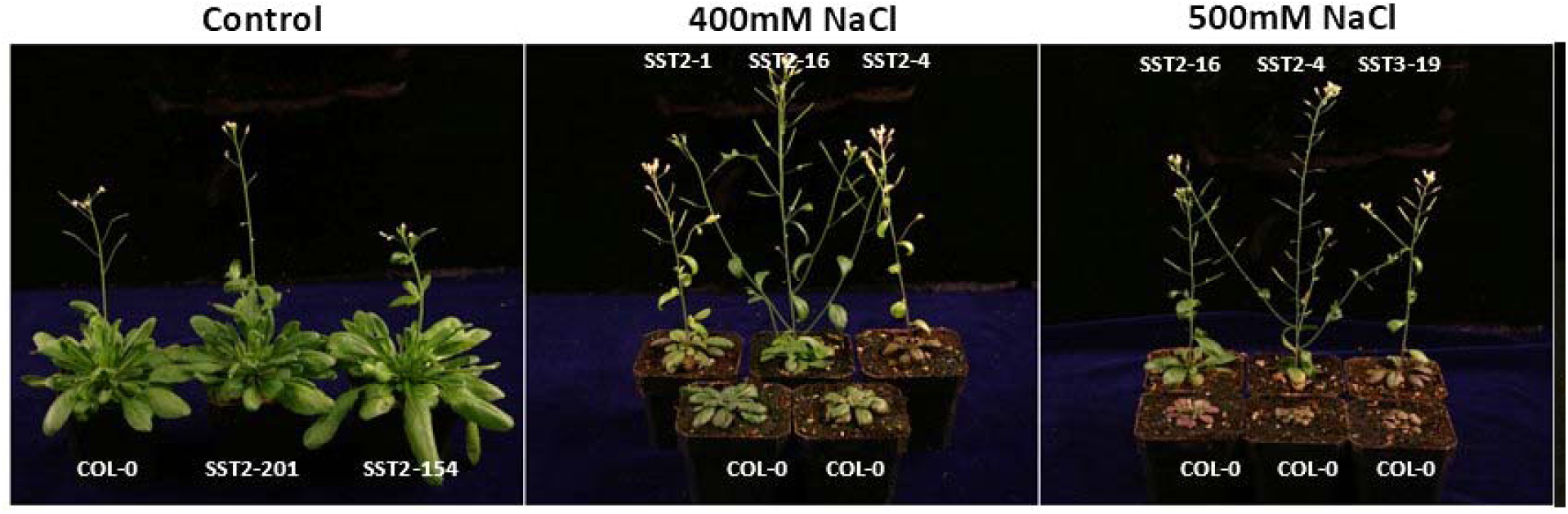
Visual analysis of plants with homozygous for Synthetic Salt Tolerance (SST) Circuit treated with 400 or 500 mM NaCl. SST plants treated with 400 or 500 mM NaCl and photographed after the last day of the salt treatment. Both SST lines and Col-0 control lines are stunted but SST lines can flower and set seed whereas control plants eventually die. SST3 has two T-DNAs and instead of using mannitol in the OSM module, SST3 uses glycine betaine as the osmoticum (see Materials and Methods). The SST3 line showed signs of stress without salt and was not studied further.

**Extended Data Fig. 8.**
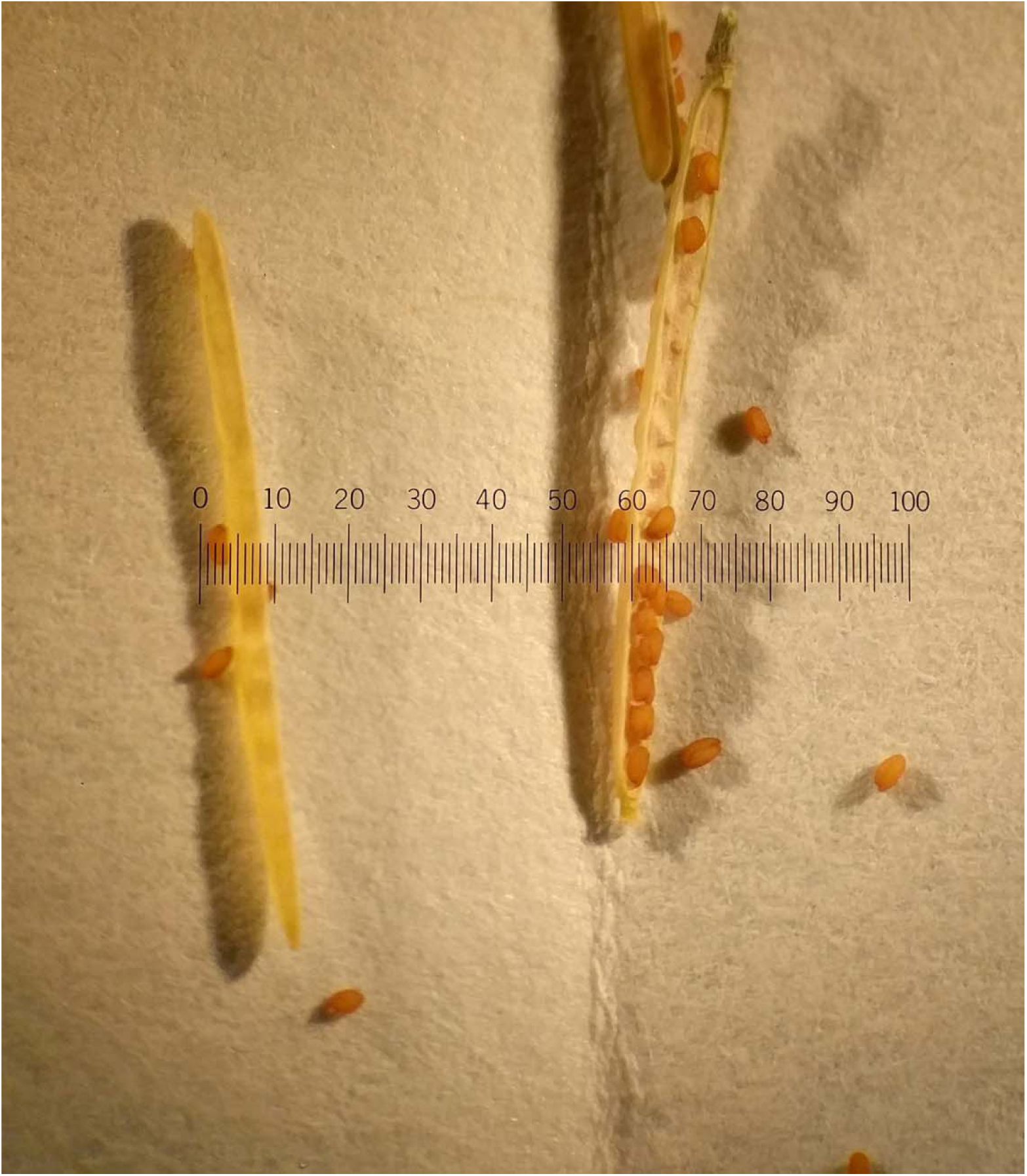
Seed pod (silique) and seed formation in SST2-1 plants treated continuously with 600 mM NaCl (table S3). Entire scale bar (0-100) = 1 cm; each increment is 0.1 mm.

**Extended Data Figure 9.**
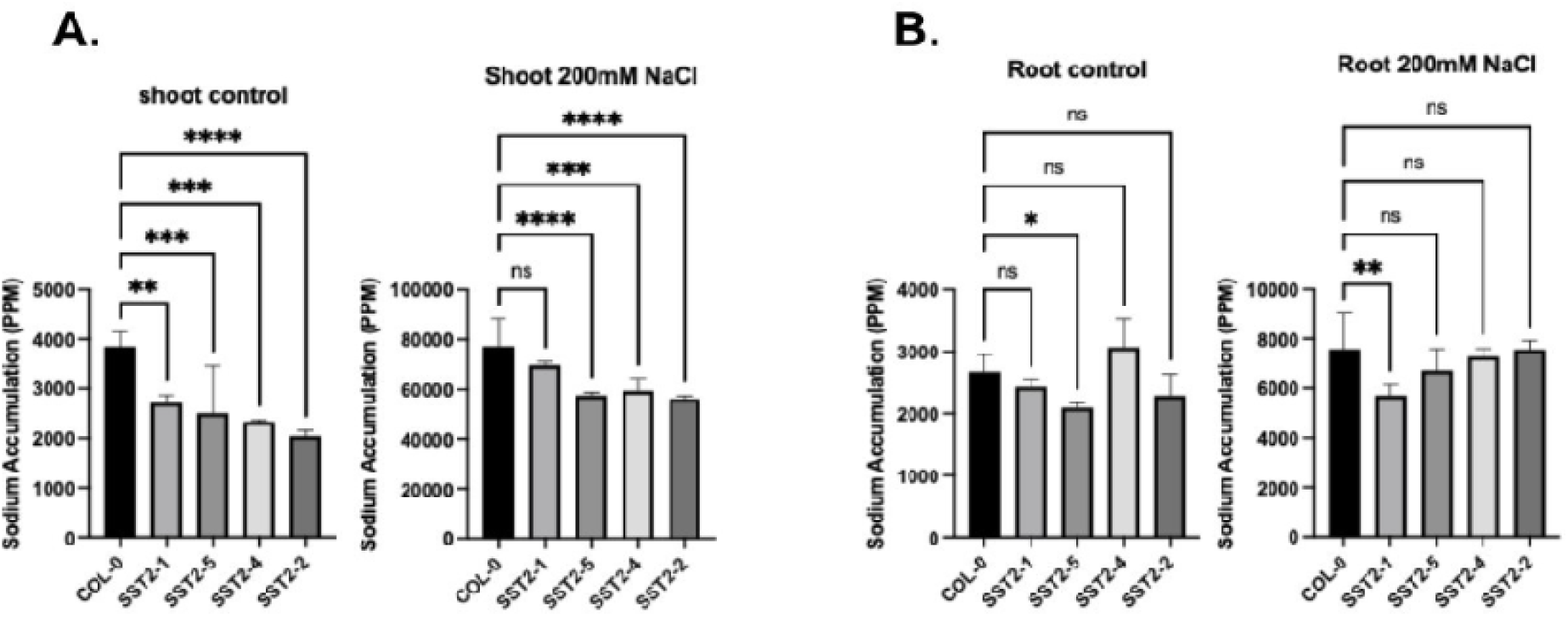
ICP analysis of sodium accumulation, plants grown with water or with 200 mM NaCl. Data normalized to individual controls. A. Shoots, water control and plants grown with 200 mM NaCl. B. Roots, water control and plants grown with 200 mM NaCl. One-way ANOVA. Dunnett’s post-hoc multiple comparisons test. *Indicate P<0.05, **P<0.01, ***P,0.001, ****P<0.0001, and ns signifies no significant difference.

**Extended Data Fig. 10.**
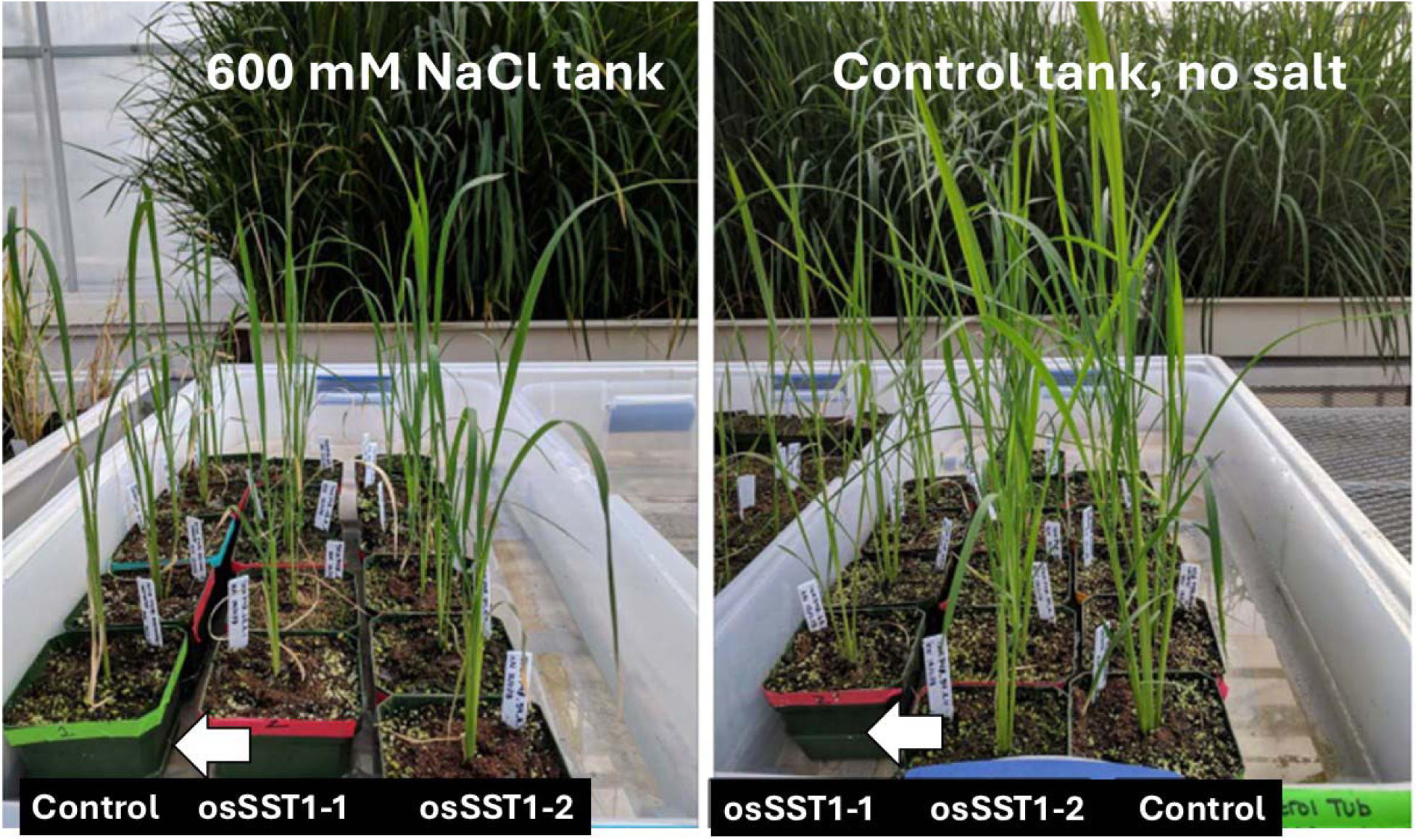
Greenhouse screening of rice containing the OsSST gene circuit. Tank with 600 mM NaCl (left) and control (water) tank, right. Plants photographed 2 days (two) after plants were placed at this level of salt. Control lines and a less tolerant OsSST line (left, center) begin to show damage whereas the tolerant OsSST1-2 line thrives. White arrows indicate levels of 600 mM NaCl (left) or water (right).

**Extended Data Fig. 11.**
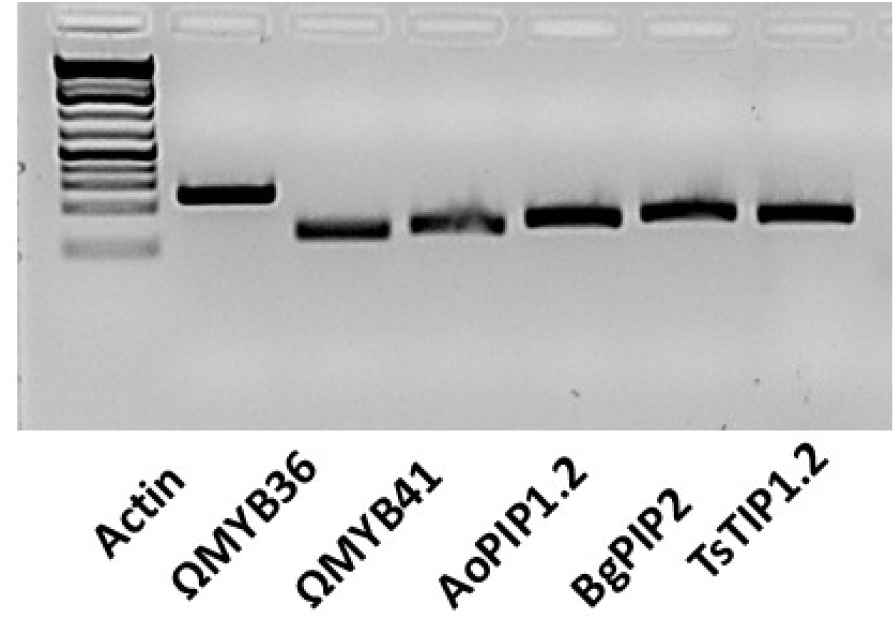
Expression of the OsSST components in rice as assayed by endpoint RT-PCR. Expression was assayed in rice root tips of plants grown without salt treatment.

**Extended Data Table 1.** Seed formation in SST plants continuously grown with high levels of salt. Number of plants for which floral organ initiation, seed production, and if applicable, seed germination was observed in wild type Arabidopsis plants (Col-0) and three transgenic SST2 lines under continuous salt (NaCl) treatment. Fourteen-day old plants were transferred to soil, and salt treatment was started four days after acclimation to soil (18-day old plants). Plants were treated with salt with 100 mM incremental steps until the respective maximum salt concentration (400, 500 or 600 mM NaCl) was reached. Plants were sub-irrigated 2-3 times per week with their maximum salt concentration and evaluated for ability to initiate floral organs and set seed. *NA, not applicable;* plants did not produce seeds.

| Line | 400 mM NaCl<br>Continuous salt treatment |  |  |  | 500 mM NaCl<br>Continuous salt treatment |  |  |  | 600 mM NaCl<br>Continuous salt treatment |  |  |  |
| --- | --- | --- | --- | --- | --- | --- | --- | --- | --- | --- | --- | --- |
|  | Initiated<br>floral organs | Produced<br>seed | Total plants<br>tested | Germination<br>% | Initiated<br>floral organs | Produced<br>seed | Total plants<br>tested | Germination<br>% | Initiated<br>floral organs | Produced<br>seed | Total plants<br>tested | Germination<br>% |
| Col-0 (wild type) | 1 | 0 | 3 | NA | 0 | 0 | 3 | NA | 1 | 0 | 4 | NA |
| SST2-1 | 3 | 2 | 3 | 100% | 3 | 1 | 3 | 70% | 4 | 2 | 4 | 77% |
| SST2-2 | 2 | 1 | 3 | 58% | 2 | 2 | 3 | 9% | 2 | 1 | 3 | 13% |
| SST2-3 | 2 | 1 | 3 | 76% | 0 | 0 | 3 | 61% | 1 | 0 | 4 | 16% |

**Extended Data Table 2.**
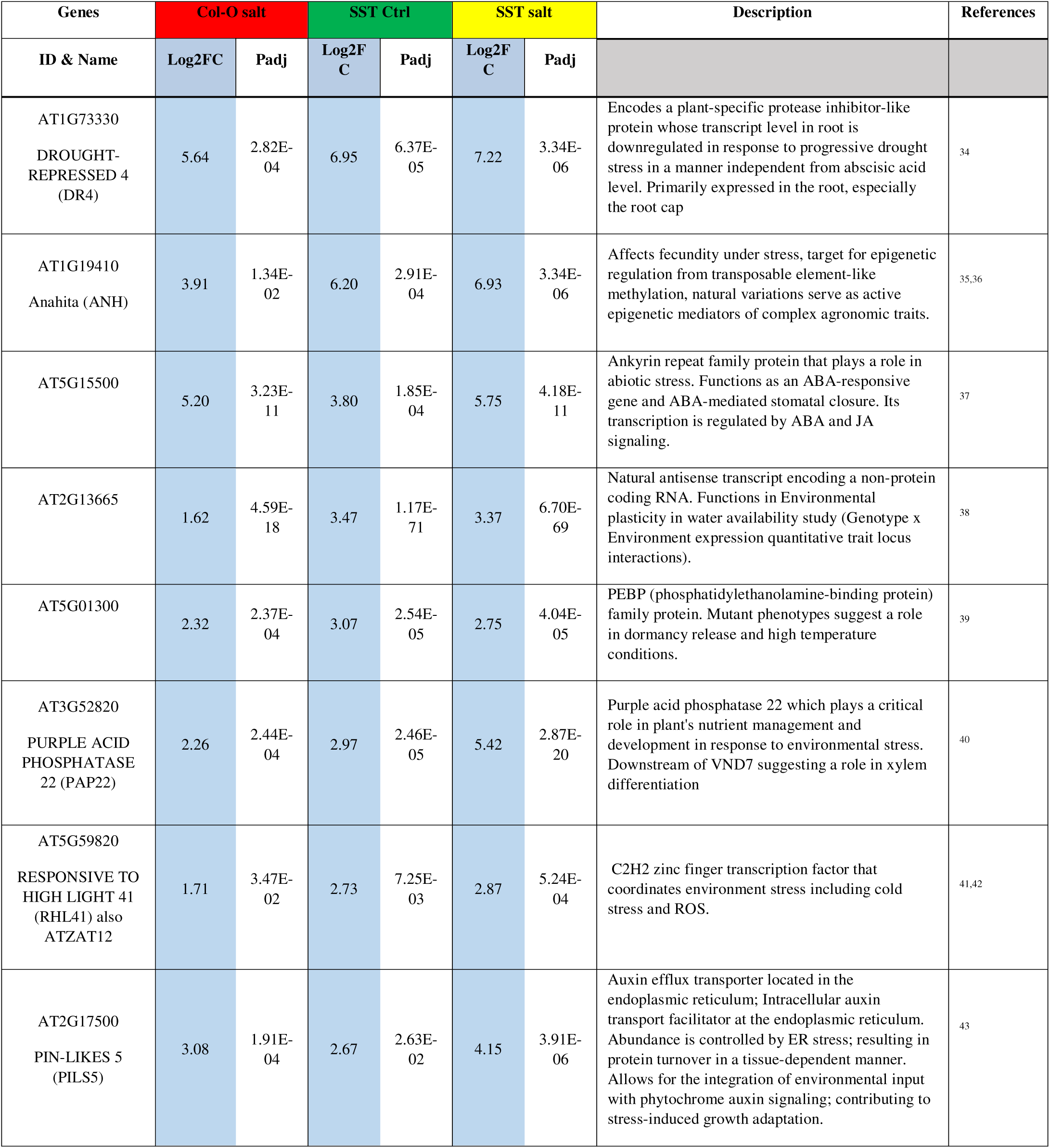
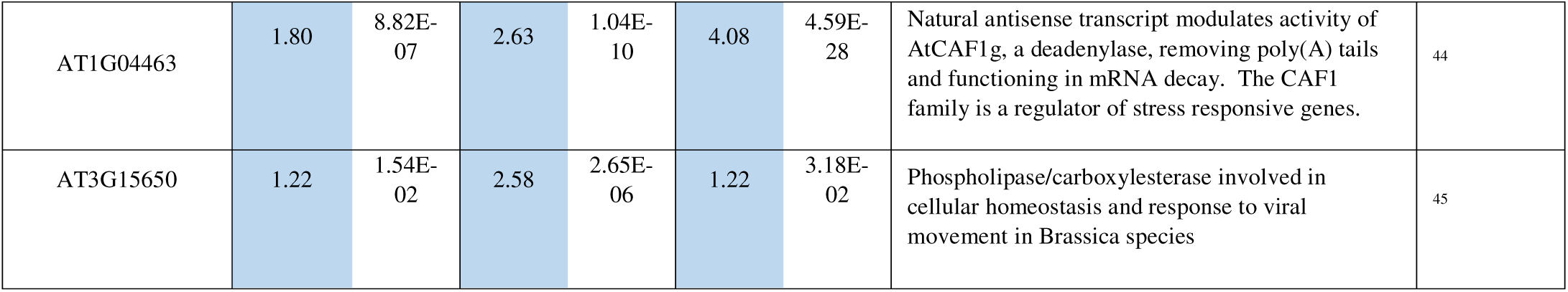
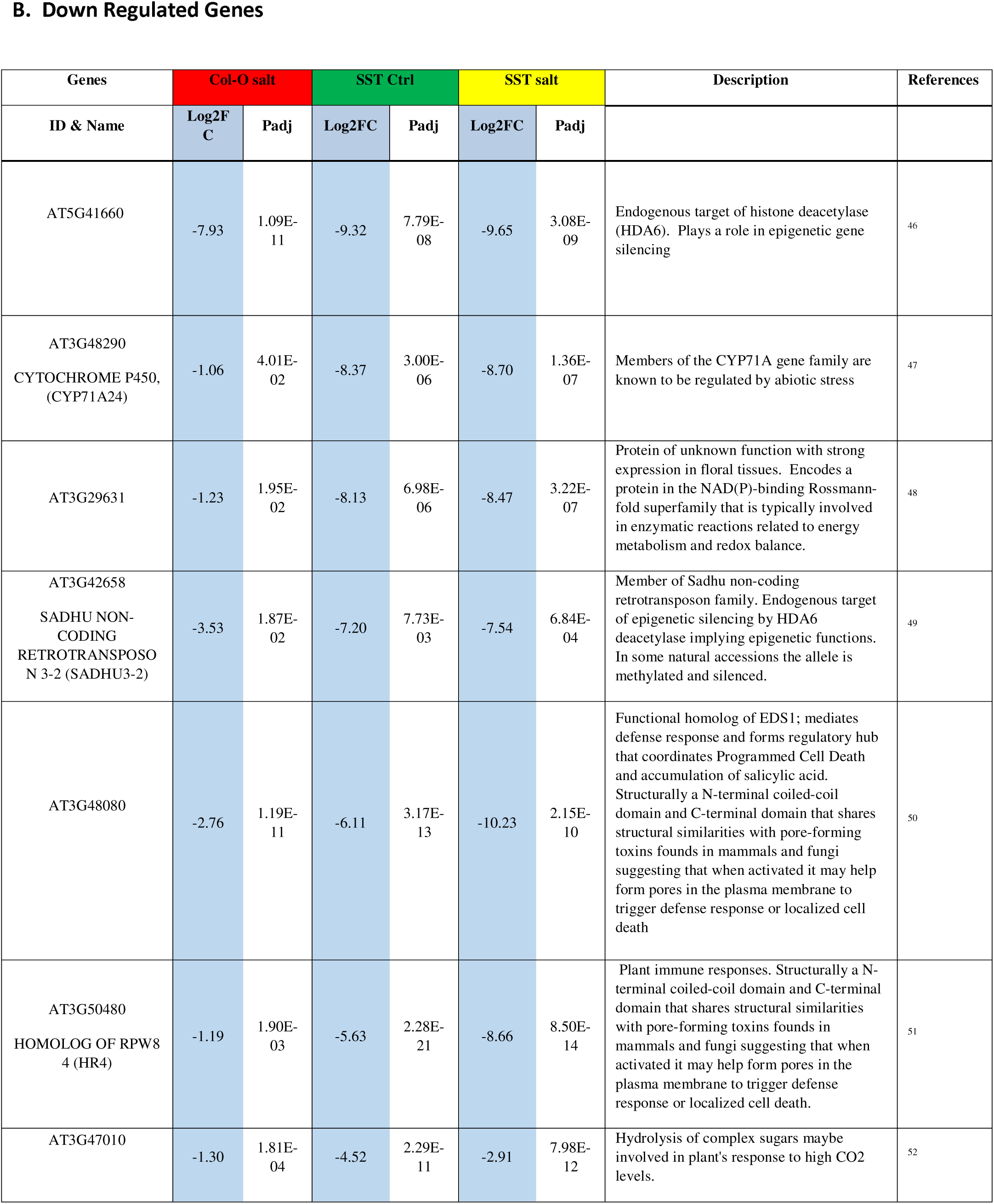
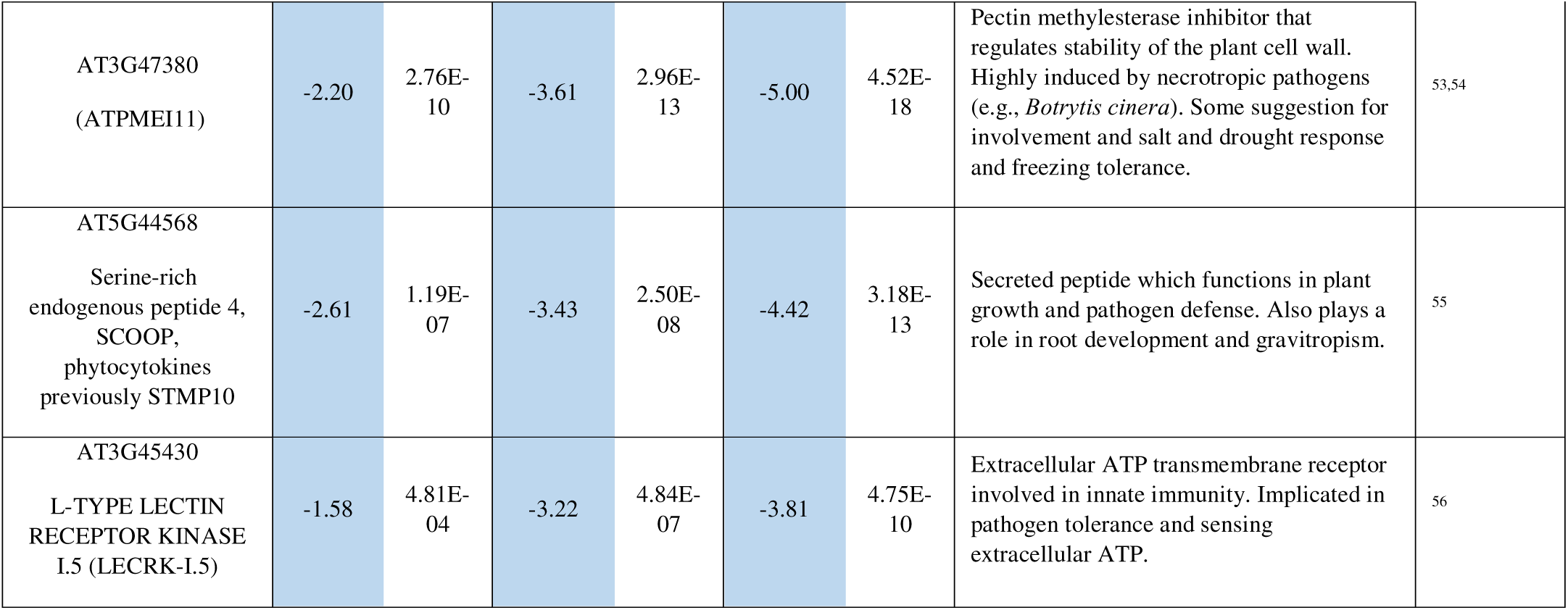
RNA-seq detailed analysis of top 10 up regulated (A) and down regulated (B) genes in SST2 plants versus control plants (Col-0). A. Up Regulated Genes B. Down Regulated Genes.

