## Supplemental Info - Materials and Methods for "A synthetic gene circuit confers salt tolerance in plants"

**SUPPLEMENTARY TABLES**

**Supplementary Table 1. Arabidopsis root promoters used in the Synthetic Root Barrier (SRB).** Because we found some discrepancy with published reports, we experimentally verified each promoter’s function by directing expression of reporter genes (mCherry or GFP) in transgenic plants.

| Promoter Name | Reference | Brightfield image | mCherry or GFP image |
| --- | --- | --- | --- |
| ABCG37/PDR9  ATP-BINDING CASSETTE G37 | (*16*) | 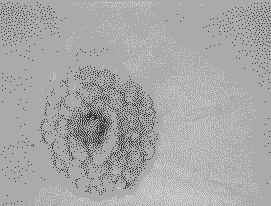 | 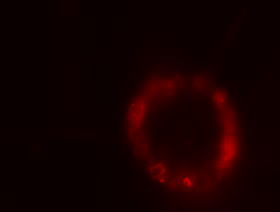 |
| KC1  K+ RECTIFYING CHANNEL 1 | ^31^ | 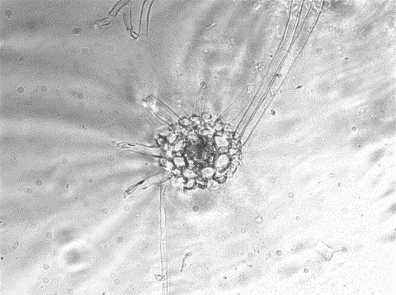 | 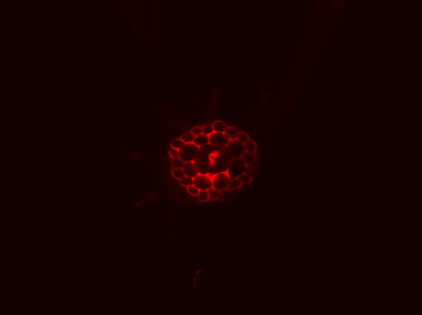 |
| XCP1  XYLEM CYSTEINE PEPTIDASE 1 | ^32^ | 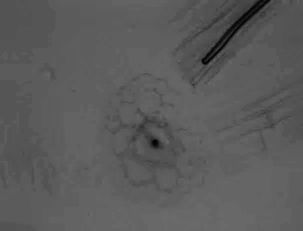 | 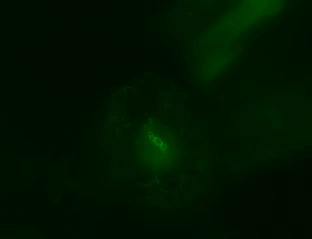 |
| CESA7/IRX3  CELLULOSE SYNTHASE CATALYTIC SUBUNIT 7 | ^33^ | 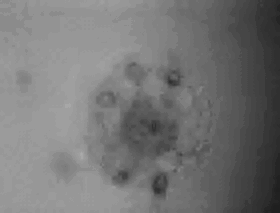 | 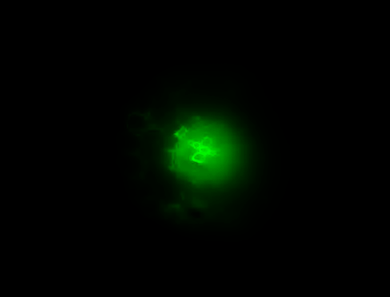 |

**Supplemetary Table 2.** **Functional screening of refactored aquaporins in transgenic Arabidopsis plants**.

Drought and salt tolerance assays were performed to determine if the refactored aquaporins functioned in Arabidopsis. We evaluated transgenic plants with the aquaporins targeted to the endodermis (mimicking our synthetic barriers) and the epidermis for drought and salt tolerance. Aquaporins highlighted in green showed enhanced drought stress tolerance; the aquaporin highlighted in blue gave high salt tolerance with plants subjected to 350 mM NaCl (without salt step up procedure). We selected aquaporins based on the two abiotic stress responses for use in the full SST circuits. See supplementary figs. 3-5 for documentation of drought and salt tolerance. Species IDs: Ac = *Aegiceras corniculatum*, Ao = *Avicennia officinalis*, Bg = *Bruguiera gymnorhiza*, Kc = *Kandelia candel,* Ts = *Thellungiella salsuginea*. *AcPIP2 did not have predicted tissue specific localization. **BgTIP showed a difference between N-terminal and C-terminal reporter protein fusions. The C-terminal fusion protein was not stable, but the N-terminal protein was stable.

|  | Endodermal SCR promoter | | |  | Epidermal ABCG37 promoter | | |
| --- | --- | --- | --- | --- | --- | --- | --- |
| Gene | Fusion protein stable in roots? | Tested for drought tolerance | Drought tolerant lines/ tested lines |  | Fusion protein stable in roots? | Tested for drought tolerance | Drought tolerant lines/ tested lines |
| **PIPs** |  |  |  |  |  |  |  |
| AcPIP1 | no | - | - |  | yes | yes | 5/9 |
| AcPIP2 | yes* | - | - |  | no | - | - |
| AoPIP1.1 | yes | Yes | ¾ |  | yes | no | n/a |
| AoPIP1.2 | yes | Yes | 10/10 |  | yes | yes | 5/7 |
| AoPIP2 | no | - | - |  | yes | - | - |
| AoPIP2.2 | yes | Yes | 3/7 |  | yes | yes | 3/8 |
| BgPIP2 | yes | Yes | 3/8 |  | yes | yes | 5/5 |
|  | Endodermal SCR promoter | | |  | Epidermal ABCG37 promoter | | |
| Gene | Fusion protein stable in roots? | Tested for salt tolerance | Salt tolerant lines/ tested lines |  | Fusion protein stable in roots? | Tested for salt tolerance | Salt tolerant lines/ tested lines |
| **TIPs** |  |  |  |  |  |  |  |
| BgTIP** | no | - | - |  | yes | yes | 5/8 |
| KcTIP | no | - | - |  | no | - | - |
| TsTIP1.1 | yes | Yes | 1/6 |  | yes | yes | 7/8 |
| TsTIP1.2 | yes | Yes | 1/5 |  | yes | yes | 6/8 |

**Supplementary Table 3.** Suberin fold change for lipids and polymeric lipids

|  | **Compound** | **Fold Change (vs Col-0)** | | | |
| --- | --- | --- | --- | --- | --- |
|  |  | **Water** | | **200 mM NaCl** | |
|  |  | **SST2-1** | **SRB2-1** | **SST2-1** | **SRB2-1** |
| Lipids | Behenic acid | 1.20 | 1.58 | 0.94 | 1.46 |
|  | Monoacylglycerol #3 | 0.58 | 0.35 | 0.89 | 2.04 |
| Polymeric Lipids | Methyl *p-*coumarate | 0.71 | 1.35 | 0.80 | 1.20 |
|  | Cinnamic acid-like | 0.97 | 0.11 | 1.82 | 2.71 |
|  | Stearic acid | 5.34 | 2.55 | 2.44 | 0.55 |
|  | Methyl ferulate | 1.06 | 1.55 | 0.85 | 1.25 |
|  | Docosanoic acid (RT 27) | 1.15 | 0.84 | 1.02 | 1.08 |

**Supplementary Table 4.** Suberin fold change calculated when plants were grown normally (with water) versus in 200 mM NaCl.

|  | **Compound** | **Fold Change (Water vs 200 mM NaCl)** | | |
| --- | --- | --- | --- | --- |
|  |  | **Col-0** | **SST2-1** | **SRB2-1** |
| Waxes | Stearyl alcohol | 13.22 | 1.56 | 1.98 |
|  | Palmitic acid | 0.93 | 0.83 | 1.35 |
|  | Phthalic acid-like | 2.57 | 1.24 | 0.81 |
|  | Docosanoic acid, 2,3-propyl ester | 1.29 | 0.93 | 1.99 |
| Lipids | Behenic acid | 1.92 | 1.50 | 1.78 |
|  | Monoacylglycerol #3 | 0.71 | 1.08 | 4.07 |
| Polymeric Lipids | Methyl *p-*coumarate | 1.84 | 2.06 | 1.64 |
|  | Cinnamic acid-like | 0.11 | 0.20 | 2.67 |
|  | Stearic acid | 2.04 | 0.93 | 0.44 |
|  | Methyl ferulate | 3.96 | 3.15 | 3.19 |
|  | Docosanoic acid (RT 27) | 1.26 | 1.11 | 1.63 |

**Supplementary Table 5**. Complete list of genes (A) Up or (B) Down regulated in our untreated SST2-1 plants that are in common with those altered by salt treatment in Col-0 control plants starting after the ten listed above.

**A. Upregulated genes (38)**

| **38 Up Regulated Genes** | | | | | | | | **Description** |
| --- | --- | --- | --- | --- | --- | --- | --- | --- |
| **Genes** | | **Col-O salt** | | **SST Ctrl** | | **SST salt** | |  |
| **ID** | **Name** | **Log_2_FC** | **Padj** | **Log_2_FC** | **Padj** | **Log_2_FC** | **Padj** |  |
| AT1G73330 | DROUGHT-REPRESSED 4 (DR4) | 5.64 | 2.82 E-04 | 6.95 | 6.37 E-05 | 7.22 | 3.34 E-06 | Encodes a plant-specific protease inhibitor-like protein whose transcript level in root is downregulated in response to progressive drought stress. The decrease in transcript level is independent from abscisic acid level. Primarily expressed in the root, especially the root cap. |
| AT1G19410 |  | 3.91 | 1.34 E-02 | 6.2 | 2.91 E-04 | 6.93 | 3.34 E-06 | Affects fecundity under stress, target for epigenetic regulation from TE methylation, F-box associated/Leucine Rich Repeat domains containing protein. |
| AT5G15500 |  | 5.2 | 3.23 E-11 | 3.8 | 1.85 E-04 | 5.75 | 4.18 E-11 | Ankyrin repeat family protein that plays a role in abiotic stress. Functions as an ABA-responsive gene and ABA-mediated stomatal closure. Its transcription is regulated by ABA and JA signaling. |
| AT2G13665 |  | 1.62 | 4.59 E-18 | 3.47 | 1.17 E-71 | 3.37 | 6.70 E-69 | Natural antisense transcript encoding a non-protein coding RNA. Functions in Environmental plasticity in water availability study (Gentoype x Environment expression quantative trait locus interactions). |
| AT5G01300 |  | 2.32 | 2.37 E-04 | 3.07 | 2.54 E-05 | 2.75 | 4.04 E-05 | PEBP (phosphatidylethanolamine-binding protein) family protein. Mutant phenotypes suggest a role in dormancy release and high temperature conditions. |
| AT3G52820 | PURPLE ACID PHOSPHATASE 22 (PAP22) | 2.26 | 2.44 E-04 | 2.97 | 2.46 E-05 | 5.42 | 2.87 E-20 | Purple acid phosphatase 22 plays a critical role in plant's nutrient management and development in response to environmental stress. Downstream of VND7 suggesting a role in xylem differentiation. |
| AT5G59820 | RESPONSIVE TO HIGH LIGHT 41 (RHL41) | 1.71 | 3.47 E-02 | 2.73 | 7.25 E-03 | 2.87 | 5.24 E-04 | C2H2 zinc finger transcription factor that act as a central hub for coordinating a wide range of environmental stresses. |
| AT2G17500 | PIN-LIKES 5 (PILS5) | 3.08 | 1.91 E-04 | 2.67 | 2.63 E-02 | 4.15 | 3.91 E-06 | Auxin efflux transporter located in the endoplasmic reticulum. Intracellular auxin transport facilitator at the endoplasmic reticulum. Auxin transport facilitator which abundance is controlled by ER stress; resulting in protein turnover in a tissue-dependent manner. This process allows for the integration of environmental input with phytochrome auxin signaling; contributing to stress-induced growth adaptation. |
| AT1G04463 |  | 1.8 | 8.82 E-07 | 2.63 | 1.04 E-10 | 4.08 | 4.59 E-28 | Natural antisense transcript modulates activity of AtCAF1g, a deadenylase, removing poly(A) tails and functioning in mRNA decay. The CAF1 family is a regulator of stress responsive genes. |
| AT3G15650 |  | 1.22 | 1.54 E-02 | 2.58 | 2.65 E-06 | 1.22 | 3.18 E-02 | Phospholipase/caborxylesterase involved in cellular homeostasis and response to viral movement in Brassica species |
| AT1G01640 |  | 2.19 | 3.21 E-06 | 2.5 | 6.43 E-06 | 3.38 | 2.94 E-12 | BTB/POZ adaptor protein in CUL3 E3 ubiquitin ligase complexes, contributing to regulated protein degradation pathways that potentially influence flower development and other growth-related processes. |
| AT1G15870 |  | 1.27 | 9.82 E-04 | 2.46 | 2.45 E-09 | 2.51 | 1.15 E-10 | Mitochondrial matrix MAM33‑like protein with no gene‑specific function yet demonstrated in Arabidopsis. |
| AT1G76820 |  | 1.01 | 2.34 E-04 | 2.44 | 6.69 E-17 | 2.39 | 3.63 E-17 | eIF5B/IF2‑like translation‑initiation factor pseudogene candidate. The paralog eIF5B1 (AT1G76810) performs essential roles in heat acclimation. |
| AT5G45150 | RNASE THREE-LIKE PROTEIN 3 (RTL3) | 1.64 | 2.23 E-02 | 2.13 | 2.99 E-02 | 3.44 | 9.08 E-07 | Ribonuclease III–like protein 3 that functions as a double‑stranded RNA (dsRNA)–specific endoribonuclease involved in RNA processing. |
| AT3G16360 | HPT PHOSPHOTRANSMITTER 4 (AHP4) | 3.7 | 7.77 E-15 | 2.06 | 2.27 E-03 | 5.08 | 8.26 E-25 | Histidine phosphotransfer protein that is involved in the cytokinin His‑to‑Asp phosphorelay pathway. shuttles phosphate from cytokinin sensor kinases to response regulators, acting as a redundant positive regulator of cytokinin signaling. |
| AT1G80570 |  | 2.34 | 1.32 E-12 | 1.98 | 8.93 E-07 | 2.46 | 8.63 E-12 | Contains an F-box protein containing leucine-rich repeat domains. By analogy with other F-box/LRR proteins, it is predicted to act as the substrate-recognition component of an SCF (SKP1-CUL1-F-box) E3 ubiquitin ligase and thereby contribute to ubiquitin-dependent protein degradation. |
| AT4G16940 | SIDEKICK SNC1 1 (SIKIC1) | 1.57 | 2.33 E-05 | 1.92 | 2.09 E-05 | 2.12 | 9.24 E-08 | TIR-NBS-LRR protein associated with Arabidopsis immune signaling. Available evidence primarily links it to modulation of SNC1-dependent immune and autoimmune responses, suggesting a role in fine-tuning defense signaling |
| AT1G76530 | PIN-LIKES 4 (PILS4) | 1.67 | 1.12 E-03 | 1.84 | 7.73 E-03 | 2.13 | 1.17 E-04 | ER localized intracellular auxin transporter that regulates auxin homeostasis and limits nuclear auxin signaling, thereby integrating environmental and hormonal signals with plant growth responses. |
| AT5G66960 |  | 2.23 | 7.84 E-09 | 1.82 | 2.89 E-04 | 2.86 | 8.03 E-12 | Mitochondrially localized S9-family (prolyl oligopeptidase) serine peptidase associated with proteolytic peptide processing. |
| AT1G24265 |  | 1.05 | 1.07 E-02 | 1.79 | 2.21 E-04 | 1.32 | 3.21 E-03 | DUF1664 domain protein of unknown function; subcellular localization and biological role remain uncharacterized. |
| AT1G71330 | NON-INTRINSIC ABC PROTEIN 5 (NAP5) | 1.17 | 2.11 E-02 | 1.79 | 5.22 E-03 | 2.69 | 4.57 E-08 | Putative ATP-binding cassette (ABC) transporter involved in ATP-coupled transmembrane transport; specific substrate and physiological role remain uncharacterized. |
| AT5G24280 | GAMMA-IRRADIATION AND MITOMYCIN C INDUCED 1 (GMI1) | 1.79 | 4.07 E-05 | 1.77 | 2.23 E-03 | 3.1 | 4.57 E-12 | SMC hinge domain DNA repair protein required for homologous recombination mediated double strand break repair and maintenance of genome stability. |
| AT1G16030 | HEAT SHOCK PROTEIN 70B (Hsp70b) | 1.1 | 1.03 E-08 | 1.75 | 1.92 E-16 | 2.31 | 4.54 E-31 | Cytosolic Hsp70 chaperone that uses ATP to fold/refold proteins, supporting proteostasis during heat and other stresses. |
| AT1G54575 |  | 2.26 | 7.09 E-05 | 1.74 | 4.41 E-02 | 2.49 | 8.39 E-05 | Small (≈102 aa) unknown protein; strongly predicted nuclear localization and abundant cDNA support, but no defined molecular function or phenotype reported. |
| AT5G42200 | ARABIDOPSIS TOXICOS EN LEVADURA 23 (ATL23) | 1.59 | 2.62 E-05 | 1.66 | 6.25 E-04 | 2.43 | 1.51 E-09 | Plasma‑membrane RING‑type E3 ubiquitin ligase of the ATL family; its precise physiological role in planta remains unclear, though ATL family proteins broadly modulate stress and signaling pathways. |
| AT1G09500 |  | 1.77 | 2.11 E-04 | 1.64 | 1.53 E-02 | 2.83 | 1.38 E-08 | Encodes a Rossmann‑fold oxidoreductase/alcohol dehydrogenase like enzyme, but its substrate(s), pathway context, and in‑planta function are unresolved. |
| AT3G14990 | DJ-1 HOMOLOG A (DJ1A) | 2.07 | 7.27 E-10 | 1.47 | 1.22 E-03 | 2.96 | 5.44 E-16 | DJ‑1/PfpI glyoxalase/deglycase that detoxifies reactive carbonyls and boosts oxidative‑stress tolerance by activating cytosolic SOD1. |
| AT5G59220 | HIGHLY ABA-INDUCED PP2C GENE 1 (HAI1) | 3.23 | 1.45 E-13 | 1.47 | 3.40 E-02 | 2.86 | 5.69 E-09 | HAI1 dampens ABA/osmotic signaling—it is induced under drought yet acts to attenuate aspects of the response. |
| AT2G36800 | DON-GLUCOSYLTRANSFERASE 1 (DOGT1) | 1.08 | 1.60 E-02 | 1.46 | 1.64 E-02 | 1.62 | 7.20 E-04 | Broad-substrate UDP-glycosyltransferase involved in brassinosteroid conjugation and DON detoxification, with wider roles in hormone-related stress responses. |
| AT5G64080 | XYLOGEN PROTEIN 1 (XYP1) | 1.1 | 1.46 E-03 | 1.43 | 1.03 E-03 | 2.81 | 2.03 E-15 | GPI-anchored extracellular lipid-transfer protein (xylogen) that mediates local cell-cell signaling required for xylem differentiation and proper vascular development. |
| AT3G50910 |  | 1.46 | 1.15 E-09 | 1.39 | 2.15 E-06 | 2.11 | 5.55 E-16 | Uncharacterized Arabidopsis protein annotated to the nucleus and detected in protein-interaction networks, with no experimentally established biological role. |
| AT1G20490 |  | 1.88 | 5.80 E-15 | 1.36 | 9.81 E-06 | 1.63 | 2.37 E-09 | 4-coumarate-CoA ligase-like acyl-activating enzyme predicted to convert hydroxycinnamates such as 4-coumarate to CoA esters, providing intermediates for phenylpropanoid metabolism. |
| AT2G31751 |  | 1.62 | 1.29 E-08 | 1.31 | 9.83 E-04 | 1.07 | 1.58 E-03 | Natural antisense lncRNA overlapping the auxin glycosyltransferase gene UGT74D1 potentially regulating that locus. |
| AT4G30460 |  | 2.46 | 1.14 E-09 | 1.29 | 4.80 E-02 | 2.68 | 2.05 E-09 | Encodes a secreted/endomembrane glycine‑rich protein with unknown function. But Plant GRPs as a superfamily are often implicated in stress acclimation, cell‑wall/extracellular functions, and RNA‑related processes. |
| AT4G02390 | POLY(ADP-RIBOSE) POLYMERASE 2 (PARP2) | 1.36 | 2.62 E-04 | 1.23 | 2.24 E-02 | 2.2 | 2.67 E-08 | AT4G02390 is a nuclear poly(ADP‑ribose) polymerase that that is active in base‑excision repair and backup end‑joining pathways. |
| AT3G63010 | GA INSENSITIVE DWARF1B (GID1B) | 1.46 | 1.54 E-05 | 1.21 | 1.29 E-02 | 1.58 | 2.14 E-05 | High-affinity gibberellin receptor that binds bioactive GA, promotes DELLA repressor degradation, and activates GA-dependent growth and developmental programs. |
| AT5G46640 | AT-HOOK MOTIF NUCLEAR LOCALIZED PROTEIN 8 (AHL8) | 1.41 | 8.64 E-09 | 1.19 | 1.97 E-04 | 1.76 | 4.15 E-11 | AHL8 is a nuclear AT‑hook protein predicted to bind AT‑rich DNA; it is likely a transcription‑related regulator, though gene‑specific biological functions remain to be conclusively defined. |
| AT3G13380 | BRI1-LIKE 3 (BRL3) | 1.46 | 1.17 E-08 | 1.02 | 4.38 E-03 | 1.26 | 1.57 E-05 | Along with BRL1, BRL3 acts as a brassinosteroid receptor specialized for vascular tissues, contributing to cell growth and vascular differentiation. |

**B. Down regulated genes (54)**

| **54 Down Regulated Genes** | | | | | | | | **Description** |
| --- | --- | --- | --- | --- | --- | --- | --- | --- |
| **Genes** | | **Col-O salt** | | **SST Ctrl** | | **SST salt** | |  |
| **ID** | **Name** | **Log_2_FC** | **Padj** | **Log_2_FC** | **Padj** | **Log_2_FC** | **Padj** |  |
| AT5G41660 |  | -7.93 | 1.09 E-11 | -9.32 | 7.79 E-08 | -9.65 | 3.08 E-09 | Endogenous target of histone deacetylase (HDA6). Plays a role in epigenetic gene silencing. |
| AT3G48290 | CYTOCHROME P450, FAMILY 71, SUBFAMILY A, POLYPEPTIDE 24 (CYP71A24) | -1.06 | 4.01 E-02 | -8.37 | 3.00 E-06 | -8.7 | 1.36 E-07 | Members of the CYP71A gene family are known to be regulated by abiotic stress. |
| AT3G29631 |  | -1.23 | 1.95 E-02 | -8.13 | 6.98 E-06 | -8.47 | 3.22 E-07 | GTPase-activating protein. Nuclear and cytoplasmic located. Nuclear localization is a critical Positive regulator of flowering transition in Arabidopsis. The G3BP family are nucleators involved in the formation of stress granules to regulate mRNA stability. |
| AT3G42658 | SADHU NON-CODING RETROTRANSPOSON 3-2 (SADHU3-2) | -3.53 | 1.87 E-02 | -7.2 | 7.73 E-03 | -7.54 | 6.84 E-04 | Member of Sadhu non-coding retrotransposon family. Endogenous target of epigenetic silencing by HDA6 deacetylase implying epigenetic functions. In some natural accessions the allele is methylated and silenced. |
| AT3G48080 |  | -2.76 | 1.19 E-11 | -6.11 | 3.17 E-13 | -10.2 | 2.15 E-10 | Functional homolog of EDS1; mediates defense response and forms regulatory hub that coordinates Programmed Cell Death and accumulation of salicylic acid. Structurally a N-terminal coiled-coil domain and C-terminal domain that shares structural similarities with pore-forming toxins founds in mammals and fungi suggesting that when activated it may help form pores in the plasma membrane to trigger defense response or localized cell death. |
| AT3G50480 | HOMOLOG OF RPW8 4 (HR4) | -1.19 | 1.90 E-03 | -5.63 | 2.28 E-21 | -8.66 | 8.50 E-14 | Plant immune responses. Structurally a N-terminal coiled-coil domain and C-terminal domain that shares structural similarities with pore-forming toxins founds in mammals and fungi suggesting that when activated it may help form pores in the plasma membrane to trigger defense response or localized cell death. |
| AT3G47010 |  | -1.3 | 1.81 E-04 | -4.52 | 2.29 E-11 | -2.91 | 7.98 E-12 | Hydrolysis of complex sugars may be involved in plant's response to high CO2 levels. |
| AT3G47380 | (ATPMEI11) | -2.2 | 2.76 E-10 | -3.61 | 2.96 E-13 | -5 | 4.52 E-18 | Pectin methylesterase inhibitor that regulates stability of the plant cell wall. Highly induced by necrotrophic pathogens (e.g., *Botrytis cinera*). May be involved in salt and drought response as well as freezing tolerance. |
| AT5G44568 | (STMP10) | -2.61 | 1.19 E-07 | -3.43 | 2.50 E-08 | -4.42 | 3.18 E-13 | Secreted peptide which functions in plant growth and pathogen defense. Also plays a role in root development and gravitropism. |
| AT3G45430 | L-TYPE LECTIN RECEPTOR KINASE I.5 (LECRK-I.5) | -1.58 | 4.81 E-04 | -3.22 | 4.84 E-07 | -3.81 | 4.75 E-10 | Extracellular ATP transmembrane receptor involved in innate immunity. Implicated in pathogen tolerance and sensing extracellular ATP. |
| AT5G52940 | DUF295 ORGANELLAR B 5 (ATDOB5) | -1.74 | 5.57 E-03 | -3.17 | 1.73 E-03 | -7.77 | 5.00 E-06 | Highly expressed mitochondrial DUF295 protein of unknown function |
| AT5G57760 |  | -3.35 | 1.35 E-05 | -3.07 | 7.71 E-03 | -6.15 | 3.42 E-04 | Small nuclear protein of unknown molecular and biological function. |
| AT1G24145 |  | -1.28 | 3.06 E-02 | -2.84 | 3.88 E-04 | -7.32 | 4.85 E-06 | Small protein that when cleaved by subtilases to produce SERINE RICH ENDOGENOUS PEPTIDES (SCOOP) proteins that are involved in signaling |
| AT1G31710 | COPPER AMINE OXIDASE ALPHA 3 (CuAOalpha3) | -1.62 | 9.53 E-03 | -2.81 | 5.13 E-04 | -1.48 | 4.12 E-02 | Extracellular copper amine oxidase that catalyzes the oxidative deamination of primary amines, functioning in amine metabolism and redox‑related processes, and contributing to hydrogen peroxide production involved in plant metabolism and signaling. |
| AT5G45670 |  | -4.18 | 1.29 E-12 | -2.79 | 1.30 E-04 | -3.7 | 3.48 E-08 | Encodes a GDSL-like lipase/acylhydrolase family protein containing the conserved catalytic GDSL motif and is predicted to function as a hydrolase acting on ester bonds in lipid metabolic processes. |
| AT4G32280 | INDOLE-3-ACETIC ACID INDUCIBLE 29 (IAA29) | -2 | 1.12 E-04 | -2.51 | 2.87 E-04 | -3.6 | 2.71 E-08 | Member of the Aux/IAA transcriptional repressor family that are involved in the regulation of early auxin‑responsive gene expression and developmental signaling. |
| AT3G46880 |  | -3.74 | 2.03 E-06 | -2.33 | 3.74 E-02 | -3.93 | 5.15 E-05 | Uncharacterized gene that is expressed across many tissues and developmental stages with a predicted localization in the chloroplast; suggesting that it is involved in an as‑yet‑unknown aspect of plastid function. |
| AT1G01620 | PLASMA MEMBRANE INTRINSIC PROTEIN 1C (PIP1C) | -2.11 | 1.23 E-27 | -2.3 | 5.91 E-25 | -3 | 2.58 E-42 | Encodes PIP1;3 a member of the plasma membrane intrinsic protein (PIP) aquaporin family, that are essential for water transport, growth, and reproductive physiology. |
| AT1G02230 |  | -4.82 | 1.08 E-08 | -2.13 | 2.78 E-02 | -3.84 | 9.46 E-06 | NAC‑family transcription factor that localizes to the nucleus and regulates gene expression involved in multicellular development and general transcriptional programs in Arabidopsis. |
| AT1G72970 | HOTHEAD (HTH) | -1.53 | 6.97 E-10 | -2.02 | 7.21 E-12 | -1.78 | 1.84 E-10 | FAD‑dependent oxidoreductase is essential for cuticle formation and proper organ separation during flower development. |
| AT1G02450 | NIM1-INTERACTING 1 (NIMIN1) | -3.25 | 3.16 E-09 | -1.98 | 1.01 E-02 | -5.83 | 1.25 E-09 | Nuclear adaptor protein that modulates NPR1‑dependent salicylic‑acid signaling and fine‑tunes NPR1-dependent systemic acquired resistance. |
| AT1G13750 | PURPLE ACID PHOSPHATASE 1 (PAP1) | -1.08 | 1.01 E-04 | -1.93 | 4.52 E-09 | -3.03 | 1.83 E-21 | Encodes a purple acid phosphatase family protein whose expression is expressed broadly in the cell walls and the apoplast |
| AT4G15680 | (GRXS4) | -1.2 | 9.45 E-03 | -1.76 | 4.36 E-03 | -1.76 | 5.66 E-04 | Thioredoxin‑superfamily redox protein that acts downstream of cytokinin signaling to negatively regulates primary root growth in response to nitrate sufficient conditions. |
| AT4G19380 |  | -1.17 | 4.19 E-02 | -1.75 | 3.92 E-02 | -1.46 | 2.42 E-02 | Encodes an FAD‑dependent long‑chain fatty alcohol dehydrogenase, likely functioning in lipid and wax‑related metabolism, though its in‑vivo physiological role remains incompletely resolved. |
| AT1G23830 |  | -1.48 | 4.17 E-03 | -1.71 | 2.33 E-02 | -2.82 | 2.49 E-06 | Encodes an uncharacterized, plant‑specific multi‑pass membrane protein. Its molecular and biological functions remain unknown. |
| AT1G29450 | SMALL AUXIN UPREGULATED RNA 64 (SAUR64) | -2.87 | 1.35 E-12 | -1.64 | 2.70 E-03 | -2.97 | 8.58 E-10 | (Small Auxin Up RNA) SAUR‑family auxin‑responsive protein likely involved in early auxin‑mediated growth regulation. |
| AT5G44020 |  | -3.61 | 1.91 E-89 | -1.64 | 7.69 E-15 | -7.2 | 7.54 E-136 | Encodes a HAD family IIIB acid phosphatase. The protein is associated with vacuolar, extracellular/cell-wall, and plasmodesmata compartments, while its predicted enzymatic activity suggests possible roles in phosphate remobilization or scavenging, a specific in vivo physiological function has not yet been experimentally established. |
| AT5G19120 |  | -1.2 | 1.50 E-03 | -1.63 | 1.77 E-03 | -1.03 | 1.96 E-02 | Pepsin‑like aspartic protease involved in proteolysis, likely functioning in the endomembrane or extracellular compartments. |
| AT3G15540 | INDOLE-3-ACETIC ACID INDUCIBLE 19 (IAA19) | -2.31 | 9.37 E-07 | -1.61 | 1.97 E-02 | -2.19 | 5.83 E-05 | Nuclear Aux/IAA repressor that modulates auxin‑dependent growth and tropic responses. |
| AT3G52430 | PHYTOALEXIN DEFICIENT 4 (PAD4) | -1.88 | 6.99 E-19 | -1.55 | 2.37 E-09 | -2.77 | 4.34 E-27 | Regulator of Immune response that promotes salicylic‑acid‑dependent defense and systemic acquired resistance. |
| AT1G16260 | WALL ASSOCIATED KINASE LIKE 8 (WAKL8) | -1.22 | 6.29 E-11 | -1.43 | 1.75 E-10 | -3.02 | 9.02 E-42 | Encodes a wall‑associated receptor kinase that catalyzes protein phosphorylation, indicating a role in signal transduction. But a specific physiological function has not yet been experimentally established. |
| AT1G22650 | ALKALINE/NEUTRAL INVERTASE D (A/N-InvD) | -2.03 | 2.92 E-13 | -1.39 | 1.22 E-04 | -4.24 | 1.79 E-31 | Encodes a member of the cytosolic neutral/alkaline invertase that catabolizes sucrose to glucose and fructose, contributing to sucrose metabolism/carbon partitioning in diverse tissues. |
| AT4G15660 | (GRXS8) | -1.11 | 9.89 E-03 | -1.32 | 3.55 E-02 | -2.45 | 3.07 E-07 | CC‑type glutaredoxin induced by nitrate; acts downstream of cytokinin to negatively regulate primary root growth in response to nitrate. |
| AT5G02760 | ARABIDOPSIS PP2C CLADE D 7 (APD7) | -2.97 | 6.51 E-15 | -1.32 | 1.81 E-02 | -4.59 | 2.24 E-20 | Encodes a (protein phosphatase 2C) PP2C‑type phosphatase that suppresses premature leaf senescence. |
| AT1G53920 | GDSL-MOTIF LIPASE 5 (GLIP5) | -1.61 | 6.62 E-05 | -1.32 | 3.26 E-02 | -2.22 | 2.86 E-06 | Extracellular GDSL esterase/lipase involved in lipid metabolism. The GDSL lipase family is a large, multifunctional family involved in lipid metabolism, stress/immune responses, and cell‑wall/barrier remodeling. |
| AT1G14280 | PHYTOCHROME KINASE SUBSTRATE 2 (PKS2) | -1.4 | 2.22 E-13 | -1.29 | 2.97 E-08 | -2.24 | 1.81 E-24 | Encodes a photoreceptor‑associated signaling protein involved in blue‑light‑mediated phototropism. |
| AT5G66740 | BOUNDARY OF ROP DOMAIN8 (BDR8) | -1.33 | 8.59 E-04 | -1.29 | 3.28 E-02 | -3.27 | 9.44 E-11 | Part of the BDR (Boundary of ROP Domain) family that is strongly expressed in pollen and pollen tubes. It is predicted to function in maintaining cell polarity during pollen tube growth bases off of other BDR functions. |
| AT1G29500 | SMALL AUXIN UPREGULATED RNA 66 (SAUR66) | -2.14 | 7.95 E-10 | -1.22 | 1.69 E-02 | -2.24 | 1.08 E-07 | Encodes a member of the auxin‑responsive regulatory family that functions in the auxin signaling pathway and is believed to similarly promote auxin‑dependent cell and organ elongation during plant growth. |
| AT1G65490 | (STMP5) | -1.4 | 8.48 E-06 | -1.19 | 7.56 E-03 | -2.25 | 4.81 E-10 | Encodes a small, secreted precursor protein that is processed into a (Serine‑rich Endogenous Peptide) SCOOP signaling peptides, which activate activates immunity and modulates growth. |
| AT5G25460 | DUF642 L-GALL RESPONSIVE GENE 2 (DGR2) | -1.28 | 3.80 E-12 | -1.19 | 1.14 E-07 | -1.43 | 5.32 E-12 | DUF642 cell‑wall protein localized to the apoplast that modulates root development and cell‑wall (pectin) dynamics, is specifically induced by L‑GalL resulting in the shorter roots and smaller rosettes. |
| AT5G15580 | LONGIFOLIA1 (LNG1) | -1.15 | 1.98 E-05 | -1.18 | 9.99 E-04 | -1.14 | 2.01 E-04 | TRM‑family protein that promotes longitudinal (polar) cell elongation to control organ shape (leaves, petals, siliques) |
| AT5G41740 |  | -1.22 | 1.85 E-04 | -1.14 | 1.58 E-02 | -2.3 | 5.15 E-10 | TIR–NBS–LRR class disease resistance protein that functions as an intracellular immune receptor. It participates in pathogen recognition and activation of innate immune signaling. |
| AT3G56710 | SIGMA FACTOR BINDING PROTEIN 1 (SIB1) | -1.5 | 1.26 E-13 | -1.14 | 8.66 E-06 | -2.65 | 3.38 E-30 | VQ‑motif protein that functions as a positive regulator of plant immunity by co‑activating the WRKY33 transcription factor, integrating SA/JA hormone signaling and enhancing resistance to pathogens. |
| AT5G24530 | DOWNY MILDEW RESISTANT 6 (DMR6) | -1.44 | 9.46 E-06 | -1.13 | 1.64 E-02 | -2.23 | 5.89 E-10 | Salicylic acid 5‑hydroxylase that negatively regulates plant defense by catabolizing (salicylic acid) SA, thereby limiting immune activation and maintaining growth–defense balance. |
| AT1G68840 | RELATED TO ABI3/VP1 2 (RAV2), TEM2 (TEMPRANILLO 2) | -1.58 | 1.94 E-06 | -1.12 | 2.48 E-02 | -2.83 | 1.10 E-12 | Nuclear AP2/ERF–B3 transcription factor that acts to repress (FLOWERING LOCUS T) FT and delaying flowering as well as participating in ethylene‑ and stress‑responsive gene regulation, thereby linking developmental timing with environmental signaling pathways. |
| AT1G24147 |  | -1.9 | 1.23 E-16 | -1.11 | 2.66 E-04 | -3.97 | 7.89 E-47 | Encodes a small, secreted precursor protein that is proteolytically processed into SERINE‑RICH ENDOGENOUS PEPTIDES (SCOOPs). Members of the SCOOP peptide family function as plant signaling peptides involved in immune and stress responses. |
| AT5G02890 |  | -2.9 | 6.09 E-17 | -1.11 | 4.32 E-02 | -2.64 | 2.27 E-11 | ER‑associated HXXXD‑type acyl‑transferase that modulates cuticular wax biosynthesis by limiting very‑long‑chain fatty acid elongation beyond 28 carbons. Its activity shapes stem wax composition, influences cuticle properties, and indirectly affects phytohormone homeostasis, linking lipid metabolism with developmental and stress‑response pathways. |
| AT1G69160 | BIG GRAIN  LIKE 1 (BGL1) | -1.3 | 2.51 E-06 | -1.1 | 4.28 E-03 | -1.52 | 1.48 E-06 | Nuclear regulator involved in auxin transport and auxin‑activated signaling pathways. Through its role in modulating auxin distribution and response, BIG1E contributes to plant growth and developmental regulation. |
| AT3G06070 |  | -2.42 | 7.22 E-29 | -1.1 | 5.43 E-05 | -1.98 | 1.81 E-15 | Small, broadly expressed nuclear‑localized protein, Its sequence composition and nuclear localization suggest a possible regulatory function, but this remains speculative. |
| AT3G04720 | PATHOGENESIS-RELATED 4 (PR4) | -2.88 | 8.06 E-17 | -1.1 | 4.24 E-02 | -3.81 | 9.82 E-21 | Component of the plant innate immune system, functioning primarily as an extracellular antimicrobial defense protein induced during biotic stress. |
| AT5G22390 |  | -2.1 | 1.12 E-20 | -1.09 | 2.45 E-04 | -2.63 | 7.42 E-24 | Nuclear protein of unknown function containing the DUF3049 conserved domain. |
| AT1G29430 | SMALL AUXIN UPREGULATED RNA 62 (SAUR62) | -1.98 | 4.33 E-11 | -1.07 | 1.58 E-02 | -2.56 | 1.89 E-12 | SAUR‑family auxin‑responsive gene implicated in auxin‑mediated growth during reproduction, with pollination/pollen tube‑associated expression supporting pollen tube development/function. |
| AT5G54610 | ANKYRIN  (ANK) | -2.86 | 1.85 E-33 | -1.06 | 1.22 E-03 | -3.36 | 3.23 E-35 | Plasma‑membrane ankyrin‑repeat protein that acts as a positive regulator of plant immunity, functioning downstream of the receptor‑like protein SNC2 to promote defense signaling and pathogen resistance. |
| AT5G48900 | (ATPLL21) | -1.31 | 7.72 E-16 | -1.01 | 5.82 E-07 | -1.4 | 2.79 E-14 | Ca²⁺‑dependent pectate lyase that is involved in pectin catabolism. The pectate lyase family functions as part of cell wall remodeling and catabolism, supporting growth‑related wall dynamics. |

**Supplementary Table 6.** Detailed components and function of the first synthetic desalination genetic circuit (osSST1, lab designation, TKK740) in rice. These components were assembled without refactoring and as such, a bioinformatics analysis suggests they could retain various endogenous regulations (e.g., cryptic splice sites, microRNA regulatory sites).

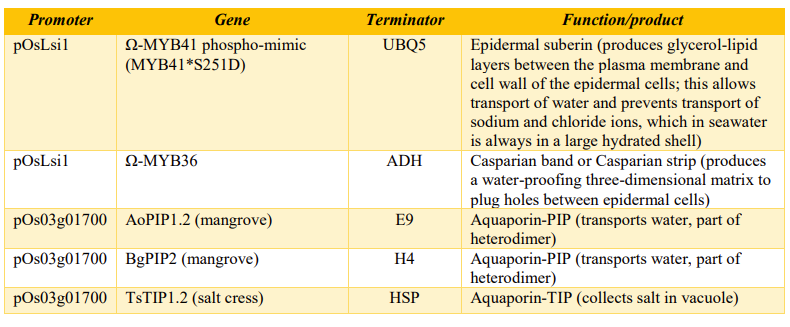

**Supplementary Table 7. List of clones used in this paper.** Genetic Component, Module, or Circuit Name refers to the name used in this publication or in our laboratory. The list is in the order in which the specific component, module or circuit is described in the text.

| **Genetic Component, Module or Circuit Name** | **Description** | **Genes** |
| --- | --- | --- |
| **TKK670** | ABCG37 promoter screening | ABCG37::mCherry |
| **TKK633** | ABCG37 promoter screening (table S1) | ABCG37::AoPIP2.2_mCherry |
| **SGP110** | KC1 promoter screening (table S1) | KC1::mCherry |
| **SGP106** | CESA7 promoter screening (table S1) | CESA7::GFP |
| **SGP107** | XCP1 promoter screening (table S1) | XCP1::GFP |
| **TKK649** | Casparian strip in root epidermis (Fig. 2A) | ABCG37::omega_MYB36_TNOS |
| **SRB2-1 or KJM325** | Suberin in root epidermis (Fig. 2B) | ABCG37::MYB41^S251D^_UBQ5 |
| **KJM326** | GPAT5 marker for suberin biosynthesis (Fig. 2B) | GPAT5::mCherry_SYP122 |
| **KJM725** | Marker for suberin biosynthesis, and suberin in root epidermis (Figs. 2B-D, and fig S1, S2) | **KJM325** pCambia Kan ABCG37::MYB41^S251D^; **cotransformed with**  **KJM326** pCambia BASTA GPAT5::AtmCherry-Syp122 |
| **TKK733** | Filter-barrier line with both suberin and Casparian strip in root epidermis (Fig. 3A) | **TKK730** ABCG37::omega_MYB36; **cotransformed with KJM325** ABCG37::MYB41^S251D^_UBQ5 |
| **TKK734** | Filter-barrier line with both suberin and Casparian strip in root epidermis, and aquaporins (Fig. 3A) | **TKK731** ABCG37::omega_MYB36 + KC1::TsTIP1.2; **cotransformed with**  **TKK732** ABCG37::MYB41^S251D^_UBQ5 + KC1::AoPIP1.2 + KC1::BgPIP2 |
| **TKK630** | Individual aquaporin functional screens (fig. S7 and S8) | ABCG37::AoPIP1.2_mCherry |
| **TKK632** | Individual aquaporin functional screens (fig. S7 and S58 | ABCG37::BgPIP2_mCherry |
| **TKK669** | Individual aquaporin functional screens (fig. S7 and S8) | ABCG37::TsTIP1.2_GFP |
| **AQP1 or SGP105** | Mannitol osmoticum (Fig. S3, S10) | XCP1::M6PR + CESA7::AgMaT2 |
| **AGP2 or JTP001** | Mannitol osmoticum plus omega enhancer (fig. S10) | XCP1::omega_M6PR + CESA7::AgMaT2 |
| **TKK777** | Aquaporin expression in root epidermis/cortex (fig. S11) | KC1::AoPIP1.2_mCherry |
| **TKK778** | PIP expression in root epidermis/cortex (fig. S11) | KC1::BgPIP2_mCherry |
| **TKK779** | TIP expression in root epidermis/cortex (fig. S11) | KC1::TsTIP1.2_GFP |
| **SST1 or TKK716** | Two T-DNA version of complete circuit (Figs. 4A) | **TKK711** ABCG37::MYB41^S251D^_UBQ5 + KC1::AoPIP1.2_E9 + KC1::BgPIP2_H4 + XCP1::omega_M6PR_TNOS; **cotransformed with** **TKK710** ABCG37::omega_MYB36_ADH + KC1::TsTIP1.2_HSP + CESA7::AgMaT2_rbsC |
| **SST2 or**  **TKK717** | One T-DNA version of complete circuit (Figs. 4B-C, Fig. 5, Fig. 6, fig. S1,fig S3G, fig S13,fig. S14, Tables S1, S4, S5, S6, S15, S16) | ABCG37::MYB41^S251D^_UBQ5 + KC1::AoPIP1.2_E9 + KC1::BgPIP2_H4 + XCP1::omega_M6PR_TNOS + ABCG37::omega_MYB36_ADH + KC1::TsTIP1.2_HSP + CESA7::AgMaT2_rbsC |

**SUPPLEMENTARY FIGURES**

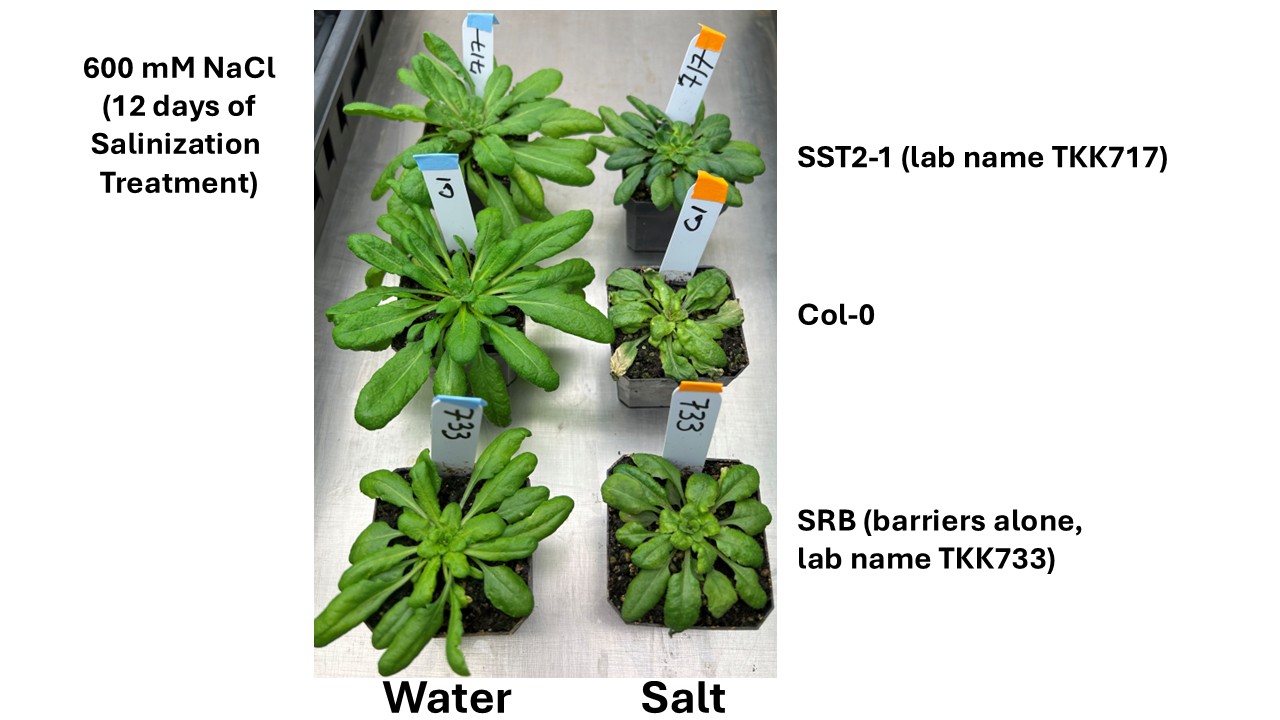
**A.**

**B.**

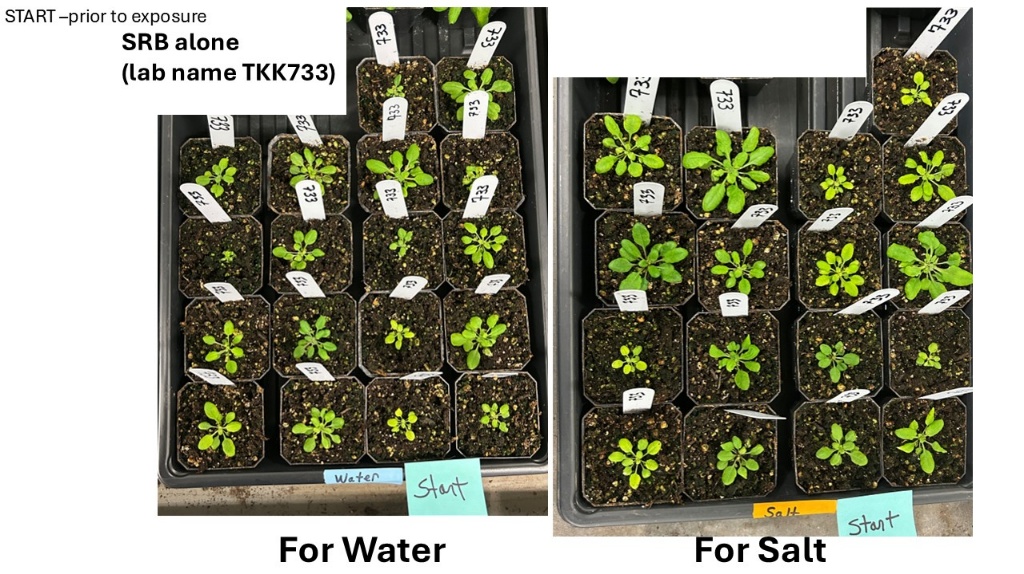

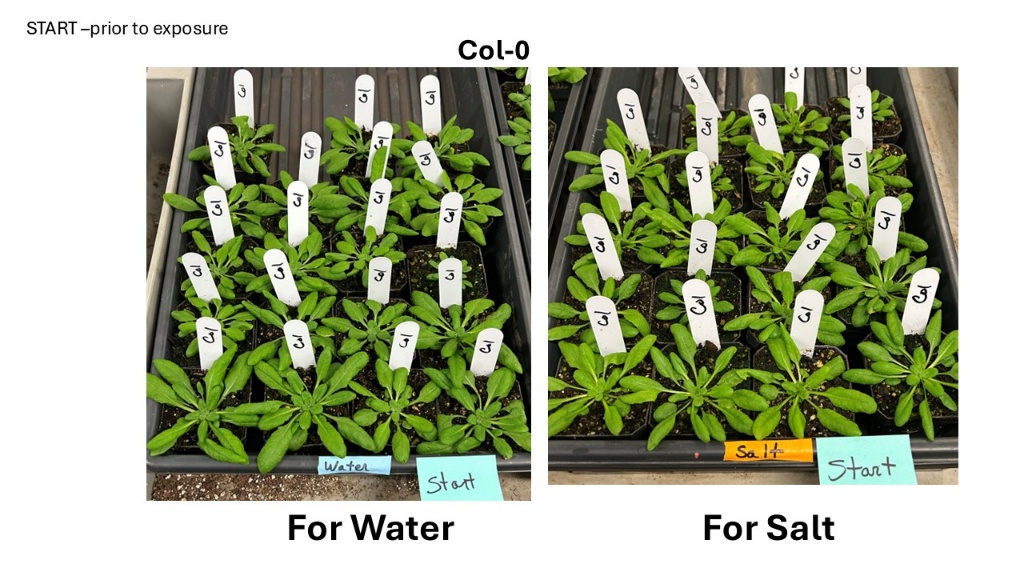

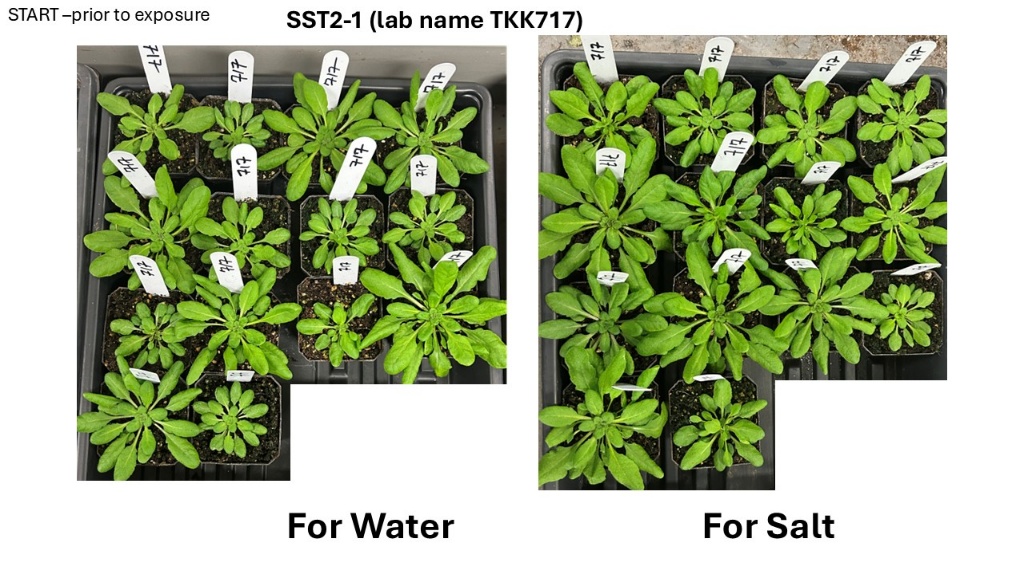

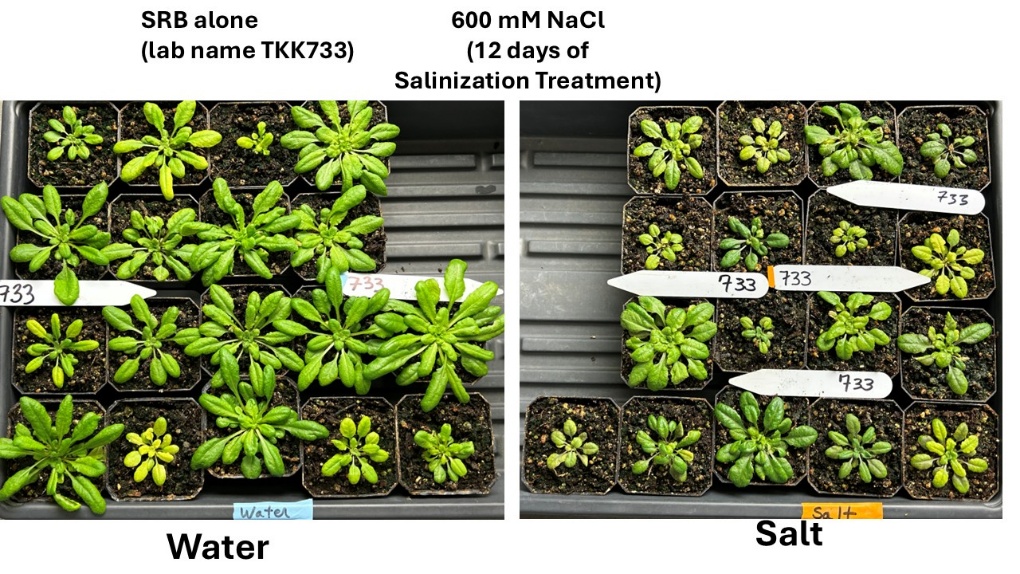
**C.**

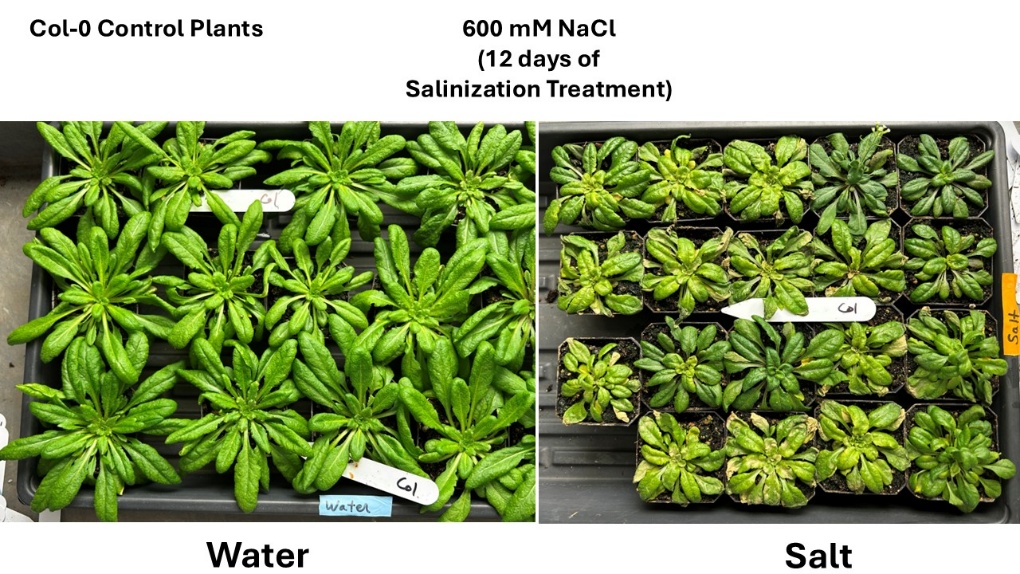

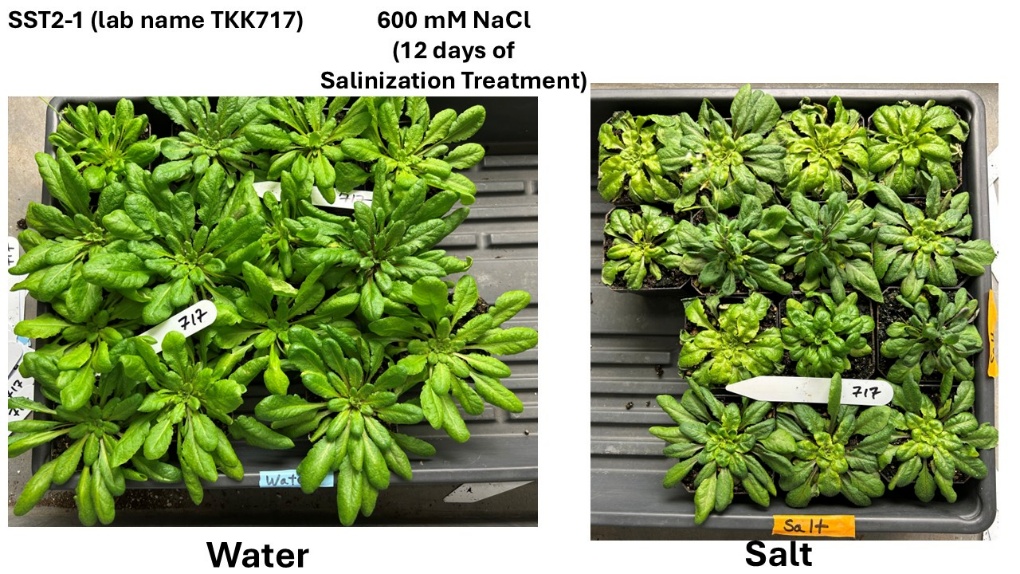
**Supplementary Fig. 1 Under low light conditions, SRB plants show variable stunting yet high salt tolerance**. **A.** Rapid salinization treatment to 600 mM NaCl (12 days total). Left, plants grown in short days and treated normally with tap water. Right, plants grown in short days and treated with 600 mM NaCl show salt tolerance. **B**. Diverse stunting and chlorosis in groups of SRB plants and images of all plants prior to salt treatment (next page). **C.** SRB salt treatment and salt treatment of all plants with 600 mM NaCl (12 days total salinization).

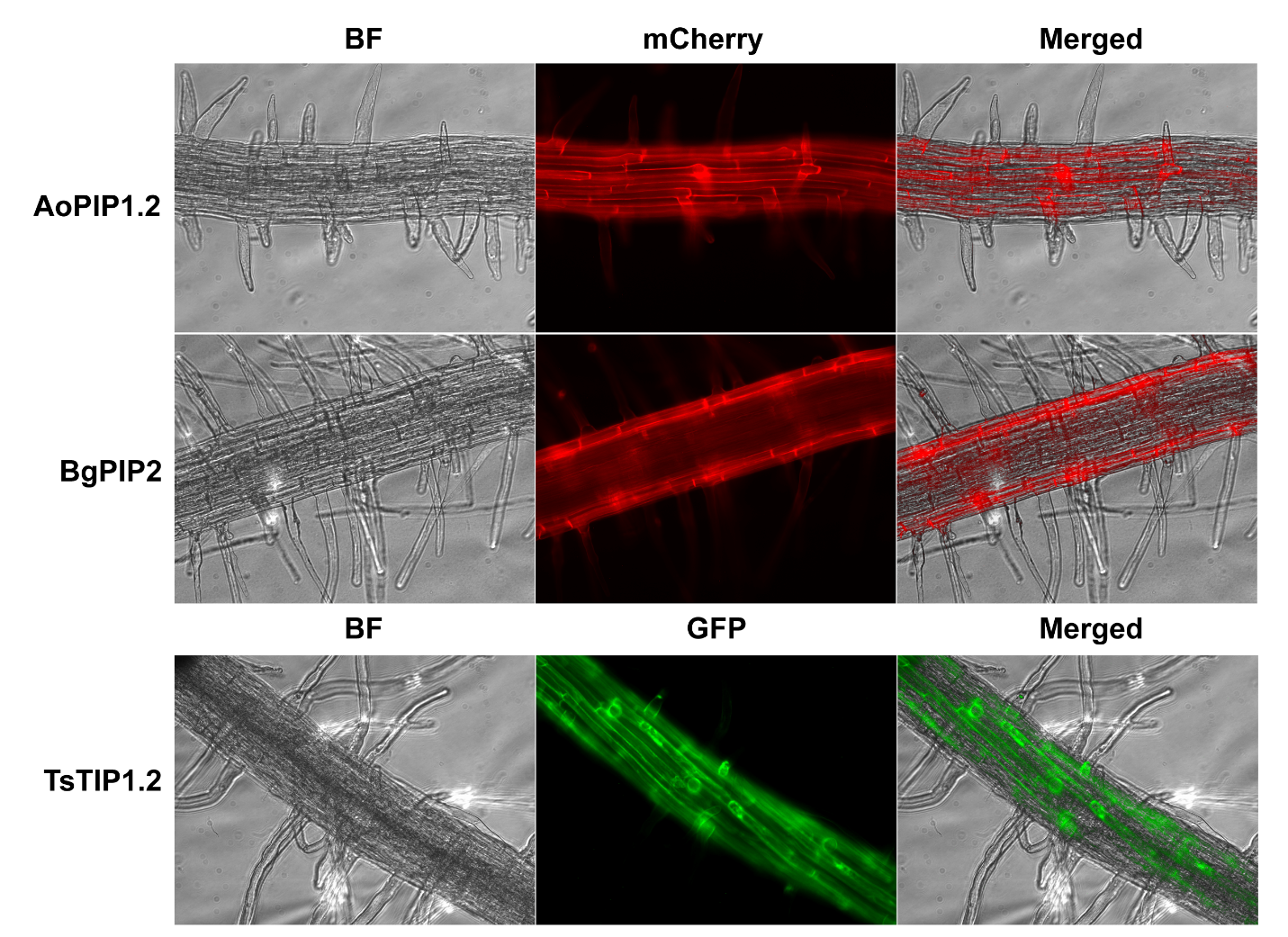

**Supplementary Fig. 2. Aquaporin-reporter gene fusions demonstrate that mangrove and Thellungiella aquaporins are stable in Arabidopsis roots.** Aquaporins from mangroves and Thellungiella used in the genetic circuits were individually fused in frame (C-terminal) to mCherry or GFP reporter proteins and expressed in Arabidopsis plants using the ABCG37 promoter. AoPIP1.2, Ao = *Avicennia officinalis*; BgPIP2, Bg = *Bruguiera gymnorhiza*; TsTIP1.2, Ts = *Thellungiella salsuginea*. Only the aquaporins used in the AQP and SST genetic circuits described in the text are shown.

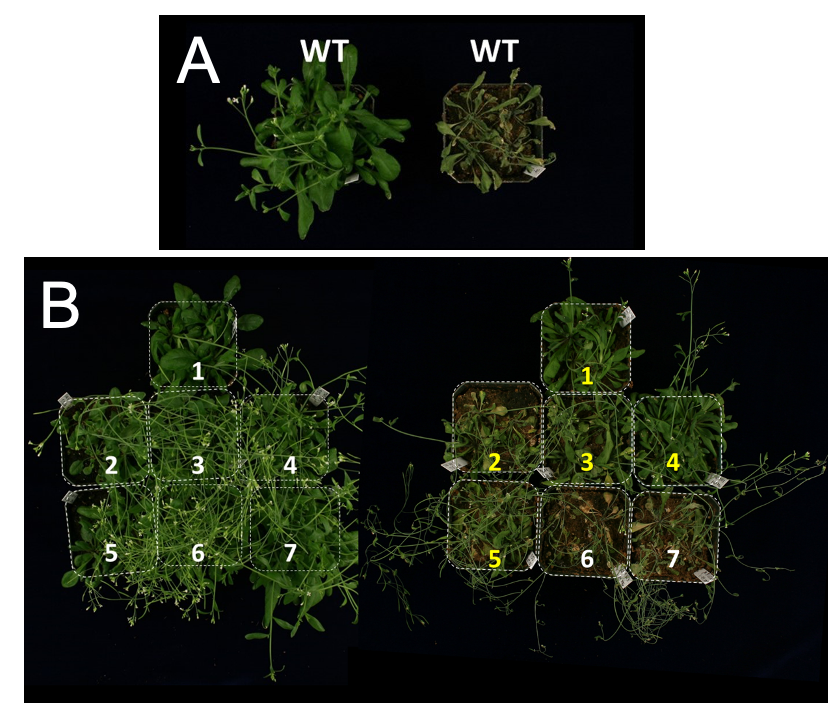

**Supplementary Fig. 3.** **Drought stress assay on Arabidopsis plants expressing the mangrove aquaporin AoPIP1.2.** (**A**) Wildtype (WT) plants, watered daily with tap water (left) and not watered (right) for 10 days. (**B**) Transgenic Arabidopsis plants expressing ABCG37::AoPIP1.2-mCherry (TKK630). Transgenic plants watered daily with tap water for 10 days (left) or not watered for 10 days (right). Numbers indicate independent lines. Yellow numbers denote the independent transgenic lines showing drought tolerance, indicating that the AoPIP1.2 is functional in Arabidopsis. (Dotted lines approximate edges of pots. Pot weights were not recorded.

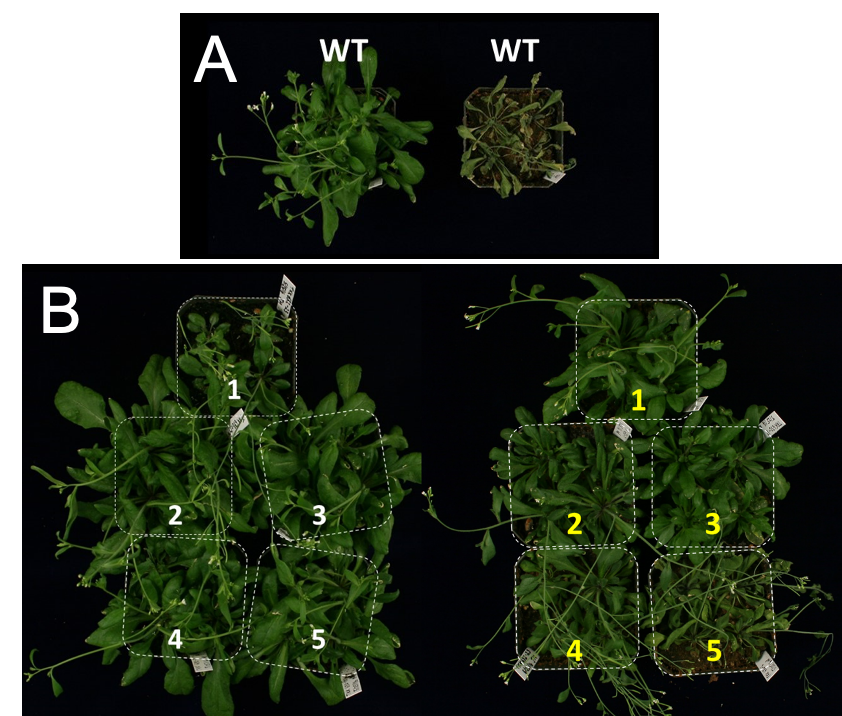

**Supplementary Fig. 4. Drought stress assay on Arabidopsis plants expressing the mangrove aquaporin BgPIP2**. (**A**) Wild type (WT) plants, watered daily with tap water (left) and not watered (right) for 10 days. (**B**) Transgenic plants ABCG37:: BgPIP2-mCherry (TKK632) Transgenic plants watered daily with tap water for 10 days (left) or not watered for 10 days (right). Numbers indicate independent lines. Yellow numbers denote the best transgenic lines showing drought tolerance indicating that BgPIP2 functions in Arabidopsis. Dotted lines approximate the edges of the pots. Pot weights were not recorded.

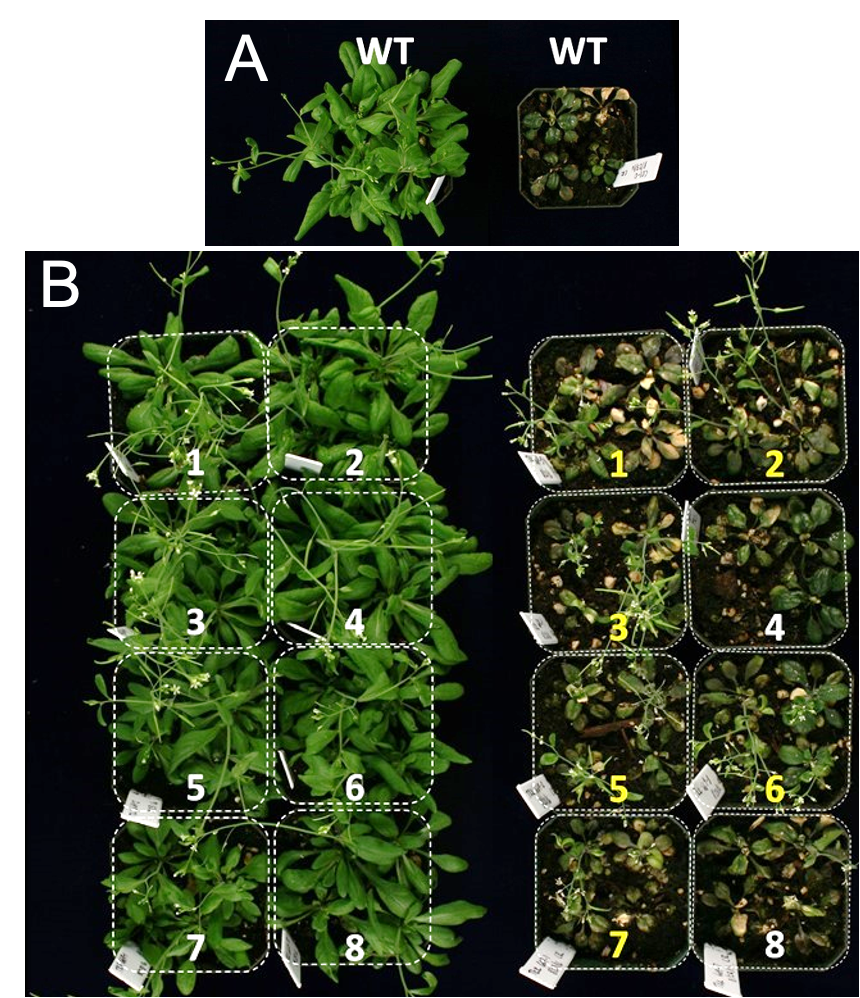

**Supplementary Fig. 5.** **Salt tolerance assay on Arabidopsis expressing aquaporin TsTIP1.2.** (**A**) Wild type (WT) plants, watered with tap water (left) and watered with 350 mM NaCl (right) for 12 days. (**B**) Arabidopsis plants expressing ABCG37::TsTIP1.2-GFP (TKK669). Transgenic plants watered with tap water for 12 days (left) or watered with 350 mM NaCl for 12 days (right). Numbers indicate independent lines. Yellow numbers denote the transgenic lines showing salt tolerance by the ability to form siliques (seed pods), thus indicating TsTIP1.2 function. Sixteen of 31 surviving ABCG37::TsTIP1.2-GFP transgenic plants bolted and seven of these were fertile (set seed) with watering with 350 mM NaCl. In contrast, wildtype plants did not show further growth, and none bolted or set seed.

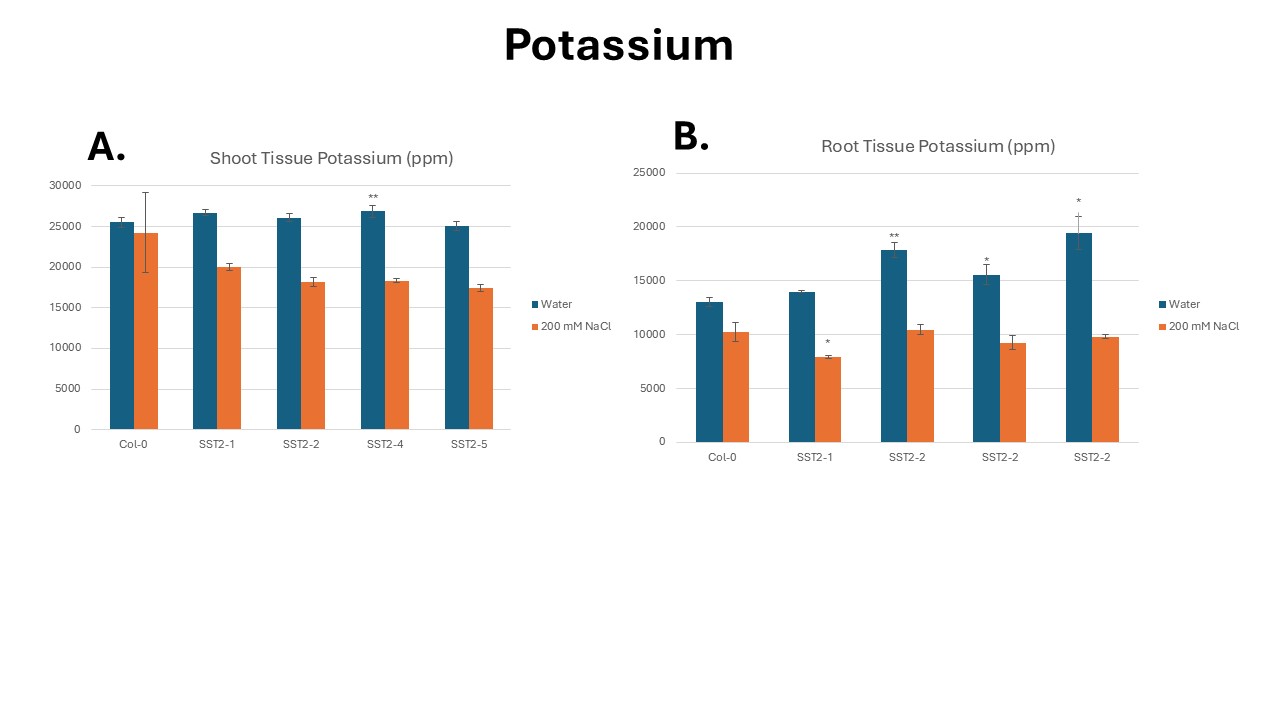

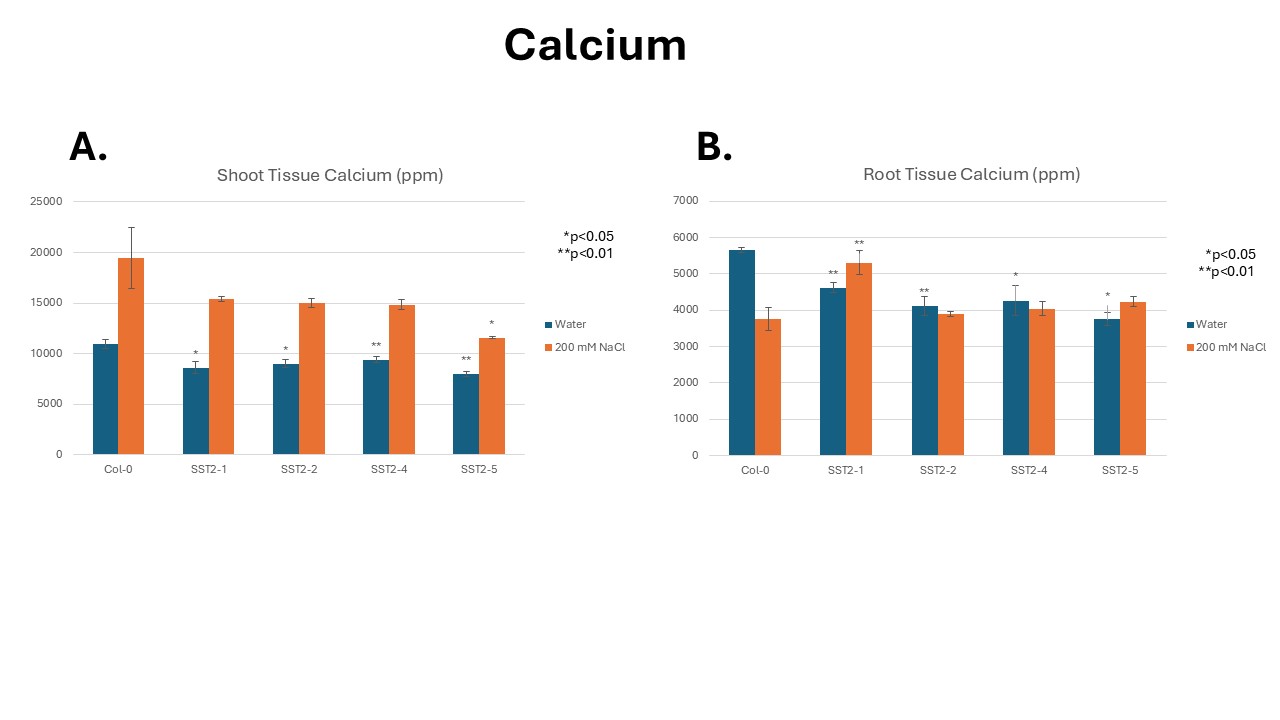

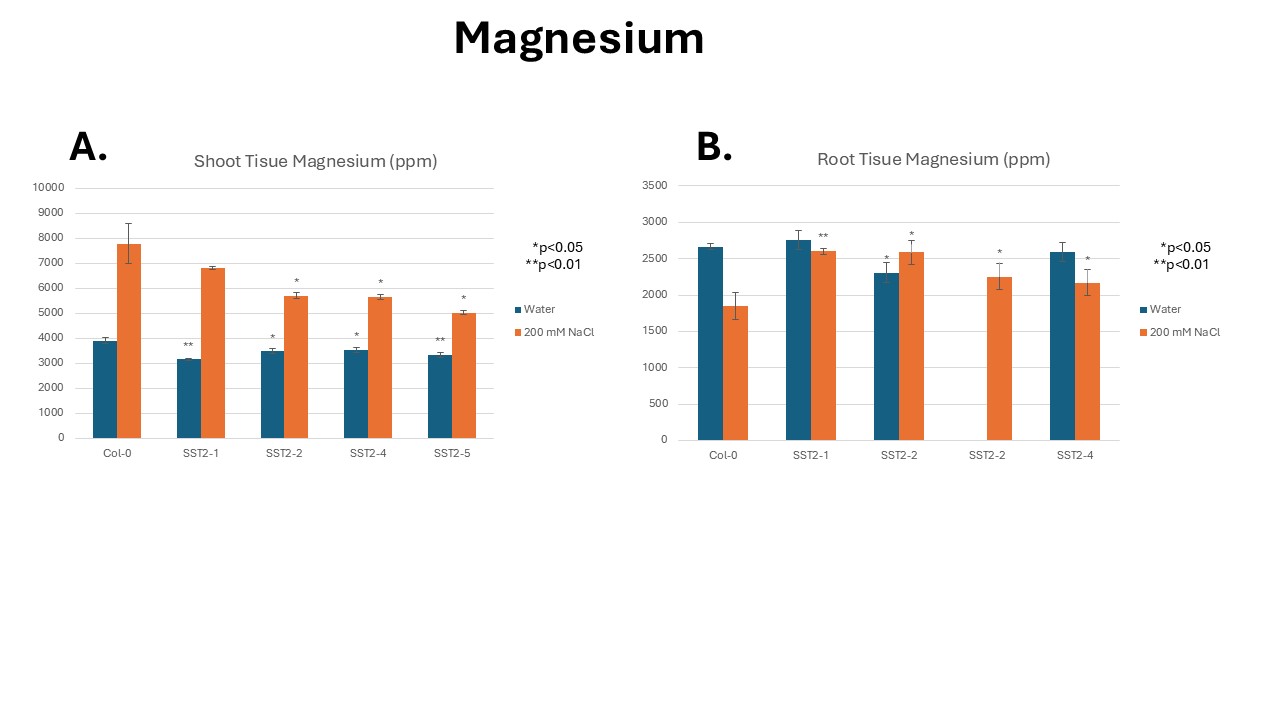

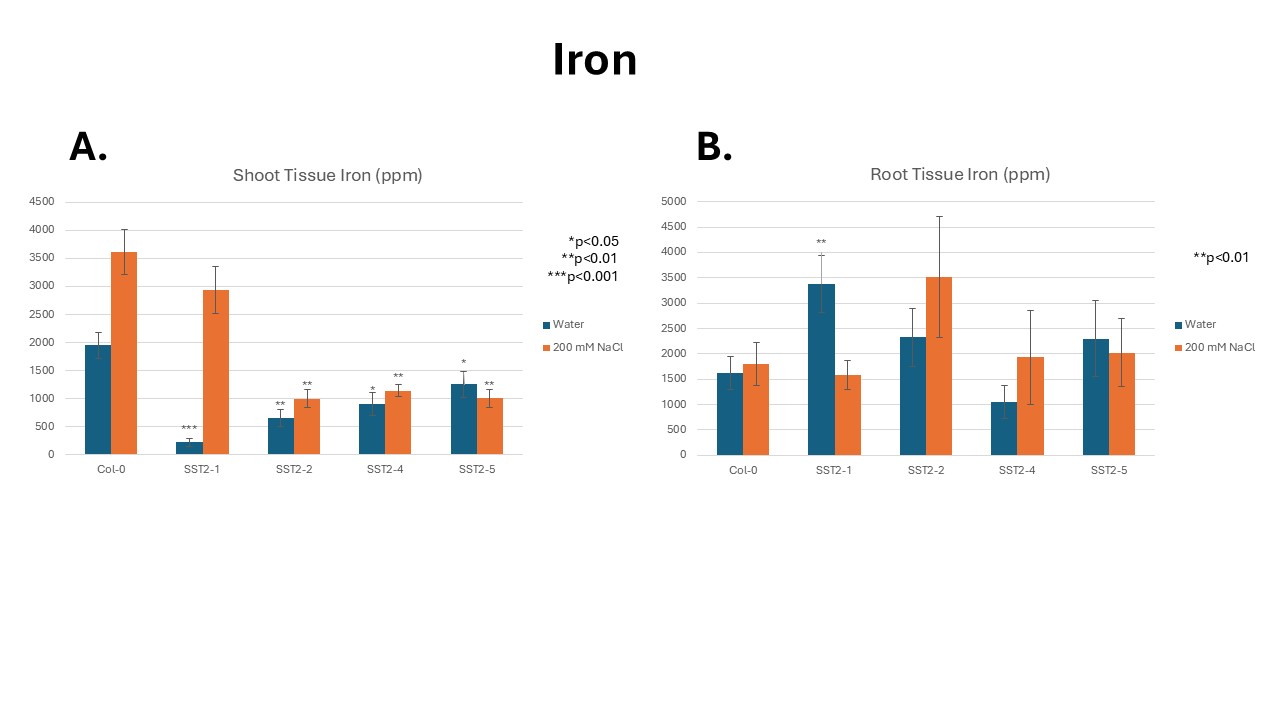

**Supplementary Fig. 6**. **ICP analysis of the accumulation of different elements including iron, magnesium, calcium and potassium.** (**A**) Shoots, water control and plants grown with 200 mM NaCl. (**B**) Roots, water and control and plants grown with 200 mM NaCl.

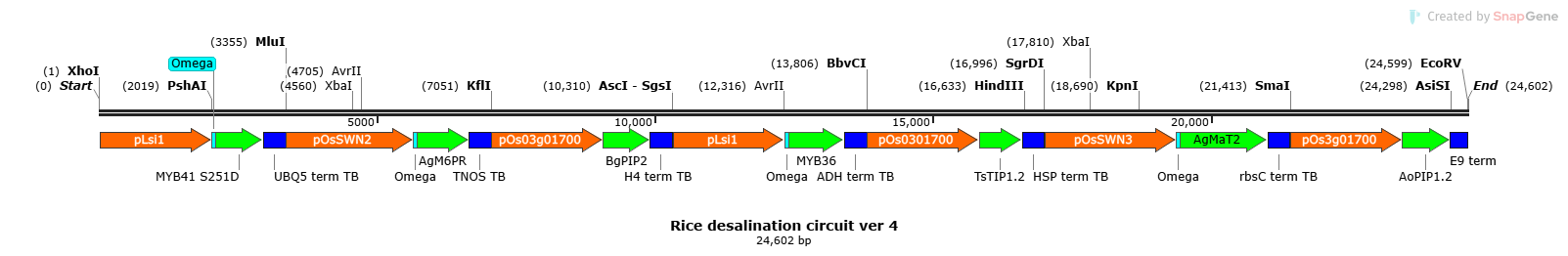

**Supplementary Fig. 7**. **Schematic of the rice salt tolerant circuit, osSST1.** Details are summarized below and found in Supplementary Data S1.

**SUPPLEMENTARY DATA DESCRIPTIONS**

**Data S1. (separate file)**

DNA sequences

**Data S2. (separate file)**

List of primers used in this study

**Data S3 (separate file)**

Complete details of preliminary suberin analysis

**Data S4 (separate file)**

Chlorophyll fluorescence kinetics data for screening of primary transgenic SST1 and SST2 lines (Excel file)

**Data S5 (separate file)**

Installation instructions and documentation for Principal Component Analysis

**Data S6 (separate file)**

Software Application for PCA Analysis

**Data S7 (separate file)**

Detailed RNA-seq data for SST plants

**Data S8 (separate file)**

Inductively Coupled Plasma Optical Emission Spectroscopy Detailed Data of ions and analyses

**MATERIALS AND METHODS**

**A. Construction of Genetic Components**

Supplementary Table 7 describes the complete components for all genetic constructs. Complete DNA sequences are found in Data S1.

**1. Testing constructs for cell/tissue specific promoters**

A pCAMBIA2300 vector containing the figwort mosaic virus (FMV) promoter, an mCherry reporter gene and the nopaline synthase terminator (TNOS) was constructed for testing individual components. The ABCG37/PDR9 and KC1 promoters were amplified from *Arabidopsis thaliana* genomic DNA prepared using the Qiagen DNeasy Plant Mini Kit. The FMV promoter from the pCAMBIA2300 test vector was excised with HindIII and BamHI and replaced with the promoters ABCG37 and KC1 for testing cell and tissue localization (Supplementary Table S1). The XCP1 and CESA7 xylem promoters were synthesized by Genewiz as part of two osmoticum gene cassettes (see section A4, below). The two gene modules were ligated into the multiple cloning site of PAB2300, a lab-modified pCAMBIA2300 vector conferring Basta resistance. Vectors for tissue localization testing of the CESA7 and XCP1 promoters were then engineered by excising the osmoticum gene component using AvrII and SacI sites, and AatII and PacI sites, respectively, and replacing with GFP mut3.1^34^. The expression constructs were transformed into *Agrobacterium tumefaciens* strain GV3101 and *Arabidopsis thaliana* Col-0 ecotype plants were transformed using the floral dip method ^35^.

**2. Testing constructs for individual synthetic epidermal barriers**

mCherry was excised from the modified pCAMBIA2300 vector described in section A1 above with BamHI and SacI. AtMYB36 and AtMYB41 were amplified from an Arabidopsis root cDNA library prepared using iScript™ cDNA Synthesis Kit (Bio-Rad, Hercules, CA) and ligated into the test vector with BamHI and SacI**.** The MYB41^S251D^ mutation was engineered using the QuikChange Lightning kit (Agilent, Santa Clara, CA). Epidermal expression of AtMYB36 was driven by the ABCG37 promoter, which was ligated into the vector using HindIII and BamHI sites. The 5’-leader sequence (omega) of tobacco mosaic virus (TMV) was included as a translational enhancer for AtMYB36. Epidermal expression of AtMYB41 was also driven by the ABCG37 promoter and ligated into the vector using the corresponding HindIII and BamHI sites. These constructs were transformed into *Agrobacterium tumefaciens* strain GV3101 and *Arabidopsis thaliana* plants were transformed using the floral dip method.

**3. Testing constructs for the synthetic transport system (aquaporins)**

Coding sequences for nine mangrove aquaporins, seven Plasma Membrane Intrinsic Proteins (PIPs)—AcPIP1, AcPIP2, AoPIP1.1, AoPIP1.2, AoPIP2, AoPIP2.2, and BgPIP2—and two Tonoplast Intrinsic Proteins (TIPs)—BgTIP and KcTIP—were codon optimized for expression in Arabidopsis and synthesized (Genewiz). The sequence information in some of the mangrove aquaporins was incomplete, and we used consensus amino acid sequences from orthologous halophyte proteins to fill the gaps (cf. DNA sequences, Supplemental Data S1). Because of limited availability of mangrove genomic information when this work was started, sequences for two TIPs—TsTIP1.1 and TsTIP1.2—from the halophyte *Thellungiella salsuginea* (salt cress)^36^ were amplified using the iScript cDNA Synthesis Kit. To create in-frame GFP- or mCherry-tagged plasmids, two aquaporin cassettes, FMV_AcPIP1_mCherry_E9T and CaMV35S_BgTIP_AtGFP3.1_Pea3AT, were synthesized with MluI and BamHI sites flanking the aquaporin genes and inserted into the pUC57 vector (Genewiz). The FMV and CaMV35S promoters in a pUC57 subcloning vector were replaced by Arabidopsis ABCG37, KC1, or SCARECROW (SCR)^37^ promoters using EcoRI/MluI or AvrII/MluI sites, respectively. The AcPIP1 gene in the mCherry plasmid, and the BgTIP gene in the GFP plasmid were replaced by the other PIPs or TIPs using flanking MluI/BamHI sites. For *in planta* functional characterization, all aquaporin cassetttes were transferred into the Basta pCAMBIA2300 vector (PAB2300) using standard restriction cloning. The second KpnI site in PAB2300 was mutated to retain the unique KpnI site in the multiple cloning site. These constructs were transformed into *Agrobacterium tumefaciens* strain GV3101 and *Arabidopsis thaliana* plants were transformed using the floral dip method.

**4. Testing constructs for Synthetic Osmolyte Genetic Modules (OGM)**

Sequences for a mannitol biosynthesis gene, Mannose 6-phosphate Reductase (M6PR)^38^, and a mannitol transporter gene, Mannitol Transporter (MaT2)^39^, both from *Apium graveolens*, were codon optimized for expression in Arabidopsis. The genes were synthesized by Genewiz in two gene cassettes comprising the AtCESA7_MaT2_H4 terminator, flanked by XbaI and PstI restriction sites, and AtXCP1_M6PR_ADH terminator, flanked by EcoRI and KpnI sites. The two gene cassettes were ligated into the multiple cloning site of PAB2300 to create the osmoticum genetic module (OGM). Our OGM is designed to work with known functions of the apoplast. Specifically, because the xylem parenchyma’s apoplast is contiguous with the water conducting xylem elements, mannitol enters the xylem elements, drawing water into the plant’s vascular tissue (Fig. 1B, main text).

A modified version of the OGM was also constructed by adding the TMV omega enhancer to the mannitol biosynthesis gene. An Ultramer DNA oligonucleotide (IDT) was used to add the TMV omega enhancer. The synthesized Mannose 6-phosphate Reductase (M6PR), including the upstream TMV omega enhancer sequence, was digested and ligated immediately downstream of AtXCP1 using AatII and PacI sites into PAB2300 containing the synthesized AtCESA7::MaT2-H4 terminator cassette to generate the complete OGM.

A second OGM, OGM2 was engineered to produce the osmolyte glycine betaine. The OGM2 circuit contains the glycine betaine synthesis genes GSMT (Glycine Sarcosine N-Methyltransferase) driven by the CESA7 promoter and SDMT (Sarcosine Dimethylglycine N-Methyltransferase) driven by the XCP1 promoter. The genes are derived from a halophilic methanoarchaeon, *Actinopolyspora halophila* and refactored for Arabidopsis expression. The circuit also contains an Arabidopsis transporter AtPro2 (AT3G55740) which was shown to have an affinity for glycine betaine, driven by the vascular related NAC-domain protein 7 (VND7) promoter.  The OGM2 circuit was added to the two-plasmid Kan and Basta circuits. To the Kan-resistant half of the circuit, we synthesized the CESA7 promoter driving expression of an omega translational enhancer and glycine sarcosine *N*-methyltransferase (GSMT) with an ADH terminator, using SbfI and BamHI sites for cloning. To the Basta-resistant half of the circuit, we synthesized the XCP1 promoter driving expression of an omega enhancer and sarcosine dimethylglycine *N*-methyltransferase (SDMT) with NOS terminator and the VND7 promoter controlling expression of the Pro2 transporter with an E9 promoter using PmeI and SmaI.

**5. Assembly of the Synthetic Epidermal Barrier (SST1)**

We assembled the complete SST from components described above that provide duplicated and expanded epidermal barriers, aquaporins, and xylem targeted osmolytes. We designed our SST circuit initially in two plasmids (two T-DNAs) because of the size and then re-assembled the SST in one plasmid (SST2).

**a. SST1 or two T-DNA version (lab number TKK716).** The complete epidermal barrier circuit was drafted using Gene Construction Kit (Textco Biosoftware, Inc., Raleigh, NC). The complete circuit was divided into two T-DNAs in the binary vectors pCAMBIA2300 (providing kanamycin selection) and PAB2300 (providing Basta selection). Circuits for each T-DNA were divided into six DNA fragments of 1.8 to 2 kb each, with 30-45 base overhangs for Gibson assembly. DNA fragments were synthesized by Synthetic Genomics, Inc. (La Jolla, CA), and were assembled into the vectors (7-fragment reaction) using the Gibson Assembly® Ultra Kit from Synthetic Genomics, Inc., following the company’s protocol. Two microliters of the Gibson assembly reaction were transformed into Endura *Escherichia coli* electrocompetent cells (Invitrogen).

The epidermal barrier circuits were initially checked for complete assembly using restriction digestion. Circuits showing the correct restriction digest patterns were sequenced using the Genewiz Sanger sequencing service. Errors in the sequence (mutations or deletions) were repaired by restriction digest excision of the DNA segment containing the error and replacement with a DNA segment with the correct sequence. The two error-free plasmids were named TKK710 (Basta circuit) and TKK711 (Kanamycin circuit). Collectively in this manuscript, plants with both the Kan and Basta circuits are TKK716. Arabidopsis plants were transformed using the standard floral dip method.

**b. SST2 or one T-DNA version (lab number TKK717).** We assembled the complete Synthetic Salt Tolerance (SST) in one plasmid using the pGreenII backbone ^40^. The pGreenII 0229 vector (Basta) was opened using the BamHI/SacI restriction sites. For TKK711, BamHI/SmaI sites were used and for TKK710, SmaI/SacI sites were used. Three-way ligation was done using T4-DNA ligase (New England Biolabs) following the company’s protocol. The epidermal barrier circuit was initially checked for complete assembly using restriction digestion with enzymes PvuII and EcoRI. Circuits showing the correct restriction digest patterns were sequence verified using the Genewiz Sanger sequencing service, with primers designed to cover the junctions between the two inserts, including upstream regions of the ABCG37 driving MYB41^S251D^ that originated on the kanamycin resistance T-DNA circuit (TKK711). The assembled vector (TKK717) and the pSOUP helper plasmid were co-transformed into *Agrobacterium tumefaciens* (GV3101). Arabidopsis plants were transformed using the standard floral dip method.

**6. Construct providing marker for suberin biosynthesis**

To follow suberin production in transgenic plants, we assembled a marker for suberin biosynthesis ^41^. The GPAT5 promoter was PCR amplified from Arabidopsis genomic DNA, adding HindIII and BamHI restriction sites, and ligated into PAB2300. mCherry-Syp22-TNOS flanked by BamHI and SacI sites was synthesized by Genewiz and ligated downstream to the GPAT5 promoter. We then co-transformed this vector with the ABCG37::MYB41^S251D^ vector (section A2) in Arabidopsis plants using the standard floral dip method.

**7. Testing plants with the synthetic root epidermal barriers (SRB) with and without aquaporins (AQP)**

We assembled two distinct plasmids containing gene circuits with the filter-barrier genes. SRB alone (epidermal suberin, epidermal Casparian strip) and one with the filter-barrier genes and the aquaporins, synthetic transport, AQP. Using the Bsp119I and SacI restriction sites, we excised all genes except ABCG37::omega_MYB36_ADH from TKK710 (section A5a), then blunted and re-ligated the digestion product. This plasmid was then co-transformed with the plasmid containing ABCG3::MYB41^S251D^ (section A2) to produce transgenic lines (SRB alone, lab name TKK733) with the suberin and Casparian strip gene components of the filter-barrier. Plants in Figure 3, center. Plants homozygous for each plasmid were obtain and used for analysis. To produce plants the filter-barriers (SRB) and the aquaporins (SRB and AQP), we digested TKK710 (section A5a) with KpnI and SacI to remove the gene CESA7::MaT2, then blunted and re-ligated the digestion product to produce a plasmid with ABCG37::omega_MYB36 and KC1::TsTIP1.2. We next digested TKK711 (section A5a) with SgsI and AfeI to remove the gene XCP1::M6PR, then blunted and re-ligated the digestion product to produce a plasmid with ABCG37::MYB41^S251D^, KC1::AoPIP1.2, and KC1::BgPIP2. These two plasmids were co-transformed to produce transgenic lines (SRB and AQP, lab name TKK734) with gene components encoding both the filter-barrier (SRB)and the three aquaporin genes (AQP) of the synthetic transport system. Plants in Figure 3, left, (SRB & AQP). After 2 years of repeatedly selecting and analyzing progeny, all plants remained heterozygous for each plasmid (Kan^R^ and Basta^R^).

**B. RT-PCR analysis of the Synthetic Epidermal Barrier circuit transcripts**

Root tissue from independent SST circuit transgenic lines was harvested and quickly froze in liquid nitrogen. Total RNA from the root tissue was prepared using the Qiagen RNeasy Plant Mini Kit. Potential contaminating genomic DNA was removed from the RNA samples using the Invitrogen TURBO DNA-free Kit. cDNA was generated from 100 ng of total RNA using the Bio-Rad iScript™ cDNA Synthesis Kit. Two microliters of the cDNA reaction were added to a 50 μl RT-PCR reaction. PCR amplification was performed using Promega GoTaq® Green Master Mix and gene-specific primers. Thermocycler conditions were as follows: initial 2-min denaturation at 95°C, 30-sec denaturation at 95°C, 30-sec annealing at 55°C, 30-sec elongation at 72°C, 30 cycles. Fifteen microliters of the reaction were resolved by electrophoresis on a 2% agarose gel and visualized with ethidium bromide staining using a Bio-Rad ChemiDoc imaging system.

### **C. Clearing, staining and microscopic imaging of plants**

Five-day-old roots were fixed in paraformaldehyde, then cleared in ClearSee as per Kurihara et al. ^42^. Basic Fuchsin staining was carried out as described in^43^, and fluorol yellow staining with aniline blue counterstaining was carried out as described in^41^. Berberine-aniline blue staining was performed as per Brundrett et al.^44^. Plants in Figure S9 were grown on MS plates with 1 μM 2-aminoethoxyvinyl glycine (AVG), to inhibit ethylene biosynthesis that might break down suberin, before sectioning and staining^41^. Plants in Figure 2 were grown on MS plates without AVG. The following are the excitation (ex.) and the detection/emission (em.) parameters: GFP, ex. 488 nm, em. 530-560 nm; Basic Fuchsin/mCherry, ex. 543 nm, em. 610 nm; Fluorol Yellow 88, ex. 488 nm, ex. 530 nm; Berberine-Aniline Blue, ex. UV, em. 365 nm. Root tips were imaged using a 10X objective, and cross sections were imaged using a 20X or 40X objective. Free-hand cross sections were generated by hand slicing using double-edged razor blades ^45^. Hand-sectioned or intact stained roots of ABC::MYB41^S251D^/ GPAT5::AtmCherry-Syp122 were observed under either a LEICA CTR DM5500 microscope (hand sections) or a Zeiss LSM 510-Meta confocal microscope. KC1::AoPIP1.2_mCherry, KC1::BgPIP2_mCherry, pKC1::TsTIP1.2_GFP, ABCG37::MYB36, and all other GFP/mCherry images (from intact roots or hand-sectioned roots) were imaged on a Leica DM5000B using TRITC and FITC filter cubes.

Berberine-aniline blue stained roots, expressing ABCG37::MYB36, were hand sectioned, mounted on 50% glycerol, and subsequently covered with a cover glass (VWR International) for imaging. Confocal micrographs with fluorescence from berberine hemisulfate (lignin stain) signal were acquired with IX81 Inverted Spinning Disk Confocal Microscope (Olympus) using 40x objective under the GFP setting (ex. 488 nm, em. 530-560 nm).

**D. Plant Materials, Growth Conditions, and Methods for Evaluation of Transgenic Lines**

**1. Standard growth of Arabidopsis plants**

Seeds of wildtype (*Arabidopsis thaliana*, Columbia-0) were surface sterilized and germinated on Murashige and Skoog (MS) media full-strength MS salts, 0.8% phyto agar, 3% sucrose, pH 5.7, and grown for six days at 22°C under short days (10h of light) or long days (16h of light). Typically, after two weeks in short day conditions or 7-10 days in long day conditions, plants were transferred to soil and growth continued in plant growth chambers (Percival) under short day or long day conditions. Plants for seed production were transferred to light racks and grown at 22°C under a long day cycle (16h light/8h dark cycle). For plant transformation, short day grown plants were transferred to a long day growth chamber set at 24°C with typical light conditions (50-100 E) to induce flowering.

**2. Testing Synthetic Symplastic Transport System: mangrove and halophyte aquaporins function in Arabidopsis**

Overexpression of aquaporins in roots has been shown to improve salt and drought tolerance in plants ^46-48^. Hence, mangrove and halophyte aquaporins that we determined to be stable in Arabidopsis roots (Supplementary Fig. 2 and Supplementary table 2) were tested for function using salt and drought tolerance assays.

**a. Drought tolerance assay on transgenic plants expressing individual aquaporin genes.**

We performed functional assays on transgenic lines expressing four PIPs to determine whether they could confer drought tolerance to plants. Based on previous studies ^46-48^, if the aquaporins are functional in Arabidopsis, we expect more efficient water transport and hence drought tolerance in transgenic plants. We separately expressed in plants the aquaporin genes AoPIP1.1, AoPIP1.2, AoPIP2.2, and BgPIP2 under control of the SCR or ABCG37 root epidermal promoter. Primary transgenic plants were selected and second generation (T1) seed obtained. Independent transgenic lines were plated on selectable media and transgenic plants transferred to soil in duplicate 4-inch pots which were placed in a growth rack with cool white lights under long-day conditions, providing humidity less than 20%. One week after transplanting to soil, water was withheld from one set of plants (transgenic and Col-0) for 10 to 11 days while the second set (transgenic and wildtype) was watered daily. These drought tolerant transgenic lines confirmed aquaporin function in Arabidopsis (Supplementary table 2, Supplementary Fig. 3 & 4).

**b. Salt tolerance assay on transgenic plants expressing individual aquaporin genes**

We followed a previously developed method where plants with 50 - 200 mM salt tolerance continue to grow, bolt, flower and set seed in the presence of salt^49^. We separately expressed in plants the aquaporin genes TsTIP1.1 and TsTIP1.2 under control of the SCR or ABCG37 root epidermal promoter. Primary transgenic plants were selected and second generation (T1) seed obtained. Transgenic plants were grown as described in section D2a. To test salt tolerance, one week after transplanting in soil, we watered one set of plants (transgenic and wildtype) with 350 mM NaCl (~58% seawater, without salt step up procedure) every day for 12 days and another set of plants with tap water. On the 12^th^ day, all plants were photographed and assayed for continued growth, bolting and seed set, reported indicators of salt tolerance^49^ (Supplementary table 2, Supplementary Fig. 5).

**3. Screening Primary SST Transgenic Lines**

**a. Arabidopsis plant growth and Salt Treatment for Selection of Primary SST Engineered Lines**

Figures below provide schematics for plant growth, selection and salt treatments. For the experiments described herein, we kept the initial plant treatments (Supplementary Fig. 8A) and salt step uniform so that we could access the best primary transgenic lines (Supplementary Fig. 8B), and the ability of transgenic lines to survive continuous salt treatment (Supplementary Fig. 8C).

Transgenic Arabidopsis seeds were surface sterilized, stratified at 4°C and germinated on MS selection plates. Transgenic plants were selected on plates with kanamycin (50 μg/ml) or Basta (25 μg/ml). For selection of primary transgenics, in addition to kanamycin or Basta, MS plates included carbenicillin (50 μg/ml) or cefotaxime (100 μg/ml) to inhibit growth of Agrobacterium. Columbia-0 (Col-0), wildtype seed were treated as described for transgenic plants except the germination media was simply MS. Propagation of transgenic seed were after plating on a 15 cm petri dish with MS media and appropriate selectable markers.

After seed were plated, seedlings were then grown on plates for 18 days in a growth chamber set at 22°C under short days (10-h light/14-h dark cycle) before transfer to soil in pots (details, Supplementary Fig. 8A). Wild type and transgenic plants were allowed to acclimatize in soil in 2-inch pots for 10 days (details, Supplementary Fig. 8A-B) in a growth chamber under short day (10-h light at 22°C /14-h dark at 18°C) and high humidity (70-72%) conditions at an altitude in Colorado of approximately 5000 feet (1524 meters).

**Salt step-up and treatment methods (Supplementary Fig. 8B-C).** After 18 days on plates and 10 days of acclimation to soil, plants were then sub-irrigated (flooded up to the rim of the 2-inch pot) with a NaCl solution each day. We increased the salt concentration every two days (i.e., 2 days at 100 mM NaCl, 2 days at 200 mM NaCl, 2 days at 300 mM NaCl, 2 days at 400 mM NaCl and 2 days at 600 mM NaCl, Supplementary Fig. 8B). Fresh solutions of NaCl were used to water plants each day with pots remaining in salt water for 1 hour. After the step-up with two days of salt at 600 mM NaCl, plants were transferred to a low humidity environment to increase transpiration (typically 10-12%, less than 20% relative humidity) in growth racks with cool white lights. Plants were maintained at the same salt treatment, 600 mM, for 6-8 days to select the best primary transgenic lines (Supplementary Fig. 8B). After these additional days at low humidity, a strong visual difference was noted (Fig. 4), plants were photographed and chlorophyll fluorescence kinetics were measured (section D3b). Out of initial concern to obtain seed from these lines for further study, we flushed salt from the soil and recovered primary transgenic seed from plants treated with fresh water (tap water). In one exception (Fig. 4C), plants starting the salt step-up procedure were three-week-old plants, rather than 28-day-old plants.

**Supplementary Fig. 8A.** Plant growth and salt treatment conditions for plants in this study (**A**) Starting seeds and growth on plates
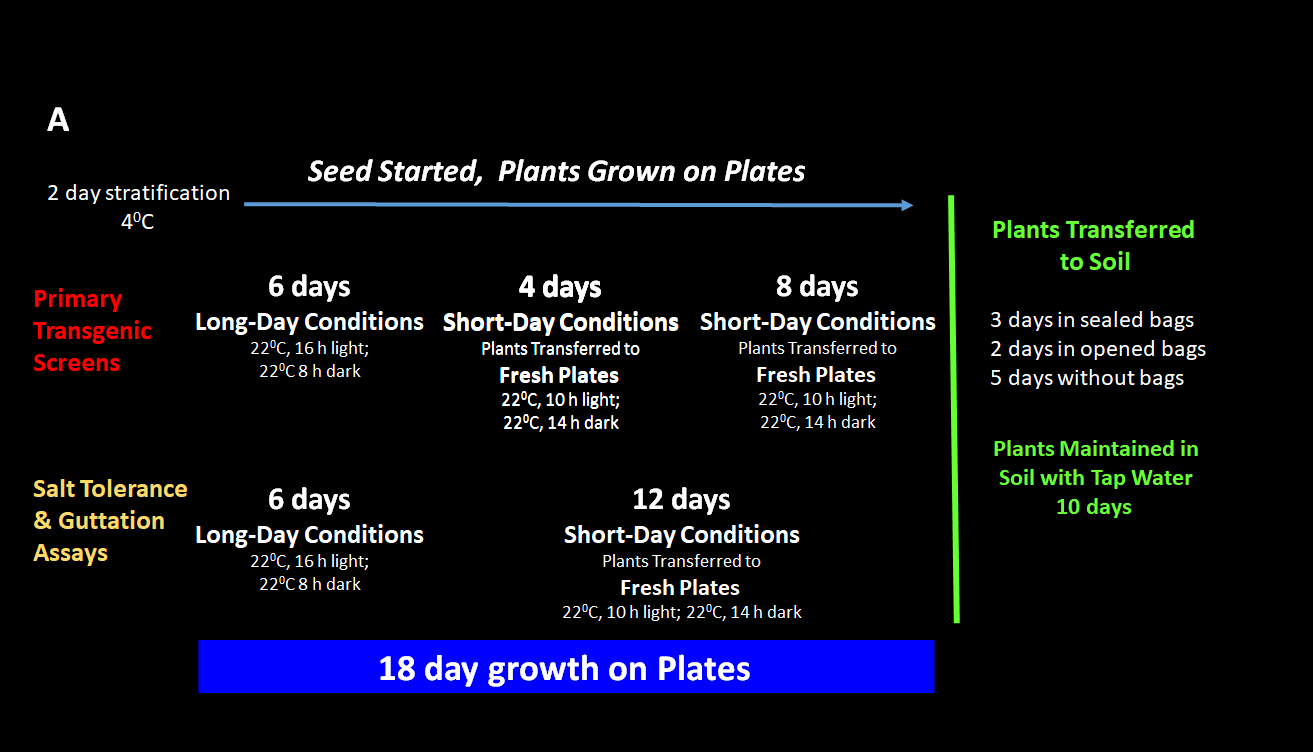

**Supplementary Fig. 8B.** Plant growth and salt treatment conditions for plants in this study. (**B**) Salt treatment for evaluation of salt tolerance and selection of primary transgenic lines

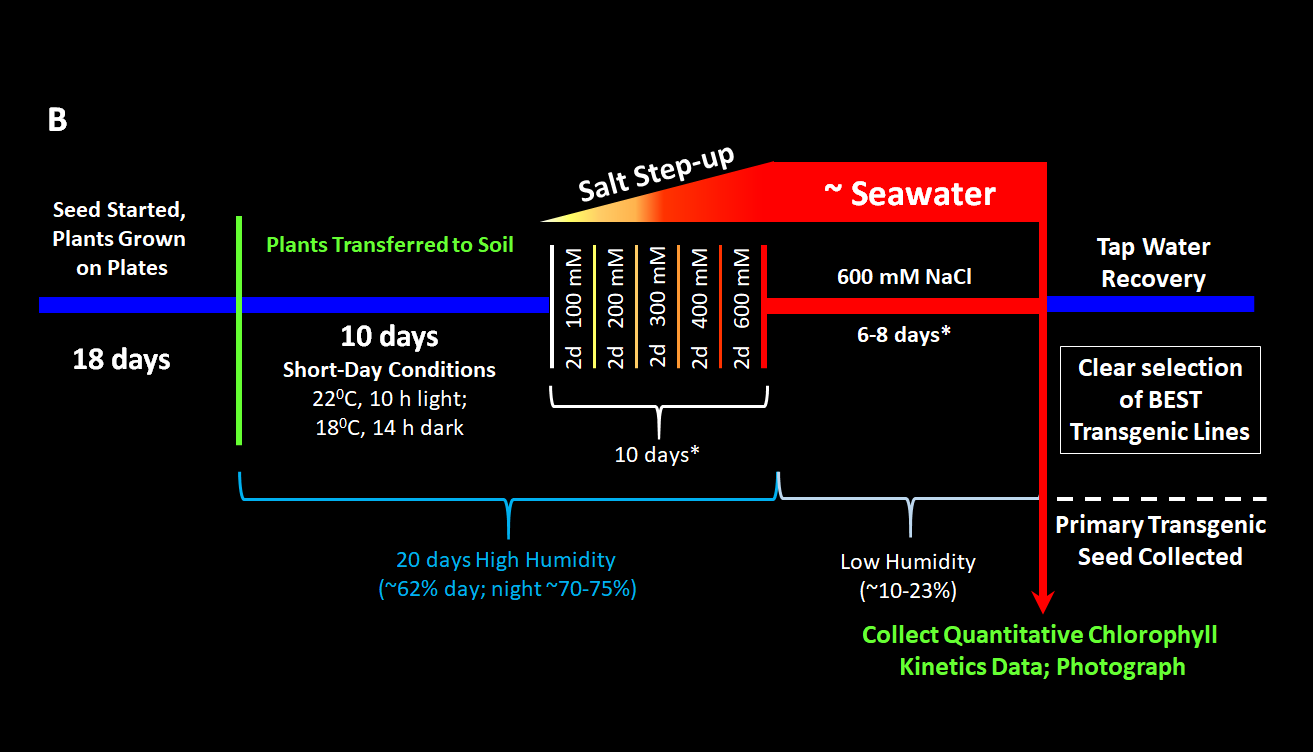

**Supplementary Fig. 8C.** Plant growth and salt treatment conditions for plants in this study. (**C**) Continuous salt treatment

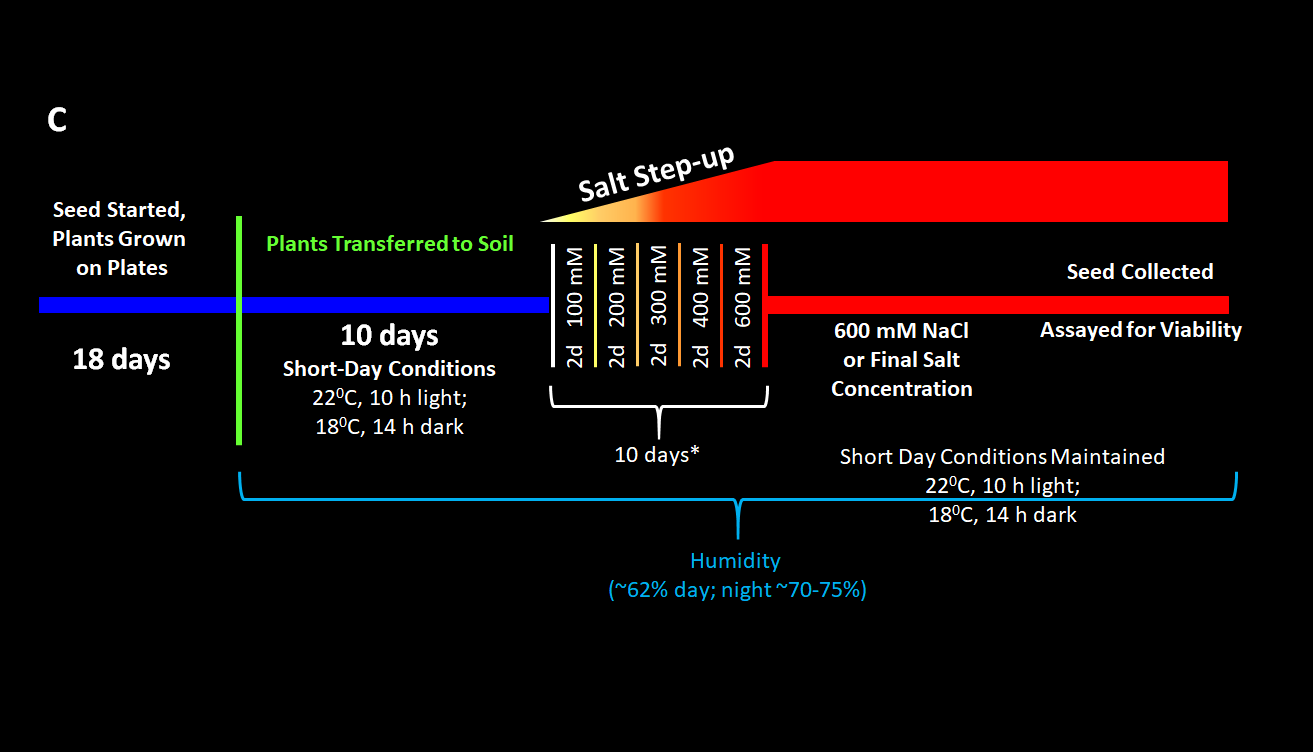

**b. Quantitative screening Parameters: chlorophyll fluorescence kinetics**

Measurements of chlorophyll fluorescence kinetics (Supplemental Data S3) were one criterion used to screen and evaluate transgenic plants containing our Synthetic Epidermal Barriers for their response to salt. Engineered plants with the SST gene circuits were screened for variation in chlorophyll fluorescence kinetics and the data were analyzed using Principal Component Analysis (PCA). Transgenic plants were treated with salt as described in section D3a (salt treatment protocol) and were compared with wildtype Arabidopsis (Col-0) plants that were controls (non‐salt treated) and salt-treated. Whole-plant chlorophyll fluorescence images were obtained using a FluorCam FC800-O instrument (Photon Systems Instruments, Drasov, Czech Republic). Plants were dark‐adapted for 30 min prior to fluorescence measurements. Data were obtained using the default fluorescence Quenching Analysis protocol in the manufacturer's software (FluorCam version 7). The software produces 71 fluorescence parameters per plant, 32 measured and 39 calculated, plus image size in pixels.

**c. Principle component analysis of chlorophyll fluorescence kinetics**

We developed an application in Matlab v. 9.4.0.813654 (R2018a) (The MathWorks, Inc., Natick, MA) to analyze the fluorescence parameters using PCA, thereby reducing 71 dimensions to two, and plot the data on the first two principal axes (Supplemental Data S4 and S5). The application utilizes Quadratic Discriminant Analysis (QDA) (Figure 4) to establish geometric regions of classification based upon the distributions of freshwater or salt-water treated wildtype controls in two-dimensional principal component space. The salt-treated transgenic plants are then classified as performing similarly to one of the two control groups, based upon their location with respect to these regions on the scatterplot. Additionally, a linear discriminant analysis (LDA) is applied to the parameters, using the controls to train the classification algorithm. The linear projection is displayed graphically and a table ranks the plants in terms of their score. PCA, QDA, and LDA can be performed on the complete set of 71 parameters, the measured parameters, or the calculated parameters.

**d. Rice transformation**

Rice transformation followed published protocols ^50^ with minor modifications from the laboratory of Professor Jan Leach at Colorado State University.

**e. Screening and Evaluation of salt tolerance – Rice.** All seeds were germinated by placing 20-30 rice seeds in large plant plates lined with filter papers soaked in 15mL of 1:1000 MAXIM XL fungicide. Seeds were germinated in a continous light at 30 ^O^C growth chamber for 5-7 days, until cotyledons form and reach about 1cm in length, then transferred to soil in bags for 3 days and allowed to grow in soil for an additional 7 days. Subsequently, plants were transferred to large 62-liter tanks (100.3 cm x 51.4 cm x 17.8 cm) containing either water or salt solutions. All salt treatments were carried out under greenhouse conditions with supplemental lighting. Rice seedlings were exposed to Initial salt concentration of 100 mM NaCl with the salt level increasing by 100 mM NaCl every 2 days until it reached 600 mM NaCl (10 days in salt step up) and then maintain in 600 mM NaCl for an additional 7-10 days. See Supplementary Fig. 9 for the detail work flow. Hence rice plants were in saline conditions for 17-20 days. The plants were photographed for publication.

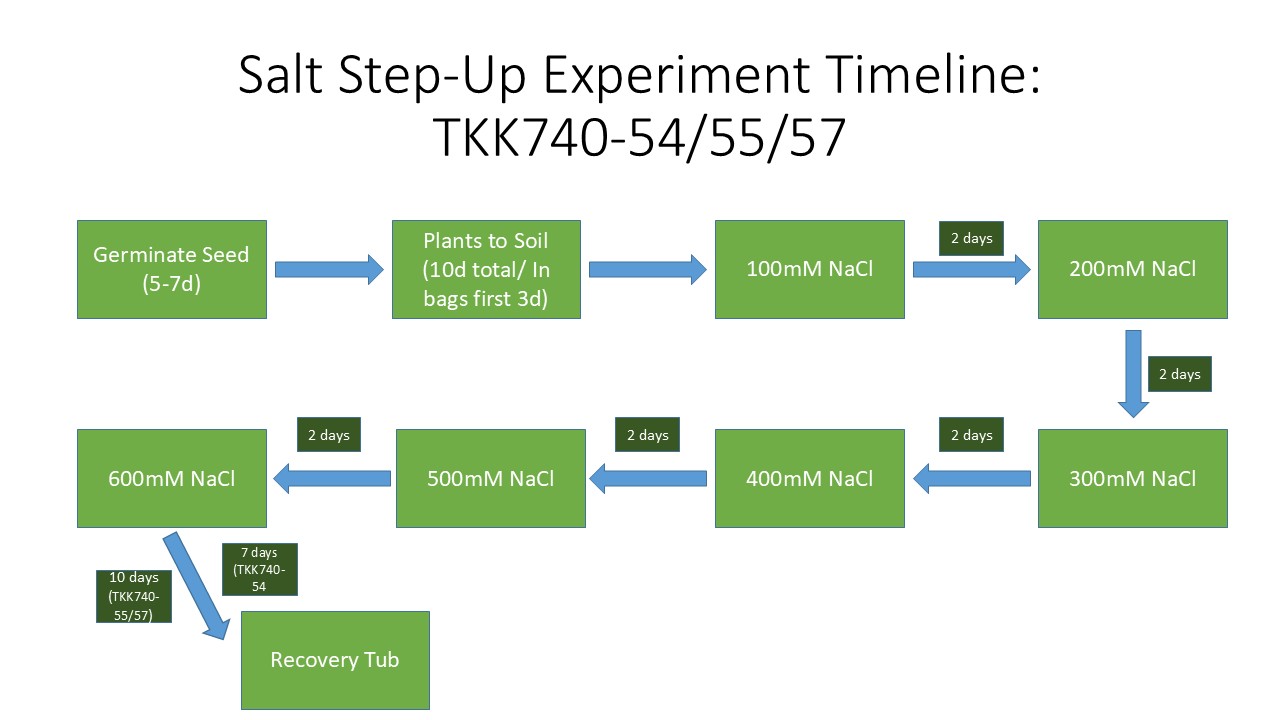
 **Supplementary Fig. 9.** Schematic of salt step-up conditions for rice engineered with our OsSST gene circuit.

**4. Plant growth and Conditions for Salt Tolerance Evaluation**

**a. Evaluation of Salt tolerance - Arabidopsis**

Please see Supplementary Fig. 8 for schematics summarizing details here. Wild type and transgenic seeds were surface sterilized, incubated in dark at 4°C for two days for stratification and germinated on MS media, with selection for transgenic lines, and grown for six days at 22°C under long-day cycle (16h light, 8h dark) (Supplementary Fig. 8A). Seedlings were transferred to new plates and grown in a short-day growth chamber (10-h light /14-h dark at 22°C) for an additional twelve days before being transferred to soil in 2-inch pots. Plants were allowed to acclimatize for 10 days in a growth chamber under short day (10h light at 22°C /14h dark at 18°C), and high humidity conditions (70-75%) (Supplementary Fig. 8A-B). Plants were salt treated as described for screening primary transgenic lines by sub-irrigation with increasing concentrations of freshly prepared NaCl solutions (Supplementary Fig. 8B) until the desired final concentration was reached and were moved to a light rack at low humidity (Supplementary Fig. 8B) for six to eight additional days at the final salt concentration. Quantitative data about the salt response were collected with a FluorCam for chlorophyll fluorescence kinetics and the plants were photographed.

**b. Continuous salt exposure - Arabidopsis**

For continuous salt treatment assays (Extended Data Fig. 8, Extended Data Table 1), seed were grown on plates and soil as previously described (Supplementary Fig. 8) and soil grown plants were exposed to the same salt step-up procedure (Supplementary Fig. 8C). After plants reached their final salt concentration (400mM, 500mM, or 600mM) they were maintained at that concentration (watered daily with sub-irrigation) until seed production under short day (10h light at 22°C /14h dark at 18C) and high humidity (70-75%) conditions and siliques harvested and seed evaluated (Extended Data Fig. 8, Extended Data Table 1).

**5. Suberin Analysis via GC-MS**

Just describe the “chloroform dip” for the waxes. Looking at the references, they used different sets of plants for the various extractions, e.g., waxes, lipids, and polymeric lipids. Conversely, we used the same set of plants/roots for all three extractions (not the best approach probably). So, we will just present the “waxes” in the main text and have the remaining (lipids and polymeric lipids) in the supplement. Regarding the actual GC-MS protocol, oven temp, ramping, etc., we will just reference the chemistry core facility so we can forego the gritty details.

**6. Ion analysis of SST Arabidopsis plants**

Arabidopsis SST plants were grown in 4-inch pots containing pre-washed Greens Grade Porous Ceramic Topdressing (Profile, Buffalo Grove, IL). Plants were sub-irrigated twice weekly with water containing basic nutrients for four weeks in short day conditions (8 hours of light) 22C/18C. After four weeks salt treated plants were stepped up in 50 mM NaCl increments every two days to a final concentration of 200mM. After one week at maximum salt concentration, root and shoot tissue was harvested for further analysis. Plant samples were dried and ground to homogenize them. 100mg of shoot tissue was weighed and put in 1.5ml tubes. 50mg of root tissue was weighed and put in 1.5ml tubes. The dried tissue was then moved to 22mm x 200mm glass acid digestion tubes. 1ml of ultra-trace grade 70% nitric acid (HNO3) was then added to each glass tube. The samples were then digested for 2 hours at 60°C and 6 hours at 130°C on a digestion block. After the samples cooled, they were diluted to 10ml with distilled water in 15ml tubes. The samples were tested for elemental analysis using Inductively Coupled Plasma- Optical Emission Spectroscopy (ICP-OES) model PerkinElmer 7300 DV. Appropriate standards and controls were followed for each sample. The data was collected as mg/L for each element and was then converted to ppm by multiplying by 10,000 and dividing by the biomass dry weight for roots or shoots.

**7. RNA-seq analysis of SST2 plants**

Plants for RNA-seq analysis were germinated and grown vertically for two weeks on large MS plates (145x20mm) with selection (BASTA). After two weeks they were transferred to large plates containing 200mM NaCl. After one week on 200mM NaCl, roots were separated and rapidly frozen in liquid nitrogen for RNA extraction. RNA was extracted using 100mg of root tissue per replicate and TRIzol Reagent (Thermo Fisher) according to the manufacturer's protocol. RIN values for all RNA samples were calculated using an RNA TapeStation system (Agilent). Three technical replicates for the following conditions, Col-0, Col-0 grown with 200mM NaCl, SST2-1, and SST2-1 grown with 200mM NaCl in the media. Sequence analysis used NOVAseq paired end (2x150) sequencing as described at the University of Colorado Anschutz campus (https://www.cuanschutz.edu/).

**8. Identification of commonly regulated salt response genes**

The differentially expressed genes (DEGs) identified from the pairwise comparisons of Col‑0 salt‑treated, (Synthetic salt tolerant) SST control, and SST salt‑treated samples relative to Col‑0 controls were subsequently analyzed using the online Interactive Venn tool (52) to identify genes associated with salt‑responsive expression that where similarly regulated with the salt treated groups as well as in SST control plants that had not experience salt exposure.

Gene annotation information used in the differentially expressed gene tables was obtained through bulk data retrieval from The Arabidopsis Information Resource website (53)
